# Metabolic and Proteomic Defects in Human Hypertrophic Cardiomyopathy

**DOI:** 10.1101/2021.08.18.455967

**Authors:** Michael J. Previs, Thomas S. O’Leary, Neil B. Wood, Michael P. Morley, Brad Palmer, Martin LeWinter, Jaime Yob, Francis D. Pagani, Christopher Petucci, Min-Soo Kim, Kenneth B. Margulies, Zoltan Arany, Daniel P. Kelly, Sharlene M. Day

## Abstract

**Rationale:** Impaired cardiac energetics in hypertrophic cardiomyopathy (HCM) is thought to result from increased ATP utilization at the sarcomere and is believed to be central to pathophysiology. However, the precise defects in cardiac metabolism and substrate availability in human HCM have not been defined.

**Objective:** The purpose of this study is to define major disease pathways and determine the pool sizes of intermediary metabolites in human HCM.

**Methods and Results:** We conducted paired proteomic and metabolomic analyses of septal myectomy samples from patients with HCM and compared results to non-failing control human hearts. Increased abundance of extracellular matrix and intermediate filament / Z-disc proteins, and decreased abundance of proteins involved in fatty acid oxidation and cardiac energetics was evident in HCM compared to controls. Acyl carnitines, byproducts of fatty acid oxidation, were markedly depleted in HCM samples. Conversely, the ketone body 3-hydroxybutyrate, lactate, and the 3 branched chain amino acids, were all significantly increased in HCM hearts, suggesting that they may serve as alternate fuel sources for the production of ATP. ATP, nicotinamide adenine dinucleotide (NADH), NADP and NADPH, and acetyl CoA were also severely depleted in HCM hearts. Based on measurements from human skinned muscle fibers, the magnitude of observed reduction in ATP content in the HCM hearts would be expected to decrease the rate of cross-bridge detachment, implying a direct effect of energy depletion on myofilament function that could contribute to diastolic dysfunction.

**Conclusions:** HCM hearts display profound deficits in cardiac energetics, marked by depletion of fatty acid derivatives and compensatory increases in other metabolites that could serve as alternate fuel sources. These results lend support to the paradigm that energy depletion contributes to the pathophysiology of HCM and also have important therapeutic implications for the future design of metabolic modulators to treat HCM.

---

Hypertrophic cardiomyopathy (HCM) affects 1 in 500 people, making it the most common genetic cardiovascular condition of Mendelian inheritance.^1^ Patients with HCM experience a progressive disease course and many suffer serious adverse events including heart failure, arrhythmias, and early mortality.^2^ The sarcomere is the primary locus of HCM variants, with 40-50% of patients with HCM carrying pathogenic variants in one of eight genes coding for sarcomere proteins.^3^ A limited number of non-sarcomere genes with an established strong association with HCM collectively make up a small fraction of cases,^4, 5^ with the remainder being classified as “genotype negative” HCM. A large proportion of this latter category are increasingly being recognized as polygenic in nature with modifying environmental factors.^6^ Despite the diverse monogenetic and polygenic causes of HCM, there is considerable overlap in terms of the phenotypic manifestations and clinical outcomes.^2^

Over the past three decades, there has been an enormous effort by basic and translational scientists to understand the molecular events that lead to HCM. The vast majority of these studies have used genetic cellular and animal models with a pathogenic variant in one of the eight sarcomere genes expressed. While these preclinical studies have yielded highly valuable insights into disease pathogenesis, it has been difficult to tease apart the primary effects of the gene modification from secondary events that occur as result of cardiac remodeling. A number of studies using human tissue from patients with HCM, with symptomatic left ventricular outflow tract obstruction who have undergone septal myectomy, have emerged over the past several years. These studies have identified a variety of defects in human HCM in comparison to control hearts, including increased myofilament calcium sensitivity,^7^ increased actomyosin ATPase activity,^8^ calcium dyshomeostasis,^9^ protein quality control abnormalities,^10^ and multiple differences in protein abundance.^11–13^ While some disease-related aberrations are more pronounced, or in some cases specific to sarcomeric HCM, many appear to be genotype-independent and thus may represent shared downstream pathways related to secondary cardiac remodeling.

Impaired cardiac energetics is one such shared downstream pathway that is believed to be central to HCM pathophysiology.^14, 15^ Energetic defects have been long been recognized in genetic mouse models of HCM,^16, 17^ as well as in patients,^18, 19^ with significant reductions in energy reserve reflected by a decreased ratio of phosphocreatine (PCr) to ATP. In patients with HCM, a lower PCr:ATP ratio is associated with progression of cardiac fibrosis as measured by cardiac magnetic resonance imaging.^20^ More recently, dysregulation of specific enzymes and proteins involved in cardiac metabolism has been described in HCM.^11, 13^ In particular, reduced abundances of enzymes involved in fatty acid oxidation in HCM has suggested a metabolic “shift” away from fatty acids as the primary fuel source of the normal heart. A similar shift in substrate utilization has been demonstrated in experimental models of pathologic cardiac hypertrophy.^21^ In the healthy heart, β-oxidation of fatty acids produces the majority of acetyl-CoA that feeds into the TCA cycle to generate ATP.^22^ In conditions of chronic myocardial stress that lead to heart failure, metabolic reprogramming of myocardial fuel utilization from free fatty acids to glycolysis occurs likely as an initially adaptive mechanism to reduce the consumption of oxygen. But over time, it is believed that this shift results in insufficient ATP production to meet the energetic demands of the heart, thus contributing to the pathogenesis of myocardial dysfunction. ^22, 23^ The impact of this apparent metabolic re-programming on the pool sizes of intermediary metabolites has not been previously measured in hearts from patients with HCM. Further, while metabolic defects have been postulated to lead to energy depletion, it has not been clear whether the reduction in adenosine triphosphate (ATP) availability is sufficient to impair cross bridge dynamics, thus contributing in a causal way to disease pathophysiology.

With these knowledge gaps in mind, we conducted paired proteomic and metabolomic analyses of heart samples obtained from patients with HCM with defined genotypes undergoing septal myectomy, and compared the results with samples from the septum of non-failing control human hearts. The proteomic analyses were consistent with major findings previously reported by other groups, including upregulation of extracellular matrix proteins and intermediate filament / Z-disc proteins, and downregulation of proteins involved in fatty acid oxidation and energy stores in all HCM groups compared to control samples.^11, 13^ By a quantitative targeted metabolomics analysis we observed uniform and marked depletion of fatty acid derivatives in all HCM groups (independent of genotype) compared to control hearts, particularly long chain acyl carnitines. Conversely, the ketone body 3-hydroxybutyrate, lactate, and the 3 branched chain amino acids, were all significantly increased in HCM hearts, suggesting that they may serve as alternate fuel sources to fatty acids for the production of ATP. ATP and other trinucleotides, nicotinamide adenine dinucleotide (NADH) and its phosphate derivatives, NADP and NADPH, were also severely depleted in HCM hearts. Finally, using a human skinned muscle fiber assay, we demonstrate that the magnitude of observed reduction in ATP content in the HCM hearts would be expected to decrease the rate of cross-bridge detachment, implying a direct effect of energy depletion on myofilament function that could contribute to diastolic dysfunction.

## Methods

### Patient sample acquisition

Myocardial tissue was obtained from the interventricular septum of unrelated subjects with HCM at the time of surgical myectomy for symptomatic left ventricular outflow tract obstruction at the University of Michigan as described previously.^9^ This study had the approval of the University of Michigan Institutional Review Board, and subjects gave informed consent. See expanded methods and materials in the online supplemental materials.

For the functional assays, epicardial biopsies were obtained from patients undergoing coronary artery bypass grafting at the University of Vermont Medical Center. A ~10-30 mg piece of muscle was excised from the left ventricular free wall and immediately submerged in oxygenated ice-cold Krebs buffer containing 30 mM, 2,3-butanedione monoxine.^24^ These procedures were in accordance with the human subjects protocol approved by the Internal Review Board of the University of Vermont Medical Center and all patients gave informed consent.

### Sample preparation for proteomic analyses

A small portion (1-2 mg) of each frozen heart biopsy was digested to peptides and processed as described.^25^ See expanded methods and materials in the online supplemental materials.

### Liquid chromatography mass spectrometry (LCMS) for proteomic analyses

The tryptic peptides were separated by liquid chromatography and directly infused into a mass spectrometer for identification and quantification of peptide abundances as described. Data were collected in data dependent MS/MS mode with the top five most abundant ions being selected for fragmentation and record as .raw files as previously described.^25^ See expanded methods and materials in the online supplemental materials.

### Quantification of protein abundances

Protein abundances were determined from the LCMS analyses using a label-free quantification routine as described.^25^ See expanded methods and materials in the online supplemental materials.

### Statistical analysis of proteomic data

To correct for the non-linear distribution in the normalized peptide abundances caused by large differences in the intrinsic ionization efficiencies of each peptide, the values were transformed using the natural log. The natural log transformation produces a normal distribution which was then subjected to a Student’s t-tests.

Unpaired student’s t-tests were performed in Excel (Microsoft) on array of natural log abundances from all peptides for a given protein in the HCM samples versus control (donor) accounting for both intra- and inter-group variability. Unpaired student’s t-test were run on the large array of the natural log normalized abundances from all peptides in all the proteins within each subcellular compartment to determine if related proteins differed significantly between the HCM groups and the donor control (e.g. testing if mitochondrial content was significantly reduced in the HCM groups).

The pooled standard deviation of each average protein ratio was calculated using a Taylor Expansion to the standard deviation for each grouped peptide ratio. That grouped peptide ratio standard deviation was pooled into a grouped protein ratio standard deviation by accounting for the degrees of freedom within each group. The standard error of the mean (SEM) was then calculated by dividing the pooled standard deviation by the square root of the number of samples within both the HCM and control groups. The SEM reflects both inter- and intra-group variability. A Bonferroni correction was performed to account for multiple comparisons (0.05/527 = 0.000095), and a P value of < 0.0001 was considered statistically significant.

### Quantitative targeted LCMS metabolomics

About 100 mg of each frozen heart sample was lyophilized overnight, powdered, and weighed to make homogenates for each targeted LCMS metabolomics assay (acylcarnitines, amino acids, organic acids, nucleotides, and malonyl and acetyl CoA). Metabolites were extracted in cold aqueous/organic solvent mixtures according to validated, optimized protocols.^26, 27^ Please see expanded materials and methods in the online supplemental materials.

### Statistical analysis of metabolomics data

Statistical analysis and visualizations for the metabolomics data were performed using R 4.0.3. A two-tailed Student’s t-test was used for the comparison between two experimental groups. For experiments with more than 2 groups, an ANOVA was performed using the aov R function. Post-hoc testing of significant results was performed using Tukey’s HSD test. PCA analysis was performed using the FactoMineR R package and visualized using the factoextra R package. Heatmaps were creating using the Complexheatmap R package.

### Estimation of ATP concentration *in vivo*

To estimate the concentration of the adenosine triphosphate (ATP) within the ventricular tissue that was available for actomyosin cross-bridge cycling (i.e. contractility), the abundance of ATP measured per gram of lyophilized heart powder was converted to a concentration. The dry weights of the lyophilized powered were assumed to be 20% of the total mass of the samples ^28, 29^ and each gram of water was assumed to occupy 1 mL of the tissue. Only 60% of this concentration was assumed to be available for actomyosin cross-bridge cycling based on previous estimates of ATP usage in cardiac muscle.^30^

### Determination of the effect of ATP on actomyosin cross-bridge detachment in skinned human epicardial tissue

To determine the effect of ATP on actomyosin cross bridge detachment, as a proxy for muscle relaxation, the samples were skinned, subject to a range of ATP concentrations and mechanics were measured. The samples were bathed in a mild detergent, 0.1% Triton X-100, to remove lipid membranes from the myocytes, sculpted to cylindrical shape of at least 500 micron length and 140-200 micron diameter, attached to aluminum clips, and placed between a length motor and force transducer. Samples were stretched to 2.2 micron sarcomere length, and myofilaments were then activated with increasing concentrations of Ca^2+^ in the presence of magnesium ATP (MgATP) over a 5.0 mM to 0.01 mM range of concentrations. A double-exponential function was fitted to the change in recorded tension, in response to the quick stretch, to characterize the rate of tension release (krel), which corresponds to the rate of myosin cross-bridge detachment.

## Results

### Patient demographics and clinical data

Heart donors and patients with HCM across the 3 genotype groups (Genotype negative, myosin binding protein C (*MYBPC3*), or beta-myosin (*MYH7*) variants) were well matched for age and sex (**Table 1**). Most of the samples came from patients of white race. Twelve percent of patients with HCM were diabetic. Maximal wall thickness and left ventricular outflow tract gradients were similar between the 3 HCM groups. Mean ejection fraction was higher in the *HCM-MYBPC3* and HCM-*MYH7* patients compared to the donors. All patients were symptomatic prior to myectomy, classified as either New York Heart Association Class II or III, with a comparatively larger percentage of patients with HCM due to *MYH7* variants being Class III. All patients were receiving either a beta blocker or calcium channel blocker at the time of myectomy, with 4 patients in the HCM-*MYH7* group being on both medications.

**Table 1.** Patient Demographics and Clinical Data.

|  | ***Non-failing***<br>***(n=8)*** | ***HCM – No***<br>***sarcomere***<br>***variants***<br>***(n=12)*** | ***HCM -***<br>***MYBPC3***<br>***(n=19)*** | ***HCM - MYH7***<br>***(n=10)*** |
| --- | --- | --- | --- | --- |
| ***Age*** | 52 ± 5 | 53 ± 4 | 42 ± 4 | 47.1 ± 5 |
| ***Sex (% male)*** | 37.5 | 50 | 37 | 60 |
| ***Race (% White)*** | 87.5 | 100 | 89 | 90 |
| ***Diabetes (% yes)*** | 0 | 17 | 16 | 0 |
| ***Maximal wall thickness***<br>***(mm)*** | 10.5 ± 0.5 | 19.6 ± 1.6 | 23.4 ± 2.2 | 22 ± 1.9 |
| ***Ejection fraction (%)*** | 64 ± 1.5 | 68 ± 1.9 | 71 | 75 ± 1.2 |
| ***Maximal LVOT gradient***<br>***(mmHg)*** | – | 91 ± 16 | 86 ± 13 | 110 ± 11.6 |
| ***NYHA Class (% Class III)*** | 0 | 42 | 37 | 80 |
| ***Beta blockers (%)*** | 0 | 75 | 79 | 80 |
| ***Calcium channel***<br>***blockers (%)*** | 0 | 25 | 21 | 40 |

HCM = hypertrophic cardiomyopathy, LVOT = left ventricular outflow tract, NYHA = New York Heart Association. 4 patients in the HCM-MYH7 group were taking both beta blockers and calcium channel blockers.

### Proteomic profiling revealed marked differences in abundance of proteins involved in contractility, the extracellular matrix, and energy metabolism in HCM compared to control hearts

A total of 527 proteins were identified in all samples from control and HCM hearts (**Supplemental Table 1**). After adjusting for multiple comparisons, a statistically significant difference (P<0.0001) was found for 98 proteins in at least one HCM group and 46 proteins in all 3 HCM groups compared to control hearts (**Figure 1**). Differences in protein abundance between the HCM and control groups were largely independent of genotype.

**Figure 1.**
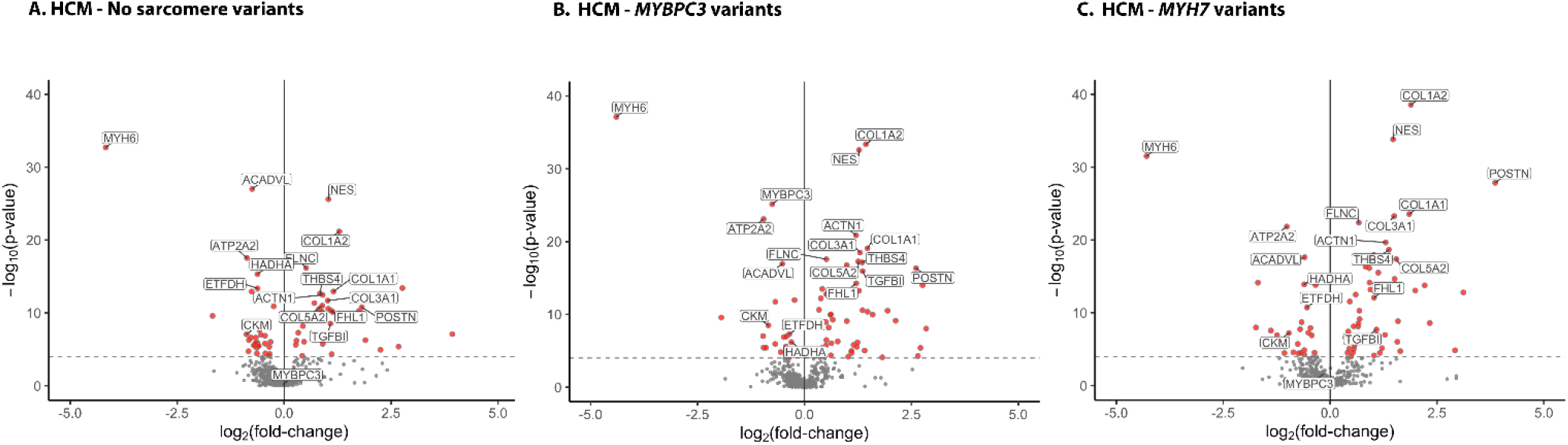
Volcano plots summarizing findings from proteomics analysis of human HCM and control hearts. Proteins with increased (+ log) or decreased abundance (- log) are shown as a function of fold change over control hearts for HCM samples with (**A**) no sarcomere variants, (**B**) MYBPC3 variants, and (**C**) MYH7 variants. Proteins that were significantly different from control, after Bonferroni correction for multiple comparisons (P<0.0001), are shown in red and a subset of the individual proteins are labeled. N=6 controls, 18 HCM-*MYBPC3*, 9 HCM-*MYH7*, and 12 HCM - No sarcomere variant.

Several sarcomeric proteins were differentially expressed in HCM compared to control hearts. For example, abundance of the alpha-myosin isoform (*MYH6*) was ~95% lower in HCM hearts. The alpha-myosin isoform comprises ~1% of total myosin in control hearts and 0.04% in HCM hearts, as the beta isoform (*MYH7*) is the predominate isoform expressed. There were no differences in abundance of the other major sarcomere protein isoforms among groups, except for a 40% reduction in myosin-binding protein C in the samples from patients with truncating *MYBPC3* variants, as we have shown previously ^25^.

Several Z-disc, intermediate filament, and cytoskeletal proteins were upregulated ~1.5-2.5-fold in HCM hearts compared to controls, including the skeletal isoform of alpha-actinin (*ACTN1*), Four and a half LIM domains protein 1 (*FHL1*), synaptopodin 2 (*SYNPO2L*), nestin (*NES*), filamin C (*FLNC*), vinculin (*VCL*) and obscurin (*OBSCN*). A number of extracellular matrix proteins were increased ~1.5-5-fold in HCM hearts compared to controls, including type 1, 5 and 6 collagens, fibronectin (*FN1*), thrombospondin 4 (*THBS4*), biglycan (*BGN*), versican (*VCAN*), and lumican (*LUM*). Cardiac fibroblast proteins established as important mediators of cardiac fibrosis were also more highly abundant in HCM compared to control hearts, including transforming growth factor 1 (*TGFB1*, ~2-2.5-fold) and periostin (*POSTN*, ~3.5-14-fold).^31^

The other major category of differentially regulated proteins between HCM and control hearts were those involved in cardiac fuel metabolism and energetics. The vast majority of these proteins were less abundant in HCM compared to control hearts, with the most highly statistically significant differences being in enzymes catalyzing key steps in long chain fatty acid beta-oxidation, including very long chain acyl-CoA dehydrogenase (*ACADVL*) and hydroxyacyl-CoA dehydrogenase (*HADHA*). A component of the electron-transfer system in mitochondria, electron transport flavoprotein dehydrogenase (*ETFDH*), was also significantly downregulated in HCM hearts. There were also ~50% reduction in creatine-kinase M-type, an enzyme involved in energy transduction, and in the ATPase sarcoplasmic reticulum calcium transporting protein 2 (*ATP2A2* that encodes SERCA2a), suggesting a significant impairment in the transfer and utilization of ATP in HCM hearts.

### Quantitative, targeted metabolomic analysis reveals marked depletion of fatty acid derivatives and ATP, increased ketone bodies, and increased branched chain amino acids in HCM samples

Given the lower abundance of multiple proteins involved in fuel and energy homeostasis in HCM hearts, we next sought to investigate metabolite pool sizes to identify precise pathway defects in cardiac energetics and metabolism. Using a targeted quantitative metabolomics platform, we identified a total of 103 metabolites, of which 58 significantly differed between at least one of the HCM groups compared to control samples after applying a false discovery rate adjustment (**Supplemental Table 2**). A principal component analysis scree plot showed that the first component accounted for most of the variability in metabolite abundance (**Supplemental Figure 1**). This first principal component was defined by disease status, and demonstrated overlapping clusters of the HCM groups, which were completely distinct from the control group (**Figure 2A**). Metabolite abundance was independent of genotype, sex, and age (**Supplemental Figure 2**).

**Figure 2.**
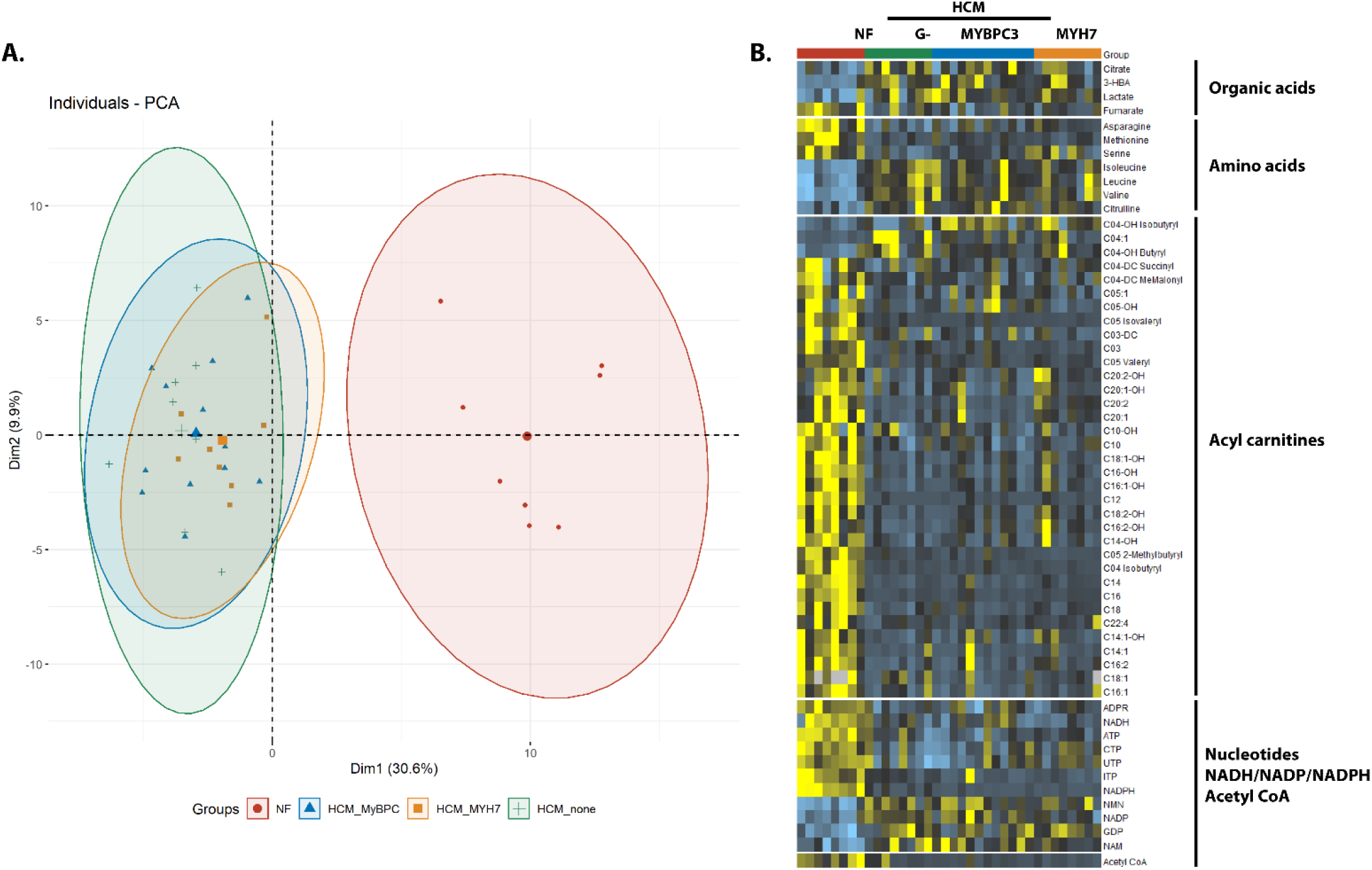
Metabolomics analysis of HCM and control hearts. **(A)** PCA plot showing the overall distinction between the control hearts and the 3 HCM groups. The 3 HCM groups overlap with each other with more variability in the HCM – no sarcomere variant group. **(B)** Heat map showing the individual metabolites that were significantly different in the HCM compared to the control heart samples after adjustment by false discovery rate. N=8 for controls, HCM-*MYH7*, HCM-no sarcomere variant and N=12 for HCM-*MYBCP3*.

The most striking difference in individual metabolites was the uniform depletion of mitochondrial fatty acid oxidation intermediates in HCM compared to control hearts, particularly long-chain acyl carnitines which were reduced ~70-95% (**Figure 2B and 3**). Conversely, the ketone body 3-hydroxybutyrate, lactate, and the 3 branched chain amino acids (BCAAs = leucine, isoleucine and valine) were all significantly increased in HCM hearts (**Figure 2B, Figure 3 and 4**). The breakdown carnitine products for ketones and valine respectively, C04-OH Butyryl and C04-OH Isobutyryl, were also increased in HCM hearts. Lactate, ketones and BCAAs may serve as alternate sources to fatty acids for the production of ATP, and have been shown to be increased in failing human hearts and in animal models of heart failure.^29, 32–35^ Most TCA cycle intermediates were not different among the groups, except for an increase in citrate and decrease in fumarate in the HCM compared to control hearts (**Figure 2B, Supplemental Table 2**). Of the non-BCAA amino acids, asparagine, methionine and serine were reduced and citrulline was increased in HCM compared to control hearts.

**Figure 3.**
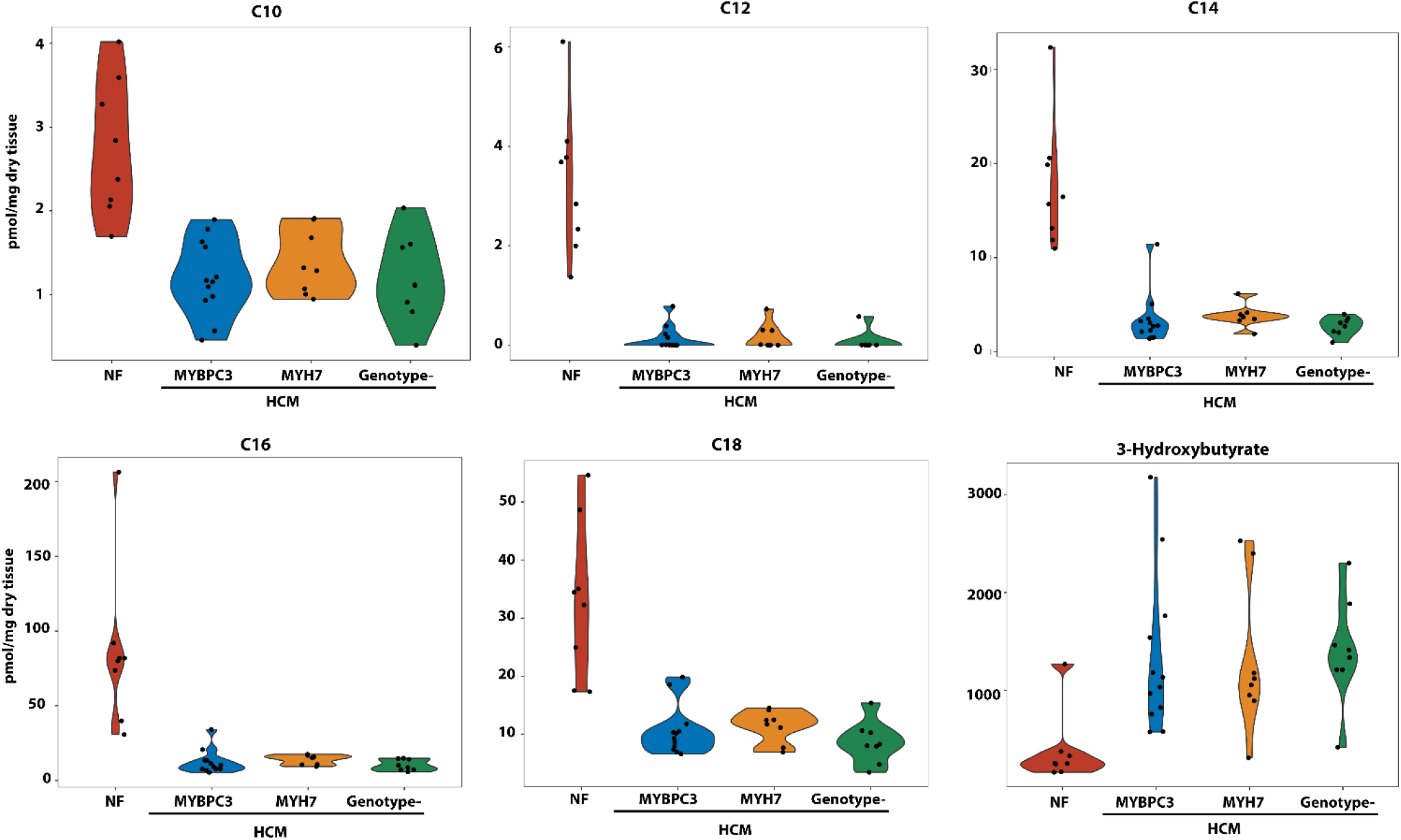
Metabolomic analysis showing representative abundances of long-chain acyl-carnitines and ketone bodies in HCM and control hearts. Abundances of C10, C12, C14, C16 and C18 were markedly and uniformly depleted in all HCM groups, independent of genotype, compared to control hearts. Ketone bodies were increased in abundance across all 3 HCM groups. ANOVA with Tukey post-hoc comparison were used to compare groups with P<0.0001 for all comparisons of HCM groups to control. N is the same as for Figure 2.

**Figure 4.**
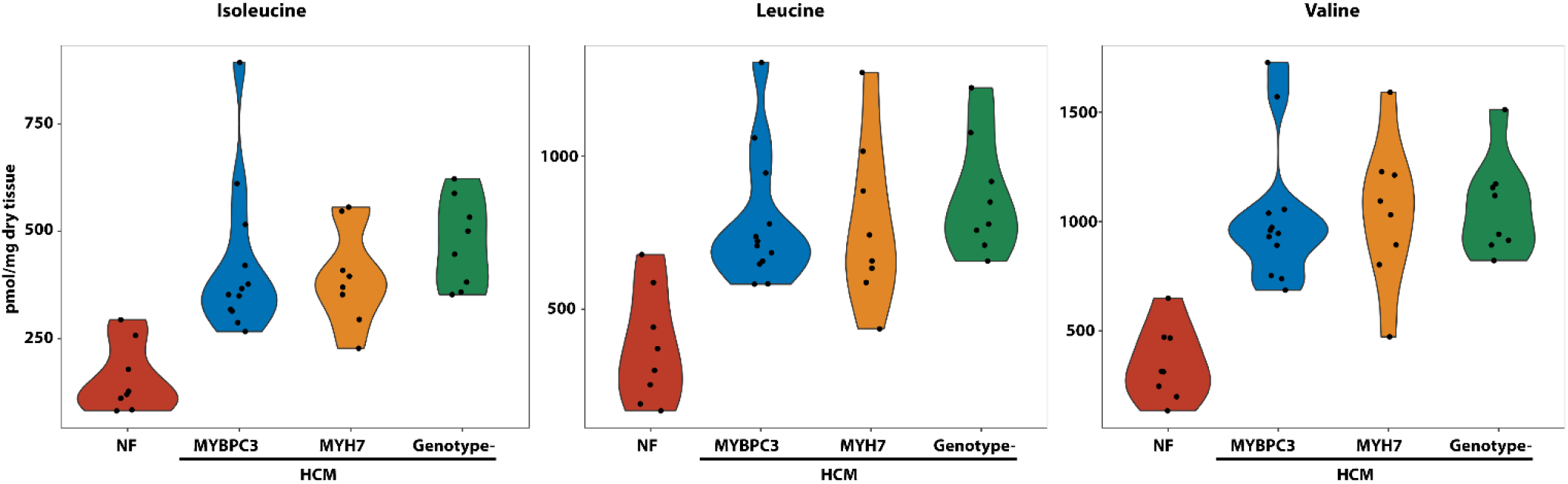
Metabolomic analysis showing increased abundance of branched chain amino acids in the hearts of patients with HCM compared to control. N is the same as for Figures 2 and 3. P<0.0001 for all comparisons of HCM to control hearts.

The nucleotides ATP, CTP, and UTP were all markedly reduced in abundance in HCM compared to control hearts (**Figure 2B, Figure 5**). Nicotinamide adenine dinucleotide (NADH) and its phosphate derivatives, NADP and NADPH, were also significantly reduced, while their precursors, nicotinamide (NAM) and NAM mononucleotide (NMN), were increased in HCM hearts. Acetyl CoA, the central molecule that delivers acetyl groups to the citric acid cycle to be oxidized for ATP production, was markedly reduced in HCM hearts. Together, this metabolomic analysis indicates a profound deficit in cardiac energetics, with depletion of fatty acid acyl carnitines likely driving a compensatory increase in alternate fuel sources, including ketone bodies, lactate and branched chain amino acids.

**Figure 5.**
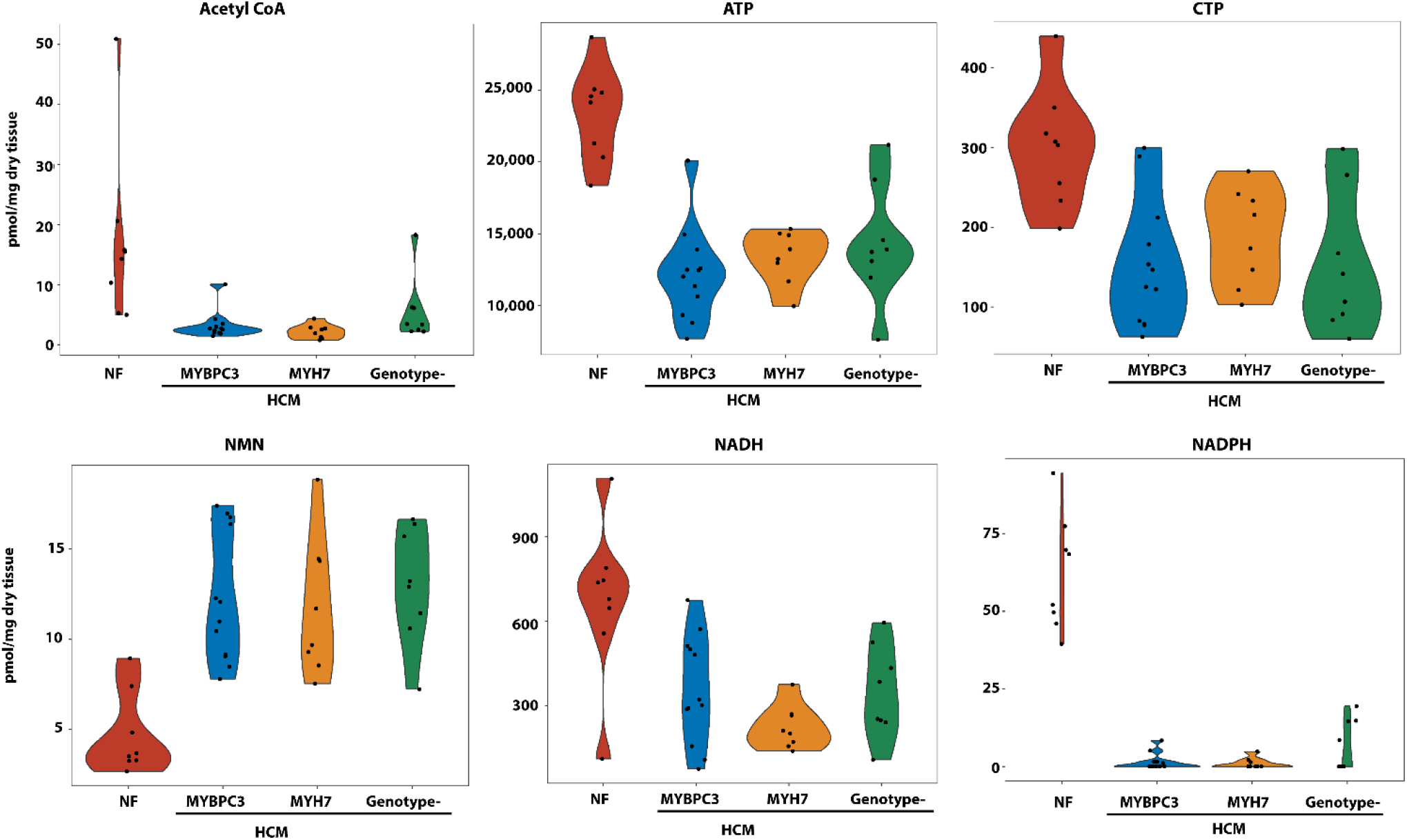
Metabolomics analysis showing depletion of acetyl CoA, trinucleotides, NADH and NADPH with increased abundance of NMN, the precursor of NAD. N=same as for Figure 3 and 4. P<0.0001 for all comparisons of HCM to control samples.

### ATP-dependent cross-bridge detachment in skinned human cardiac muscle

In order to determine the potential impact of the observed reduced ATP concentration in the HCM hearts on cardiac mechanics, we measured the kinetics of cross-bridge detachment in skinned human cardiac fibers as a function of ATP (**Figure 6**). The application of a quick 1% stretch in muscle length (**Figure 6A**) at maximal Ca^2+^-activation conditions (pCa 4.8) resulted in a rapid increase in tension, followed by a slower decrease as the myosin cross-bridges detached from the actin filament (**Figure 6B**). The rate of the cross-bridge release (krel) was slowed as the ATP concentration was diminished over the range (5.0 mM to 0.01 mM) of ATP concentrations examined (**Figure 6C**). Contrasting the mean and range of values for ATP content in HCM and control samples, we would predict that the magnitude reduction in ATP content in the HCM hearts would result in ~10% mean decrease in the rate of cross-bridge detachment when compared to control hearts.

**Figure 6.**
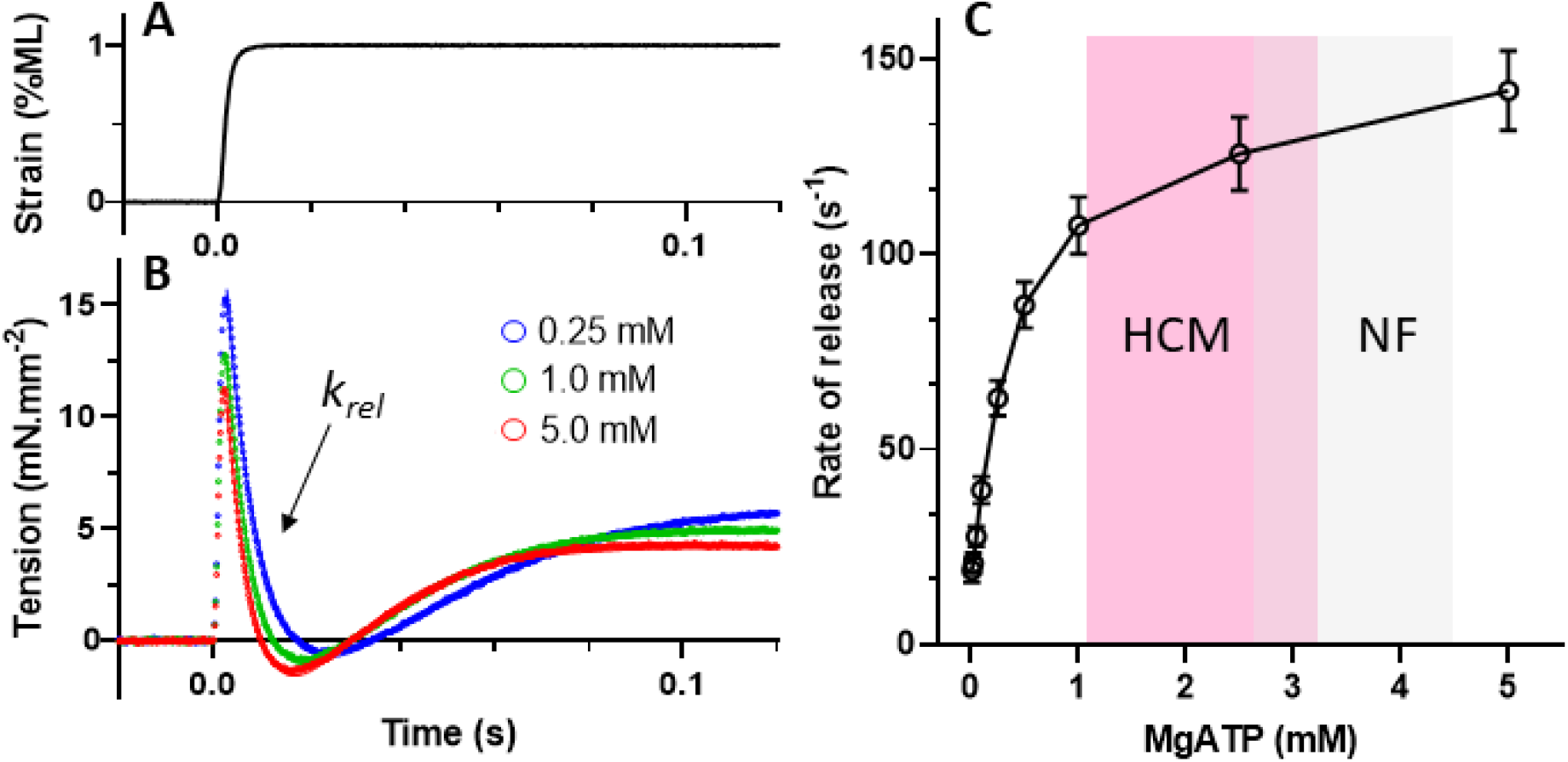
Rate of myosin cross-bridge detachment slows at lower MgATP concentrations. **A.** A quick stretch of 1% muscle length (ML) was applied to demembranated epicardial muscle that had been biopsied from patients undergoing coronary bypass grafting. **B.** The recorded tension response (solid lines) at maximum Ca^2+-^activation was fit to the double-exponential function (dotted lines). The rate of myosin crossbridge detachment is reflected in the rate of tension release (krel) immediately following the stretch. **C.** Lower concentrations of MgATP resulted in slower krel and would be expected to reduce relaxation function in the cardiac cycle during transition from late systole to early diastole. Mean ± SEM, n=32 samples from 14 patients.

## Discussion

Our comprehensive proteomic analysis of heart tissue from patients with HCM demonstrated an increased abundance of extracellular matrix proteins and mediators of cardiac fibrosis. This is an expected finding that is consistent with prior studies.^11, 13^ We also observed an increased abundance of multiple Z-disc and intermediate filament proteins in HCM hearts. Many of these proteins have well-established roles in sensing biomechanical stress, signal transduction, and myofibrillar repair and stabilization,^36, 37^ and an increase in their abundance may be an adaptive response to increased mechanical stress.^38^ In pairing proteomics with metabolomics analyses, we identified multiple defects in cardiac metabolism and energetics in heart samples obtained from patients with HCM compared to control, non-failing hearts. The reduction in the abundances of enzymes involved in long chain fatty acid oxidation correlated with a marked depletion of the corresponding acyl-carnitines, ATP, NADH and its phosphate derivatives, and acetyl-CoA. We also observed compensatory increases in ketone bodies, lactate and branched chain amino acids that may serve as alternate fuel sources. Finally, we show that the magnitude of reduction in ATP would be predicted to result in a decreased rate of actomyosin cross-bridge detachment that could contribute directly to diastolic dysfunction.

The energetic deficit in HCM is thought to relate primarily to increased ATP utilization at the sarcomere due to increased myofilament calcium sensitivity and/or a reduced number of cross-bridges in the super relaxed state as a fundamental consequence of pathogenic sarcomere variants.^39, 40^ This long-standing paradigm is supported by findings of reduced PCr:ATP ratios and myocardial efficiency in gene variants carriers even prior to the development of overt hypertrophy.^18^ How myocardial substrate availability is altered in response to this increase in ATP utilization in patients with hypertrophic cardiomyopathy has not been previously examined, but is critical for consideration of potential treatment modalities targeting cardiac metabolism. In this study, we observed a marked reduction in the abundance of proteins involved in β-oxidation of fatty acids and depletion of acyl carnitines in hearts from patients with HCM compared to control hearts. These findings are highly reminiscent of those from explanted hearts from patients with end-stage heart failure with reduced ejection fraction.^41^ Similarly, increased levels of lactate, the ketone body β-hydroxybutyrate, and the 3 branched chain amino acids in HCM hearts have also been observed in human failing hearts.^33, 35^ The fact that the metabolic signature of HCM is so similar to end-stage heart failure in terms of the direction and magnitude of change would not have been anticipated from experimental data, where no differences in metabolite levels were observed in the setting of pathologic hypertrophy in the absence of LV dysfunction or heart failure.^21^ Patients with HCM undergoing myectomy, while symptomatic from left ventricular outflow tract obstruction, are typically well compensated and have normal to increased left ventricular ejection fractions (Table 1). The surprising similarities in the metabolic profile of HCM and more advanced heart failure therefore suggest that these metabolic changes occur relatively early in the progressive stages of cardiomyopathy and heart failure, and represent a canonical response to different forms of pathologic stress and remodeling, independent of the inciting trigger. Recently it has been shown that reduced flux through fatty acid oxidation and increased glucose utilization is necessary for cardiac hypertrophic growth.^42^ Therefore, therapies aimed at restoring these metabolic defects have the potential to prevent or attenuate adverse cardiac hypertrophic remodeling and could have widespread benefit across a spectrum of cardiomyopathies.

The marked metabolic shift in substrate availability in human HCM has important therapeutic implications. Metabolic modulation is an attractive strategy because energy deprivation is considered likely to contribute directly to the pathophysiology of HCM. Indeed, our studies using skinned human cardiac trabeculae suggest that ATP depletion to levels measured in HCM hearts could contribute directly to diastolic dysfunction by decreasing the rate of actomyosin cross-bridge attachment. Since these metabolic defects were common across all genotypic forms of HCM, the therapeutic benefit of metabolic modulation would also be expected to be generalizable. Perhexiline was the first metabolic modulator to be tested in a clinical trial of patients with HCM.^43^ Perhexiline is an inhibitor of CPT1, which transports fatty acids across the outer mitochondrial membrane. The rationale behind this strategy was to induce a metabolic substrate shift away from fatty acids toward glycolysis which consumes less oxygen per ATP generated. This phase 2 trial showed some modest improvements in symptoms, diastolic filling time and cardiac energetics in patients taking perhexiline relative to placebo-treated patients, but the drug was not pursued further because of multi-organ toxicity. Trimetazidine, an inhibitor or 3 ketoacyl-coA thiolase that catalyzes the last step in fatty acid β-oxidation to acetyl CoA, was also tested in a subsequent randomized, placebo-controlled phase 2 trial of patients with nonobstructive HCM.^44^ Patients treated with trimetazidine demonstrated a decrease in their exercise capacity relative to patients treated with placebo and no improvements in any of the secondary endpoints. This trial therefore signaled a potential for harm of inhibiting fatty acid β-oxidation in patients with HCM. This outcome is perhaps not surprising, in light of the profound disease-related deficits in fatty acid oxidation and depletion of acyl carnitines in the hearts of patients with HCM demonstrated in the present study.

A more promising therapeutic approach would be to bypass the deficit in fatty acid oxidation by enhancing the availability of alternate fuel sources for the production of ATP. Therapeutic ketosis is one such approach that has recently garnered considerable interest for treating patients with heart failure.^22, 45, 46^ An elevation in ketone bodies has been observed in hearts from patients with end-stage heart failure with reduced ejection fraction,^33^ and the contribution of ketones to ATP production in the human failing heart was recently shown to be 16.4%, almost 3-fold higher than the non-failing heart.^32^ Ketone body metabolism has been shown to represent an adaptive response to energy deprivation in animal models,^47^ further supporting the concept of a metabolic shift toward ketone metabolism as a viable therapeutic strategy. The ketone body, β-hydroxybutyrate, was elevated ~3.5-fold in hearts of patients with HCM compared to controls hearts, which we presume to be an adaptive response to the decreased availability of acyl carnitines. This response could be augmented pharmacologically, either by direct administration of ketone ester formulations,^45, 48^ or by drugs that stimulate ketone body production from the liver. This is one of several mechanism by which sodium glucose co-transporter inhibitors (SGLT2i) have been postulated to afford marked cardiovascular benefit in patients with heart failure and reduced or preserved ejection fraction, recently demonstrated in large randomized clinical trials.^49, 51^ The similar metabolic profile in HCM and failing hearts suggests that the benefits of SGLT2i could extend to patients with HCM, and provides a rationale for future clinical trials.

The branched chain amino acids, valine, leucine, and isoleucine, were elevated 2.5-3 fold in HCM hearts compared to control, similar to what has been demonstrated in experimental models and in human heart failure.^34, 35^ This may also represent an adaptive response to provide an alternate fuel source when fatty acids are depleted. However, the relative contribution of amino acids to the production of ATP is quite small (<5%),^32^ and it is unclear whether an increased abundance of BCAAs would make a meaningful difference. Suppression of BCAA catabolism appears to be a maladaptive component of myocardial remodeling,^52^ and therefore enhancement of BCAA catabolism in the heart or other organs may still be a reasonable target for reasons other than enhancing ATP production.

Modulation of NAD+ metabolism is another emerging strategy for therapeutic application in heart failure. Reduced bioavailability of NAD+ has been reported with aging and heart failure with reduced ejection fraction, and most recently in an experimental model of heart failure with preserved ejection fraction.^53^ In the latter study, administration of the NAD+ precursor, nicotinamide riboside (NR), or an activator of NAMPT, both attenuated cardiac dysfunction. Most of NAD+ in the heart is synthesized via the salvage pathway.^54^ Nampt, which catalyzes the conversion of NAM to NMN, is downregulated in pressure overload hypertrophy and myocardial ischemia. Interestingly, both NAM and NMN were significantly elevated in human HCM hearts compared to control hearts, suggesting that Nampt was not the rate limiting enzyme for NAD+ production. Since ATP is required for the conversion of NAM to NMN, and NMN to NAD+, perhaps the reduced bioavailability of NADH, NADP and NADPH in HCM hearts is related to ATP depletion. If that were the case, then provision of NAD+ precursors or NAMPT activators would be predicted not to have therapeutic benefit in HCM.

Our study has some limitations. First, heart samples are only available from patients with symptomatic obstructive HCM. Therefore, we are unable to determine whether these findings are generalizable to patients with non-obstructive HCM, or to patients at an earlier stage of disease, including pre-clinical sarcomere variant carriers. As with any study comparing diseased to non-diseased hearts, hearts from patients with HCM are more fibrotic, as evidenced by the increase in collagens and other extracellular matrix proteins. This could in theory cause a “dilutional” effect of proteins and metabolites from cardiomyocytes. However, increases in multiple cardiomyocyte proteins (such as Z-disc proteins) and metabolites (such as BCAAs, β-hydroxybutyrate and lactate) in HCM compared to control hearts would argue against this being a significant issue. Finally, the kinetics of actomyosin cross bridge attachment and ATP response curves may differ in HCM hearts compared to those measured in trabeculae from patients with coronary heart disease.

In conclusion, we identified profound deficits in cardiac energetics in heart tissue from patients with HCM, marked by depletion of fatty acid derivatives and compensatory increases in metabolites that could serve as alternate fuel sources. These results provide significant biological insights that lend support to the paradigm that energy depletion contributes to the pathophysiology of HCM. These findings also have important therapeutic implications, and suggest that strategies aimed to reduce the capacity of the HCM heart to oxidize fatty acids could be ineffective or potentially harmful. Alternatively, augmentation of compensatory responses to increase utilization of alternate fuel sources, particularly ketones, holds promise for treatment of a variety of patients with different forms of cardiac hypertrophic remodeling that lead to heart failure.

## Non-Standard Abbreviations and Acronyms

None

## Acknowledgements

We would like to thank the Penn Metabolomics Core for conducting the metabolomics studies.

