## Supplemental Table 1 for "Metabolic and Proteomic Defects in Human Hypertrophic Cardiomyopathy"

| | | | | | Median Abundances (Top 5) | | | | SD of Median Abundances (Top 5) | | | | Ratio over GOL | | | SD of Ratio over GOL | | | t-test with GOL | | |
|---|---|---|---|---|---|---|---|---|---|---|---|---|---|---|---|---|---|---|---|---|---|
| GENE | # Peptides | Accession | Group | Sub-Group | Control | MYBPC3 | No variants | MYH7 | Control2 | MYBPC32 | No variar | MYH73 | MYBPC33 | No variants3 | MYH74 | MYBPC34 | No variant: | MYH75 | MYBPC35 | No variants! | MYH72 |
| | | | **Sarcomere** | | | | | | | | | | **0.989** | **0.943** | **0.891** | | | | 0.725913 | 0.4084332 | 0.0364853 |
| | | | **Sarcomere** | **Thick Filament** | 0 | 0 | 0 | 0 | | | | | **0.992** | **0.950** | **0.898** | | | | 0.33104 | 0.6240197 | 0.232321 |
| MYL2 | 14 | Q6IB42 | Sarcomere | Thick Filament | 0.8319 | 0.8806 | 0.8461 | 0.8482 | 0.125 | 0.149 | 0.133 | 0.067 | **1.019** | **0.731** | **0.477** | 0.977 | 0.825 | 0.703 | 0.6536411 | 0.0481251 | 3.405E-05 |
| MYL3 | 12 | A0A024R2Q5 | Sarcomere | Thick Filament | 0.9157 | 0.9148 | 0.8724 | 0.9312 | 0.110 | 0.088 | 0.136 | 0.051 | **0.995** | **0.950** | **0.970** | 1.715 | 0.569 | 0.483 | 0.8830684 | 0.3534733 | 0.3635753 |
| MYL4 | 13 | P12829 | Sarcomere | Thick Filament | 0.0045 | 0.0048 | 0.0192 | 0.0094 | 0.021 | 0.012 | 0.046 | 0.022 | **1.107** | **3.554** | **1.643** | 6.852 | 21.952 | 8.980 | 0.8465359 | 0.0054443 | 0.3991849 |
| MYL6 | 6 | G3V1V0 | Sarcomere | Thick Filament | 0.0027 | 0.0032 | 0.0030 | 0.0033 | 0.001 | 0.006 | 0.001 | 0.002 | **1.198** | **1.114** | **1.183** | 2.113 | 0.634 | 0.730 | 0.0120306 | 0.1702306 | 0.0154212 |
| MYL7 | 5 | Q01449 | Sarcomere | Thick Filament | 0.0009 | 0.0012 | 0.0013 | 0.0016 | 0.001 | 0.001 | 0.001 | 0.001 | **1.357** | **1.094** | **1.736** | 1.555 | 1.137 | 2.274 | 0.1113916 | 0.0171829 | 0.2580758 |
| MYH6 | 15 | P13533 | Sarcomere | Thick Filament | 0.0125 | 0.0005 | 0.0005 | 0.0005 | 0.018 | 0.000 | 0.001 | 0.000 | **0.047** | **0.056** | **0.051** | 0.125 | 0.143 | 0.117 | 7.389E-38 | 1.923E-33 | 3.056E-32 |
| MYH7 | 15 | A5YM51; P12883 | Sarcomere | Thick Filament | 1.1942 | 1.1758 | 1.1995 | 1.2123 | 0.085 | 0.091 | 0.153 | 0.111 | **0.994** | **0.995** | **1.016** | 0.190 | 0.155 | 0.122 | 0.9578138 | 0.3490311 | 0.4368207 |
| MYH6&MYH7 | 13 | A5YM51; P12883; | Sarcomere | Thick Filament | 1.1798 | 1.1678 | 1.1919 | 1.2015 | 0.085 | 0.060 | 0.082 | 0.141 | **0.998** | **1.008** | **0.998** | 0.148 | 0.153 | 0.143 | 0.9763714 | 0.9580033 | 0.9556216 |
| MYBPC3 (cDNA) | 2 | A8K6A4 | Sarcomere | Thick Filament | 0.0364 | 0.0258 | 0.0404 | 0.0326 | 0.016 | 0.010 | 0.016 | 0.006 | **1.116** | **1.186** | **1.526** | 2.559 | 2.626 | 3.467 | 0.7771505 | 0.6918864 | 0.4779739 |
| MYBPC3 protein | 6 | A5YM48 | Sarcomere | Thick Filament | 0.0680 | 0.0413 | 0.0665 | 0.0646 | 0.011 | 0.008 | 0.011 | 0.006 | **0.596** | **0.936** | **0.878** | 0.280 | 0.440 | 0.300 | 0.0198822 | 0.6640227 | 0.4095543 |
| MYBPC3 | 13 | A5YM48; A8K6A4 | Sarcomere | Thick Filament | 0.1049 | 0.0608 | 0.1032 | 0.0959 | 0.019 | 0.013 | 0.018 | 0.015 | **0.592** | **0.984** | **0.900** | 0.146 | 0.239 | 0.191 | 7.19E-26 | 0.653311 | 0.0251876 |
| | | | **Sarcomere** | **M-line** | 0 | 0 | 0 | 0 | | | | | **0.849** | **0.864** | **0.750** | | | | 0.0055318 | 0.0048512 | 1.845E-07 |
| MYOM3 | 12 | Q5VTT5 | Sarcomere | M-line | 0.0096 | 0.0091 | 0.0091 | 0.0081 | 0.002 | 0.002 | 0.002 | 0.001 | **0.936** | **0.958** | **0.832** | 0.297 | 0.317 | 0.241 | 0.6300276 | 0.4004409 | 0.0192289 |
| MYOM1 | 5 | P52179 | Sarcomere | M-line | 0.0072 | 0.0059 | 0.0063 | 0.0060 | 0.001 | 0.001 | 0.001 | 0.001 | **0.813** | **0.889** | **0.856** | 0.267 | 0.283 | 0.229 | 0.0257246 | 0.1738018 | 0.0237294 |
| MYOM1 | 12 | P52179; A8MX12 | Sarcomere | M-line | 0.0201 | 0.0190 | 0.0198 | 0.0171 | 0.003 | 0.002 | 0.002 | 0.002 | **0.906** | **0.936** | **0.795** | 0.245 | 0.257 | 0.190 | 0.2416202 | 0.4524625 | 0.0001709 |
| MYOM2 | 13 | P54296; A0A024Q: | Sarcomere | M-line | 0.0256 | 0.0200 | 0.0191 | 0.0155 | 0.005 | 0.004 | 0.004 | 0.006 | **0.765** | **0.735** | **0.615** | 0.218 | 0.220 | 0.269 | 2.405E-05 | 1.646E-06 | 1.864E-08 |
| | | | **Sarcomere** | **Thin Filament** | 0 | 0 | 0 | 0 | | | | | **0.926** | **0.869** | **0.814** | | | | 0.6157234 | 0.2656445 | 0.0739922 |
| ACT | 11 | P62736; P68133 | Sarcomere | Thin Filament | 1.4158 | 1.2653 | 1.2062 | 1.1071 | 0.317 | 0.276 | 0.334 | 0.142 | **0.921** | **0.888** | **0.815** | 1.299 | 0.922 | 1.383 | 0.5022416 | 0.016453 | 0.0001386 |
| TNNC1 | 7 | Q6FH91 | Sarcomere | Thin Filament | 0.0968 | 0.0865 | 0.0760 | 0.0779 | 0.040 | 0.022 | 0.032 | 0.016 | **0.664** | **0.521** | **0.455** | 0.463 | 0.385 | 0.277 | 0.2259676 | 0.1720074 | 0.0490155 |
| TNNT2 | 10 | P45379 | Sarcomere | Thin Filament | 0.2059 | 0.1915 | 0.1882 | 0.1642 | 0.041 | 0.047 | 0.047 | 0.019 | **0.996** | **0.923** | **0.824** | 0.719 | 0.566 | 0.682 | 0.7797887 | 0.6378496 | 0.215476 |
| TNNI1 | 5 | A0A087WT47 | Sarcomere | Thin Filament | 0.0009 | 0.0016 | 0.0020 | 0.0014 | 0.001 | 0.004 | 0.003 | 0.001 | **1.835** | **3.105** | **1.931** | 4.099 | 4.799 | 1.953 | 0.02454 | 0.000368 | 0.0317287 |
| TNNI3 | 12 | B6D427; Q6FGU5 | Sarcomere | Thin Filament | 0.2252 | 0.2117 | 0.1905 | 0.2072 | 0.036 | 0.036 | 0.040 | 0.022 | **0.896** | **0.814** | **0.850** | 0.754 | 0.590 | 0.689 | 0.3312262 | 0.2567185 | 0.150296 |
| TPM1 | 10 | H0YNC7; A0A024R | Sarcomere | Thin Filament | 0.2818 | 0.2415 | 0.2066 | 0.2206 | 0.060 | 0.050 | 0.045 | 0.023 | **0.974** | **0.821** | **0.851** | 0.831 | 0.497 | 0.485 | 0.9292395 | 0.1411896 | 0.2537791 |
| TPM2 | 5 | Q5TCU3 | Sarcomere | Thin Filament | 0.0073 | 0.0054 | 0.0066 | 0.0067 | 0.003 | 0.005 | 0.004 | 0.002 | **0.819** | **0.737** | **0.711** | 10.240 | 0.887 | 1.085 | 0.9962768 | 0.7967527 | 0.7904407 |
| TPM2 shared | 4 | Q5TCU3; B4E3P1; , | Sarcomere | Thin Filament | 0.0108 | 0.0093 | 0.0080 | 0.0086 | 0.002 | 0.005 | 0.003 | 0.002 | **0.884** | **0.756** | **0.805** | 0.825 | 0.719 | 0.751 | 0.660231 | 0.3887749 | 0.7013934 |
| TPM3 | 9 | P06753 | Sarcomere | Thin Filament | 0.0011 | 0.0035 | 0.0042 | 0.0029 | 0.001 | 0.005 | 0.008 | 0.004 | **3.843** | **3.362** | **3.006** | 6.709 | 8.124 | 4.808 | 3.145E-11 | 5.417E-11 | 1.429E-09 |
| NEBL | 14 | O76041 | Sarcomere | Thin Filament | 0.0264 | 0.0186 | 0.0220 | 0.0307 | 0.026 | 0.016 | 0.016 | 0.020 | **0.851** | **0.868** | **0.890** | 0.271 | 0.303 | 0.369 | 0.0214322 | 0.0497194 | 0.1884036 |
| CALD1 | 5 | Q05682 | Sarcomere | Thin Filament | 0.0005 | 0.0007 | 0.0006 | 0.0008 | 0.000 | 0.002 | 0.000 | 0.000 | **1.316** | **1.126** | **1.562** | 3.925 | 0.703 | 1.002 | 0.0025465 | 0.2833199 | 0.0016272 |
| | | | **Sarcomere** | **Titan** | 0 | 0 | 0 | 0 | | | | | **0.987** | **1.005** | **0.913** | | | | 0.3599452 | 0.773824 | 0.8536117 |
| TTN | 13 | Q8WZ42 | Sarcomere | Titan | 0.0213 | 0.0228 | 0.0224 | 0.0229 | 0.002 | 0.002 | 0.002 | 0.002 | **1.045** | **1.050** | **0.996** | 0.287 | 0.314 | 0.240 | 0.2955684 | 0.465375 | 0.7049998 |
| TTN (upreg section) | 12 | C0JYZ2; Q8WZ42 | Sarcomere | Titan | 0.0006 | 0.0012 | 0.0010 | 0.0010 | 0.001 | 0.001 | 0.001 | 0.000 | **2.215** | **1.757** | **1.919** | 2.343 | 2.001 | 2.141 | 1.572E-06 | 0.0073716 | 0.0009781 |
| TTN shared | 12 | C0JYZ2; Q8WZ42 | Sarcomere | Titan | 0.0901 | 0.0744 | 0.0727 | 0.0703 | 0.032 | 0.024 | 0.030 | 0.009 | **0.959** | **0.985** | **0.895** | 0.384 | 0.317 | 0.315 | 0.8038196 | 0.3160666 | 0.3082739 |
| TCAP | 5 | A2TDC0 | Sarcomere | Titan | 0.0048 | 0.0038 | 0.0044 | 0.0032 | 0.002 | 0.002 | 0.002 | 0.001 | **0.802** | **0.938** | **0.371** | 0.620 | 0.643 | 0.398 | 0.7288304 | 0.8544163 | 0.0696819 |
| | | | **Sarcomere** | **Z-line** | 0 | 0 | 0 | 0 | | | | | **1.352** | **1.276** | **1.219** | | | | 0.0063515 | 0.0753668 | 0.0585214 |
| ACTN1 | 13 | P12814 | Sarcomere | Z-line | 0.0039 | 0.0084 | 0.0068 | 0.0087 | 0.002 | 0.003 | 0.002 | 0.003 | **2.305** | **1.867** | **2.460** | 1.273 | 0.851 | 1.224 | 1.375E-21 | 3.375E-13 | 2.149E-20 |
| ACTN2 | 13 | P35609 | Sarcomere | Z-line | 0.1833 | 0.1881 | 0.1904 | 0.1793 | 0.018 | 0.017 | 0.017 | 0.017 | **1.023** | **1.031** | **0.953** | 0.155 | 0.159 | 0.138 | 0.3245612 | 0.5660503 | 0.0740763 |
| ACTN4 | 8 | O43707 | Sarcomere | Z-line | 0.0182 | 0.0108 | 0.0147 | 0.0350 | 0.036 | 0.021 | 0.020 | 0.027 | **1.196** | **0.973** | **1.085** | 1.801 | 0.843 | 1.039 | 0.6927938 | 0.8322192 | 0.9637157 |
| CAPZA2 | 6 | A4D0V4 | Sarcomere | Z-line | 0.0041 | 0.0032 | 0.0030 | 0.0027 | 0.001 | 0.001 | 0.001 | 0.001 | **0.759** | **0.730** | **0.614** | 0.284 | 0.293 | 0.196 | 0.0345849 | 0.0077724 | 0.0004906 |
| FHL1 | 12 | Q53FI7; B7Z5V0 | Sarcomere | Z-line | 0.0373 | 0.0825 | 0.0792 | 0.0776 | 0.013 | 0.020 | 0.018 | 0.014 | **2.311** | **2.138** | **2.037** | 1.139 | 1.066 | 0.961 | 5.455E-15 | 4.879E-11 | 8.601E-13 |
| MYOZ2 | 12 | Q9NPC6 | Sarcomere | Z-line | 0.0536 | 0.0445 | 0.0454 | 0.0442 | 0.008 | 0.006 | 0.008 | 0.007 | **0.864** | **0.847** | **0.759** | 0.333 | 0.302 | 0.233 | 0.2098664 | 0.1802384 | 0.0248235 |
| | | | **Cytoskeletal** | | | | | | | | | | **1.042** | **0.967** | **0.990** | | | | 1.014E-06 | 0.0024442 | 9.851E-07 |
| | | | **Cytoskeletal** | **Actin** | 0 | 0 | 0 | 0 | | | | | **1.370** | **1.194** | **1.390** | | | | 3.276E-06 | 0.006845 | 9.996E-07 |
| SYNPO2L | 12 | Q9H987 | Cytoskeletal | Actin | 0.0095 | 0.0181 | 0.0157 | 0.0190 | 0.003 | 0.004 | 0.004 | 0.004 | **1.983** | **1.632** | **2.033** | 0.732 | 0.666 | 0.763 | 1.693E-17 | 4.463E-12 | 1.134E-18 |
| SYNPO2 | 10 | B9EG60 | Cytoskeletal | Actin | 0.0027 | 0.0033 | 0.0031 | 0.0034 | 0.001 | 0.001 | 0.001 | 0.001 | **1.192** | **1.132** | **1.123** | 0.392 | 0.294 | 0.307 | 0.0103997 | 0.0504115 | 0.020287 |
| SYNPO | 9 | Q8N3V7 | Cytoskeletal | Actin | 0.0053 | 0.0065 | 0.0062 | 0.0074 | 0.001 | 0.002 | 0.002 | 0.002 | **1.481** | **1.232** | **1.335** | 0.486 | 0.396 | 0.474 | 0.0317421 | 0.2498257 | 0.0143654 |
| CFL2 | 6 | Q549N0 | Cytoskeletal | Actin | 0.0103 | 0.0110 | 0.0105 | 0.0123 | 0.003 | 0.003 | 0.002 | 0.002 | **1.101** | **0.948** | **1.258** | 0.462 | 0.401 | 0.392 | 0.4245713 | 0.8104032 | 0.2177932 |
| WDR1 | 10 | Q59ER5 | Cytoskeletal | Actin | 0.0047 | 0.0054 | 0.0048 | 0.0049 | 0.001 | 0.001 | 0.001 | 0.001 | **1.104** | **1.063** | **1.081** | 0.495 | 0.333 | 0.302 | 0.0843449 | 0.7986798 | 0.2054019 |
| XIRP1 | 13 | Q702N8 | Cytoskeletal | Actin | 0.0061 | 0.0072 | 0.0069 | 0.0076 | 0.003 | 0.002 | 0.002 | 0.002 | **1.288** | **1.159** | **1.268** | 0.728 | 0.664 | 0.770 | 0.0045303 | 0.2313754 | 0.127607 |
| XIRP2 | 3 | A4UGR9 | Cytoskeletal | Actin | 0.0003 | 0.0007 | 0.0006 | 0.0007 | 0.000 | 0.000 | 0.000 | 0.000 | **2.337** | **1.832** | **2.321** | 1.685 | 1.385 | 1.334 | 2.837E-06 | 0.0003396 | 7.43E-06 |
| NEXN | 4 | Q0ZGT2 | Cytoskeletal | Actin | 0.0020 | 0.0021 | 0.0018 | 0.0025 | 0.001 | 0.001 | 0.000 | 0.001 | **1.066** | **0.956** | **1.095** | 0.499 | 0.395 | 0.365 | 0.416318 | 0.510019 | 0.1971458 |
| CFL1 | 6 | P23528 | Cytoskeletal | Actin | 0.0089 | 0.0092 | 0.0096 | 0.0099 | 0.002 | 0.002 | 0.002 | 0.002 | **1.023** | **1.059** | **1.065** | 0.388 | 0.367 | 0.300 | 0.8058478 | 0.6214506 | 0.2178111 |
| ABLIM1 | 12 | A0A0A0MRL6 | Cytoskeletal | Actin | 0.0080 | 0.0071 | 0.0073 | 0.0082 | 0.003 | 0.002 | 0.002 | 0.001 | **0.936** | **0.900** | **0.960** | 0.264 | 0.309 | 0.282 | 0.6138865 | 0.6312403 | 0.8500766 |
| | | | **Cytoskeletal** | **Intermediate Filament** | 0 | 0 | 0 | 0 | | | | | **0.965** | **0.874** | **0.891** | | | | 0.0051331 | 0.0439032 | 0.0090293 |
| ACTB shared | 4 | Q1KLZ0; P62736; P | Cytoskeletal | Intermediate Filament | 1.2095 | 1.3066 | 1.0876 | 1.2178 | 0.482 | 0.361 | 0.398 | 0.198 | **0.959** | **0.869** | **0.882** | 0.487 | 0.443 | 0.391 | 0.9994176 | 0.9442828 | 0.8931222 |
| SYNM | 12 | O15061 | Cytoskeletal | Intermediate Filament | 0.0030 | 0.0040 | 0.0034 | 0.0039 | 0.001 | 0.001 | 0.001 | 0.001 | **1.425** | **1.089** | **1.311** | 0.448 | 0.336 | 0.451 | 3.261E-07 | 0.0349452 | 0.0001339 |
| NES | 14 | D3DVC4 | Cytoskeletal | Intermediate Filament | 0.0019 | 0.0046 | 0.0039 | 0.0051 | 0.001 | 0.002 | 0.001 | 0.002 | **2.421** | **2.049** | **2.778** | 1.790 | 1.242 | 1.752 | 2.612E-33 | 2.475E-26 | 1.51E-34 |
| | | | **Cytoskeletal** | **Linker** | 0 | 0 | 0 | 0 | | | | | **1.302** | **1.313** | **1.284** | | | | 3.606E-06 | 0.0118927 | 0.0028713 |
| CTNNA1 | 12 | P35221 | Cytoskeletal | Linker | 0.0027 | 0.0031 | 0.0027 | 0.0033 | 0.000 | 0.001 | 0.001 | 0.001 | **1.130** | **1.015** | **1.257** | 0.801 | 0.598 | 0.537 | 0.0013027 | 0.7998327 | 0.0117832 |
| CTNNA3 | 11 | A8K141 | Cytoskeletal | Linker | 0.0027 | 0.0031 | 0.0031 | 0.0023 | 0.001 | 0.001 | 0.001 | 0.001 | **1.140** | **1.115** | **0.807** | 0.547 | 0.527 | 0.298 | 0.162145 | 0.2279306 | 0.2982084 |
| CTNNB1 | 10 | P35222 | Cytoskeletal | Linker | 0.0039 | 0.0049 | 0.0049 | 0.0047 | 0.001 | 0.001 | 0.001 | 0.001 | **1.193** | **1.323** | **1.189** | 0.571 | 0.650 | 0.539 | 0.0138634 | 0.0193198 | 0.5735773 |
| DMD | 14 | P11532 | Cytoskeletal | Linker | 0.0039 | 0.0043 | 0.0037 | 0.0037 | 0.001 | 0.001 | 0.001 | 0.000 | **1.041** | **0.924** | **0.874** | 0.329 | 0.307 | 0.237 | 0.7101469 | 0.3452482 | 0.0345823 |
| DMD | 14 | P11532; Q59HC3 | Cytoskeletal | Linker | 0.0043 | 0.0045 | 0.0040 | 0.0044 | 0.001 | 0.001 | 0.001 | 0.001 | **1.114** | **0.952** | **0.980** | 0.525 | 0.496 | 0.372 | 0.1834784 | 0.4931923 | 0.9284041 |
| DSTN | 6 | V9HWA6 | Cytoskeletal | Linker | 0.0063 | 0.0055 | 0.0050 | 0.0059 | 0.002 | 0.003 | 0.001 | 0.001 | **0.834** | **0.801** | **0.841** | 0.556 | 0.428 | 0.331 | 0.3767226 | 0.1996929 | 0.5975054 |
| EZR | 6 | E7EQR4 | Cytoskeletal | Linker | 0.0048 | 0.0040 | 0.0042 | 0.0048 | 0.001 | 0.001 | 0.001 | 0.001 | **0.890** | **0.910** | **0.905** | 0.460 | 0.530 | 0.382 | 0.6930332 | 0.5390769 | 0.4250225 |
| FLNA | 13 | P21333 | Cytoskeletal | Linker | 0.0022 | 0.0027 | 0.0025 | 0.0026 | 0.001 | 0.007 | 0.001 | 0.002 | **1.416** | **1.244** | **1.343** | 4.451 | 0.888 | 1.565 | 1.145E-09 | 0.0008715 | 3.813E-08 |
| FLNC | 15 | Q14315 | Cytoskeletal | Linker | 0.0269 | 0.0377 | 0.0374 | 0.0413 | 0.007 | 0.007 | 0.006 | 0.009 | **1.426** | **1.428** | **1.591** | 0.520 | 0.486 | 0.586 | 2.639E-18 | 6.843E-17 | 4.185E-23 |
| MSN | 5 | P26038 | Cytoskeletal | Linker | 0.0020 | 0.0021 | 0.0021 | 0.0020 | 0.001 | 0.001 | 0.001 | 0.000 | **0.973** | **0.971** | **0.954** | 0.575 | 0.504 | 0.552 | 0.5011005 | 0.8628239 | 0.89349 |
| PGM1 | 8 | P36871 | Cytoskeletal | Linker | 0.0077 | 0.0084 | 0.0079 | 0.0093 | 0.003 | 0.002 | 0.002 | 0.001 | **1.029** | **1.012** | **1.181** | 0.460 | 0.479 | 0.433 | 0.9607644 | 0.9716594 | 0.2198824 |
| PGM shared | 11 | B4DDQ8; P36871 | Cytoskeletal | Linker | 0.0077 | 0.0088 | 0.0077 | 0.0077 | 0.002 | 0.002 | 0.002 | 0.001 | **1.078** | **0.984** | **1.062** | 0.577 | 0.889 | 1.008 | 0.5331575 | 0.7519939 | 0.9596509 |
| PGM5 | 12 | Q15124 | Cytoskeletal | Linker | 0.0066 | 0.0089 | 0.0083 | 0.0089 | 0.002 | 0.002 | 0.002 | 0.002 | **1.329** | **1.235** | **1.362** | 0.430 | 0.394 | 0.390 | 3.996E-06 | 0.0002087 | 8.428E-06 |
| PLEC | 12 | Q15149 | Cytoskeletal | Linker | 0.0069 | 0.0071 | 0.0065 | 0.0065 | 0.002 | 0.001 | 0.001 | 0.001 | **1.074** | **0.997** | **1.007** | 0.338 | 0.324 | 0.309 | 0.4079151 | 0.5735138 | 0.9099323 |
| SGCA | 4 | Q16586 | Cytoskeletal | Linker | 0.0031 | 0.0033 | 0.0030 | 0.0033 | 0.001 | 0.001 | 0.001 | 0.001 | **1.034** | **0.951** | **1.020** | 0.329 | 0.320 | 0.275 | 0.5970742 | 0.6120681 | 0.6975218 |
| SNTA1 | 10 | Q13424 | Cytoskeletal | Linker | 0.0041 | 0.0041 | 0.0037 | 0.0035 | 0.001 | 0.001 | 0.001 | 0.000 | **0.994** | **0.898** | **0.791** | 0.350 | 0.350 | 0.215 | 0.2614761 | 0.3787495 | 0.0119198 |
| SPTAN1 | 11 | Q13813; A0A024R( | Cytoskeletal | Linker | 0.0229 | 0.0183 | 0.0173 | 0.0189 | 0.003 | 0.024 | 0.021 | 0.005 | **0.990** | **0.920** | **0.955** | 0.359 | 0.328 | 0.414 | 0.0609404 | 0.3610949 | 0.0477251 |
| TLN1 | 13 | Q9Y490 | Cytoskeletal | Linker | 0.0013 | 0.0016 | 0.0014 | 0.0015 | 0.000 | 0.001 | 0.000 | 0.000 | **1.326** | **1.177** | **1.281** | 0.706 | 0.433 | 0.376 | 1.372E-09 | 0.0005522 | 2.506E-07 |
| TLN2 | 6 | Q9Y4G6 | Cytoskeletal | Linker | 0.0025 | 0.0016 | 0.0019 | 0.0014 | 0.002 | 0.001 | 0.001 | 0.000 | **0.779** | **0.886** | **0.832** | 0.677 | 0.740 | 0.289 | 0.7919158 | 0.8654386 | 0.8097063 |
| TNS1 | 7 | A1L0S7 | Cytoskeletal | Linker | 0.0018 | 0.0015 | 0.0014 | 0.0013 | 0.000 | 0.001 | 0.000 | 0.000 | **0.947** | **0.826** | **0.853** | 0.445 | 0.373 | 0.273 | 0.1121914 | 0.0028626 | 0.0003534 |
| TUBA1A | 5 | Q71U36 | Cytoskeletal | Linker | 0.0129 | 0.0133 | 0.0130 | 0.0160 | 0.007 | 0.005 | 0.005 | 0.003 | **1.012** | **0.956** | **1.188** | 0.527 | 0.524 | 0.669 | 0.8652519 | 0.8109345 | 0.3314314 |
| TUBA4A | 4 | P68366 | Cytoskeletal | Linker | 0.0109 | 0.0155 | 0.0152 | 0.0164 | 0.003 | 0.004 | 0.003 | 0.003 | **1.421** | **1.423** | **1.545** | 0.526 | 0.466 | 0.496 | 0.0008801 | 0.0024216 | 0.0012769 |
| TUBB shared | 10 | Q5JP53; Q5SU16; I | Cytoskeletal | Linker | 0.0153 | 0.0193 | 0.0167 | 0.0163 | 0.004 | 0.007 | 0.005 | 0.006 | **1.132** | **1.144** | **1.077** | 0.905 | 0.858 | 0.755 | 0.8873062 | 0.8778246 | 0.1561932 |
| TUBB4B | 4 | P68371 | Cytoskeletal | Linker | 0.0102 | 0.0114 | 0.0096 | 0.0089 | 0.003 | 0.004 | 0.003 | 0.002 | **0.981** | **0.876** | **0.482** | 0.907 | 0.756 | 0.234 | 0.9981561 | 0.71272 | 0.1201088 |
| TUBB6 | 4 | Q9BUF5 | Cytoskeletal | Linker | 0.0161 | 0.0112 | 0.0148 | 0.0257 | 0.024 | 0.015 | 0.018 | 0.024 | **3.719** | **3.676** | **1.912** | 2.370 | 1.923 | 1.147 | 0.3833881 | 0.3723142 | 0.3959536 |
| VCL | 12 | V9HWK2 | Cytoskeletal | Linker | 0.0168 | 0.0221 | 0.0207 | 0.0230 | 0.003 | 0.005 | 0.003 | 0.004 | **1.303** | **1.258** | **1.375** | 0.396 | 0.308 | 0.354 | 6.334E-13 | 5.16E-08 | 2.869E-12 |
| | | | **Cytoskeletal** | **M-line** | 0 | 0 | 0 | 0 | | | | | **1.055** | **1.000** | **1.023** | | | | 0.4164784 | 0.0652456 | 0.615013 |
| KRT9 | 7 | P35527 | Cytoskeletal | M-line | 0.0013 | 0.0003 | 0.0003 | 0.0003 | 0.000 | 0.001 | 0.000 | 0.000 | **0.277** | **0.320** | **0.242** | 1.034 | 0.407 | 0.481 | 0.0716537 | 0.0095093 | 0.0040377 |
| KRT10 | 4 | P13645 | Cytoskeletal | M-line | 0.0006 | 0.0007 | 0.0004 | 0.0005 | 0.001 | 0.001 | 0.000 | 0.000 | **1.210** | **0.725** | **0.598** | 1.600 | 0.712 | 0.506 | 0.9885972 | 0.0608098 | 0.8340888 |
| KRT1 | 7 | H6VRG1 | Cytoskeletal | M-line | 0.0014 | 0.0010 | 0.0008 | 0.0007 | 0.001 | 0.003 | 0.001 | 0.000 | **1.056** | **0.742** | **0.554** | 18.340 | 11.638 | 1.104 | 0.3401134 | 0.8975948 | 0.7439402 |
| OBSCN | 13 | A6NGQ3 | Cytoskeletal | M-line | 0.0022 | 0.0029 | 0.0028 | 0.0031 | 0.000 | 0.001 | 0.000 | 0.001 | **1.265** | **1.222** | **1.348** | 0.388 | 0.390 | 0.479 | 2.357E-11 | 6.542E-07 | 4.287E-07 |
| SPTBN1 | 14 | B2ZZ89 | Cytoskeletal | M-line | 0.0091 | 0.0085 | 0.0083 | 0.0087 | 0.003 | 0.004 | 0.003 | 0.001 | **1.003** | **0.991** | **0.993** | 0.303 | 0.306 | 0.236 | 0.8959428 | 0.7270513 | 0.547784 |
| | | | **Cytoskeletal** | **Transport** | 0 | 0 | 0 | 0 | | | | | **1.167** | **1.042** | **1.152** | | | | 0.0078984 | 0.720439 | 0.4846351 |
| MYH10 | 6 | P35580 | Cytoskeletal | Transport | 0.0003 | 0.0005 | 0.0005 | 0.0005 | 0.000 | 0.001 | 0.000 | 0.000 | **1.853** | **1.796** | **1.458** | 2.408 | 1.180 | 0.839 | 0.0394285 | 0.1511183 | 0.4983401 |
| MYH9 | 12 | A0A024R1N1 | Cytoskeletal | Transport | 0.0029 | 0.0030 | 0.0028 | 0.0033 | 0.001 | 0.003 | 0.001 | 0.001 | **0.943** | **0.892** | **1.002** | 1.105 | 0.405 | 0.373 | 0.0768853 | 0.472529 | 0.4174012 |
| MYH11 | 13 | A0A024QZJ4 | Cytoskeletal | Transport | 0.0020 | 0.0028 | 0.0020 | 0.0028 | 0.001 | 0.005 | 0.001 | 0.002 | **1.384** | **1.043** | **1.399** | 5.427 | 1.350 | 2.924 | 0.0004472 | 0.719956 | 0.0111727 |
| MYO1C | 6 | O00159 | Cytoskeletal | Transport | 0.0009 | 0.0011 | 0.0010 | 0.0009 | 0.000 | 0.001 | 0.000 | 0.000 | **1.092** | **0.987** | **0.993** | 0.680 | 0.418 | 0.327 | 0.026593 | 0.3138053 | 0.6003188 |
| DYNC1H1 | 13 | Q14204 | Cytoskeletal | Transport | 0.0022 | 0.0024 | 0.0022 | 0.0022 | 0.001 | 0.001 | 0.001 | 0.000 | **1.105** | **1.086** | **1.051** | 0.459 | 0.433 | 0.353 | 0.2542032 | 0.5250363 | 0.5884577 |
| | | | **Cytoskeletal** | **Z-line** | 0 | 0 | 0 | 0 | | | | | **1.355** | **1.258** | **1.277** | | | | 0.0002578 | 0.0649625 | 0.0047193 |
| CAPZB | 5 | P47756 | Cytoskeletal | Z-line | 0.0034 | 0.0026 | 0.0027 | 0.0031 | 0.001 | 0.001 | 0.001 | 0.000 | **0.774** | **0.759** | **0.766** | 0.343 | 0.367 | 0.148 | 0.3898895 | 0.3870906 | 0.4789692 |
| NRAP | 14 | A0A0A0MRM2; A0 | Cytoskeletal | Z-line | 0.0043 | 0.0069 | 0.0031 | 0.0039 | 0.007 | 0.007 | 0.001 | 0.001 | **1.987** | **1.746** | **2.037** | 1.061 | 0.729 | 0.985 | 8.293E-10 | 0.0001829 | 7.383E-05 |
| PDLIM1 | 14 | O00151 | Cytoskeletal | Z-line | 0.0081 | 0.0117 | 0.0113 | 0.0110 | 0.002 | 0.002 | 0.002 | 0.002 | **1.382** | **1.389** | **1.385** | 0.823 | 0.692 | 0.628 | 0.0683539 | 0.0440042 | 0.028891 |
| PDLIM3 | 9 | Q53GG5 | Cytoskeletal | Z-line | 0.0052 | 0.0048 | 0.0034 | 0.0040 | 0.002 | 0.003 | 0.002 | 0.002 | **0.838** | **0.561** | **0.687** | 0.668 | 0.618 | 0.451 | 0.4659105 | 0.0377161 | 0.2818501 |
| PDLIM5 | 6 | B7Z8X5; B7Z481 | Cytoskeletal | Z-line | 0.0101 | 0.0133 | 0.0130 | 0.0129 | 0.003 | 0.003 | 0.003 | 0.002 | **1.305** | **1.311** | **1.259** | 0.447 | 0.440 | 0.370 | 0.0025149 | 0.016779 | 0.0331527 |
| | | | **Cytoskeletal** | | 0 | 0 | 0 | 0 | | | | | **1.123** | **0.964** | **1.143** | | | | 0.1787018 | 0.9846213 | 0.2726495 |
| EHD2 | 10 | Q9NZN4 | Cytoskeletal | Myoblast fusion | 0.0044 | 0.0035 | 0.0034 | 0.0032 | 0.002 | 0.002 | 0.001 | 0.001 | **0.826** | **0.835** | **0.706** | 0.772 | 0.565 | 0.362 | 0.2468352 | 0.1201691 | 0.1540527 |
| LDB3 | 3 | B4DUL3 | Cytoskeletal | | 0.0351 | 0.0201 | 0.0241 | 0.0144 | 0.006 | 0.009 | 0.012 | 0.009 | **0.587** | **0.613** | **0.439** | 0.284 | 0.346 | 0.231 | 0.0005428 | 0.0220862 | 0.0001834 |
| MAP4 | 12 | E7EVA0 | Cytoskeletal | Mibrotubules | 0.0031 | 0.0048 | 0.0044 | 0.0059 | 0.001 | 0.001 | 0.001 | 0.001 | **1.523** | **1.382** | **1.894** | 0.597 | 0.555 | 0.614 | 1.165E-10 | 9.156E-07 | 7.05E-15 |
| PPP2R1A | 6 | P30153 | Cytoskeletal | | 0.0035 | 0.0033 | 0.0031 | 0.0035 | 0.001 | 0.001 | 0.000 | 0.000 | **0.984** | **0.898** | **0.974** | 0.303 | 0.266 | 0.250 | 0.8396315 | 0.6192364 | 0.8158112 |
| SORBS1 | 9 | Q9BX66 | Cytoskeletal | | 0.0051 | 0.0057 | 0.0056 | 0.0059 | 0.001 | 0.002 | 0.001 | 0.001 | **1.148** | **1.024** | **0.985** | 0.664 | 0.585 | 0.532 | 0.6059724 | 0.5185732 | 0.4699571 |
| SORBS2 | 8 | H7BXR3 | Cytoskeletal | | 0.0121 | 0.0207 | 0.0175 | 0.0207 | 0.004 | 0.005 | 0.005 | 0.005 | **1.615** | **1.360** | **1.560** | 0.623 | 0.537 | 0.563 | 0.0023482 | 0.0331999 | 0.0472421 |
| | | | **Desmosome** | | | | | | | | | | **1.283** | **1.168** | **1.563** | | | | 7.074E-06 | 0.0787071 | 0.0005254 |
| | | | **Desmosome** | **Linker** | 0 | 0 | 0 | 0 | | | | | **0.940** | **0.940** | **0.902** | | | | 0.0019071 | 0.0256947 | 0.005486 |
| PKP2 | 14 | Q99959 | Desmosome | Linker | 0.0119 | 0.0118 | 0.0124 | 0.0119 | 0.001 | 0.002 | 0.002 | 0.002 | **1.035** | **1.040** | **1.024** | 0.223 | 0.222 | 0.193 | 0.603233 | 0.48122 | 0.7592371 |
| DSG2 | 4 | Q14126 | Desmosome | Linker | 0.0012 | 0.0007 | 0.0009 | 0.0009 | 0.000 | 0.000 | 0.000 | 0.000 | **0.515** | **0.689** | **0.727** | 0.281 | 0.327 | 0.368 | 3.708E-06 | 0.0266471 | 0.0341422 |
| DSP | 13 | P15924 | Desmosome | Linker | 0.0083 | 0.0069 | 0.0070 | 0.0068 | 0.001 | 0.001 | 0.001 | 0.001 | **0.850** | **0.845** | **0.790** | 0.222 | 0.244 | 0.191 | 1.085E-12 | 1.266E-11 | 1.529E-14 |
| JUP | 12 | A0A024R1X8 | Desmosome | Linker | 0.0072 | 0.0062 | 0.0064 | 0.0069 | 0.002 | 0.001 | 0.001 | 0.001 | **0.903** | **0.884** | **0.824** | 0.261 | 0.241 | 0.226 | 0.0008272 | 0.0082394 | 0.0006925 |
| DES | 13 | A5Z217 | Desmosome | Intermediate Filament | 0.2282 | 0.2888 | 0.2717 | 0.3433 | 0.105 | 0.057 | 0.049 | 0.074 | **1.313** | **1.191** | **1.614** | 0.684 | 0.616 | 0.820 | 4.614E-05 | 0.0295795 | 7.466E-10 |
| | | | **ECM** | | | | | | | | | | **2.227** | **1.912** | **2.817** | | | | 1.252E-37 | 3.148E-27 | 4.522E-38 |
| | | | **ECM** | **Collagen** | 0 | 0 | 0 | 0 | | | | | **2.408** | **2.049** | **3.043** | | | | 3.469E-22 | 2.241E-17 | 4.357E-27 |
| COL1A1 | 13 | P02452 | ECM | Collagen | 0.0587 | 0.1566 | 0.1452 | 0.2133 | 0.040 | 0.331 | 0.199 | 0.187 | **2.774** | **2.225** | **3.606** | 6.523 | 3.995 | 4.196 | 8.479E-20 | 1.135E-13 | 2.785E-24 |
| COL1A2 | 14 | P08123 | ECM | Collagen | 0.0457 | 0.1209 | 0.1212 | 0.1707 | 0.030 | 0.274 | 0.157 | 0.154 | **2.703** | **2.445** | **3.703** | 6.190 | 4.015 | 4.314 | 4.239E-34 | 6.569E-22 | 2.747E-39 |
| COL3A1 | 13 | P02461 | ECM | Collagen | 0.0829 | 0.2044 | 0.1699 | 0.2356 | 0.043 | 0.261 | 0.212 | 0.126 | **2.459** | **2.027** | **2.825** | 3.339 | 2.921 | 2.292 | 3.214E-19 | 2.049E-12 | 5.233E-24 |
| COL4A1 | 11 | P02462 | ECM | Collagen | 0.0220 | 0.0313 | 0.0312 | 0.0332 | 0.005 | 0.010 | 0.013 | 0.009 | **1.463** | **1.391** | **1.477** | 0.632 | 0.761 | 0.626 | 0.002608 | 0.0100787 | 0.0027325 |
| COL4A2 | 13 | A0A024RDW8 | ECM | Collagen | 0.0131 | 0.0181 | 0.0176 | 0.0188 | 0.003 | 0.006 | 0.007 | 0.006 | **1.354** | **1.406** | **1.483** | 0.680 | 0.807 | 0.741 | 4.756E-05 | 0.000162 | 3.09E-06 |
| COL5A1 | 11 | A0A087WXW9 | ECM | Collagen | 0.0056 | 0.0104 | 0.0087 | 0.0122 | 0.005 | 0.015 | 0.014 | 0.013 | **2.359** | **1.771** | **2.438** | 3.433 | 2.955 | 2.767 | 7.094E-07 | 0.0002306 | 1.091E-07 |
| COL5A2 | 13 | P05997 | ECM | Collagen | 0.0035 | 0.0081 | 0.0061 | 0.0092 | 0.002 | 0.009 | 0.008 | 0.007 | **2.395** | **1.853** | **2.922** | 2.846 | 2.587 | 2.536 | 5.318E-18 | 1.015E-11 | 4.225E-18 |
| COL6A1 | 14 | P12109 | ECM | Collagen | 0.0207 | 0.0331 | 0.0273 | 0.0397 | 0.005 | 0.018 | 0.024 | 0.012 | **1.534** | **1.355** | **1.885** | 0.997 | 1.327 | 0.751 | 1.001E-10 | 6.232E-09 | 7.628E-17 |
| COL6A2 | 11 | P12110 | ECM | Collagen | 0.0161 | 0.0225 | 0.0229 | 0.0302 | 0.005 | 0.013 | 0.021 | 0.008 | **1.423** | **1.429** | **1.777** | 0.925 | 1.358 | 0.740 | 1.667E-13 | 8.48E-10 | 4.76E-17 |
| COL6A3 | 5 | P12111 | ECM | Collagen | 0.0017 | 0.0012 | 0.0019 | 0.0022 | 0.002 | 0.001 | 0.001 | 0.001 | **2.659** | **3.130** | **3.114** | 1.365 | 1.968 | 1.651 | 0.7737296 | 0.403874 | 0.2794837 |
| COL15A1 | 5 | P39059 | ECM | Collagen | 0.0024 | 0.0029 | 0.0027 | 0.0035 | 0.001 | 0.001 | 0.001 | 0.001 | **1.247** | **1.216** | **1.438** | 0.575 | 0.577 | 0.634 | 0.0123862 | 0.1396522 | 0.0015898 |
| COL18A1 | 7 | P39060 | ECM | Collagen | 0.0010 | 0.0018 | 0.0016 | 0.0023 | 0.000 | 0.001 | 0.001 | 0.001 | **2.145** | **1.609** | **2.240** | 1.860 | 1.557 | 1.161 | 1.987E-05 | 0.0004647 | 3.257E-05 |
| | | | **ECM** | **Extracellular** | 0 | 0 | 0 | 0 | | | | | **1.794** | **1.670** | **2.084** | | | | 2.398E-12 | 1.837E-08 | 2.039E-10 |
| APOH | 7 | P02749 | ECM | Extracellular | 0.0041 | 0.0039 | 0.0045 | 0.0037 | 0.001 | 0.002 | 0.002 | 0.001 | **0.922** | **1.137** | **0.903** | 0.507 | 0.591 | 0.471 | 0.4509512 | 0.3348433 | 0.732792 |
| ASPN | 4 | Q6P528 | ECM | Extracellular | 0.0083 | 0.0086 | 0.0074 | 0.0032 | 0.005 | 0.005 | 0.006 | 0.003 | **3.452** | **4.451** | **7.712** | 5.665 | 12.570 | 8.958 | 0.052318 | 0.0424122 | 0.0983843 |
| FBN1 | 14 | P35555 | ECM | Extracellular | 0.0292 | 0.0446 | 0.0336 | 0.0478 | 0.018 | 0.015 | 0.031 | 0.010 | **1.546** | **1.201** | **1.606** | 1.079 | 1.258 | 1.071 | 9.257E-07 | 0.0003648 | 5.31E-11 |
| FN1 | 13 | A0A024R462 | ECM | Extracellular | 0.0016 | 0.0032 | 0.0028 | 0.0030 | 0.001 | 0.003 | 0.001 | 0.001 | **1.980** | **1.760** | **1.574** | 2.122 | 1.340 | 0.885 | 3.213E-17 | 2.807E-11 | 7.764E-09 |
| HSPG2 | 13 | A0A024RAB6 | ECM | Extracellular | 0.0105 | 0.0141 | 0.0128 | 0.0157 | 0.002 | 0.004 | 0.004 | 0.003 | **1.333** | **1.208** | **1.513** | 0.502 | 0.511 | 0.500 | 3.141E-14 | 2.427E-06 | 3.189E-13 |
| LUM | 8 | P51884 | ECM | Extracellular | 0.0085 | 0.0202 | 0.0171 | 0.0239 | 0.004 | 0.030 | 0.021 | 0.016 | **2.421** | **2.191** | **2.846** | 4.019 | 2.786 | 2.457 | 5.668E-14 | 7.114E-11 | 2.296E-15 |
| NID1 | 10 | P14543 | ECM | Extracellular | 0.0057 | 0.0059 | 0.0056 | 0.0060 | 0.001 | 0.001 | 0.001 | 0.001 | **1.097** | **1.047** | **1.092** | 0.410 | 0.405 | 0.393 | 0.771549 | 0.7888776 | 0.8855681 |
| NID2 | 13 | Q14112 | ECM | Extracellular | 0.0059 | 0.0071 | 0.0066 | 0.0066 | 0.002 | 0.002 | 0.001 | 0.001 | **1.237** | **1.111** | **1.136** | 0.473 | 0.384 | 0.365 | 0.0005263 | 0.1740789 | 0.1204349 |

| | | | | | | | | | | | | | | | | | | | | | |
|---|---|---|---|---|---|---|---|---|---|---|---|---|---|---|---|---|---|---|---|---|---|
| TNXB | 4 | E7EPZ9 | ECM | Extracellular | 0.0005 | 0.0015 | 0.0013 | 0.0014 | 0.000 | 0.001 | 0.001 | 0.001 | 2.429 | 2.627 | 2.659 | 2.911 | 2.614 | 2.218 | 0.0011589 | 0.0011751 | 0.0008347 |
| VCAN | 9 | Q59FG9 | ECM | Extracellular | 0.0010 | 0.0033 | 0.0029 | 0.0056 | 0.001 | 0.006 | 0.004 | 0.003 | 2.780 | 2.490 | 5.033 | 5.948 | 3.758 | 5.143 | 4.354E-11 | 3.331E-07 | 2.59E-09 |
| VTN | 6 | D9ZGG2 | ECM | Extracellular | 0.0035 | 0.0034 | 0.0038 | 0.0045 | 0.002 | 0.003 | 0.004 | 0.001 | 0.977 | 1.131 | 1.347 | 0.879 | 1.136 | 0.730 | 0.8121708 | 0.0310193 | 0.0653685 |
| | | | ECM | Intercalated disc | 0 | 0 | 0 | 0 | | | | | 0.911 | 0.998 | 0.843 | | | | 0.2659795 | 0.8283219 | 0.0549891 |
| CDH2 | 11 | A0A024RC42 | ECM | Intercalated disc | 0.0083 | 0.0093 | 0.0091 | 0.0077 | 0.001 | 0.002 | 0.002 | 0.001 | 1.042 | 1.045 | 0.971 | 0.448 | 0.449 | 0.284 | 0.4090605 | 0.7478163 | 0.2232243 |
| CDH13 | 6 | B7Z9B1 | ECM | Intercalated disc | 0.0070 | 0.0045 | 0.0063 | 0.0047 | 0.002 | 0.002 | 0.004 | 0.003 | 0.643 | 0.930 | 0.634 | 0.420 | 0.573 | 0.478 | 0.0116378 | 0.9470583 | 0.1314172 |
| | | | ECM | Linker | 0 | 0 | 0 | 0 | | | | | 1.348 | 1.132 | 1.647 | | | | 1.094E-07 | 0.0077765 | 0.0002028 |
| DAG1 | 5 | A0A024R2W4 | ECM | Linker | 0.0029 | 0.0034 | 0.0029 | 0.0042 | 0.001 | 0.001 | 0.001 | 0.001 | 1.162 | 0.991 | 1.100 | 0.562 | 0.506 | 0.557 | 0.4958436 | 0.7781833 | 0.5606885 |
| ITGB1 | 7 | P05556 | ECM | Linker | 0.0036 | 0.0041 | 0.0038 | 0.0036 | 0.001 | 0.001 | 0.001 | 0.001 | 1.189 | 1.047 | 1.005 | 0.503 | 0.397 | 0.292 | 0.0999447 | 0.8944246 | 0.8246111 |
| LAMA2 | 13 | P24043 | ECM | Linker | 0.0065 | 0.0065 | 0.0058 | 0.0062 | 0.001 | 0.001 | 0.001 | 0.001 | 0.990 | 0.875 | 0.924 | 0.341 | 0.275 | 0.281 | 0.5587899 | 0.0140025 | 0.1367408 |
| LAMA4 | 6 | H0UI49; A0A0A0M | ECM | Linker | 0.0008 | 0.0011 | 0.0010 | 0.0010 | 0.000 | 0.000 | 0.000 | 0.000 | 1.403 | 1.226 | 1.020 | 0.687 | 0.553 | 0.346 | 0.0217595 | 0.3203493 | 0.5366756 |
| LAMA5 | 10 | O15230 | ECM | Linker | 0.0016 | 0.0023 | 0.0021 | 0.0029 | 0.000 | 0.001 | 0.001 | 0.001 | 1.454 | 1.292 | 1.541 | 0.793 | 0.690 | 0.548 | 0.0049015 | 0.0316177 | 0.0087816 |
| LAMB1 | 12 | G3XAI2 | ECM | Linker | 0.0046 | 0.0041 | 0.0039 | 0.0036 | 0.001 | 0.001 | 0.001 | 0.000 | 0.895 | 0.848 | 0.717 | 0.311 | 0.325 | 0.217 | 0.2391443 | 0.063744 | 0.0005846 |
| LAMB2 | 13 | A0A024R319 | ECM | Linker | 0.0072 | 0.0093 | 0.0083 | 0.0106 | 0.002 | 0.002 | 0.002 | 0.002 | 1.318 | 1.171 | 1.467 | 0.528 | 0.442 | 0.525 | 2.582E-09 | 0.0026338 | 7.191E-09 |
| LAMC1 | 14 | P11047 | ECM | Linker | 0.0113 | 0.0129 | 0.0114 | 0.0116 | 0.003 | 0.003 | 0.004 | 0.002 | 1.159 | 1.034 | 1.122 | 0.401 | 0.355 | 0.331 | 0.0129912 | 0.394536 | 0.0888477 |
| PRELP | 11 | P51888 | ECM | Linker | 0.0028 | 0.0063 | 0.0050 | 0.0100 | 0.001 | 0.013 | 0.011 | 0.009 | 2.558 | 1.869 | 3.974 | 5.469 | 4.619 | 4.310 | 2.658E-11 | 1.775E-06 | 8.87E-14 |
| TAGLN | 8 | Q5U0D2 | ECM | Linker | 0.0015 | 0.0021 | 0.0015 | 0.0021 | 0.001 | 0.009 | 0.002 | 0.006 | 1.598 | 1.205 | 1.335 | 6.576 | 1.759 | 3.583 | 0.0032097 | 0.3451402 | 0.0202172 |
| TAGLN2 | 7 | P37802 | ECM | Linker | 0.0015 | 0.0015 | 0.0014 | 0.0015 | 0.001 | 0.001 | 0.001 | 0.001 | 1.153 | 1.078 | 0.947 | 2.035 | 0.802 | 0.572 | 0.1627569 | 0.0867577 | 0.2923843 |
| | | | ECM | | 0 | 0 | 0 | 0 | | | | | 2.371 | 1.863 | 2.961 | | | | 2.07E-06 | 2.73E-09 | 9.855E-10 |
| CILP | 6 | O75339 | ECM | | 0.0017 | 0.0030 | 0.0022 | 0.0036 | 0.002 | 0.002 | 0.005 | 0.004 | 1.570 | 1.258 | 1.681 | 2.094 | 3.116 | 3.153 | 0.050059 | 0.0416643 | 0.0025105 |
| BGN | 7 | A8K7E0 | ECM | | 0.0036 | 0.0108 | 0.0089 | 0.0155 | 0.002 | 0.023 | 0.018 | 0.008 | 3.018 | 2.667 | 4.629 | 6.731 | 5.483 | 4.170 | 1.064E-10 | 3.277E-10 | 1.722E-14 |
| DCN | 10 | P07585 | ECM | | 0.0083 | 0.0164 | 0.0125 | 0.0183 | 0.005 | 0.020 | 0.017 | 0.014 | 2.142 | 1.516 | 2.203 | 3.485 | 2.458 | 2.552 | 1.035E-05 | 0.0009539 | 2.642E-07 |
| ELN | 10 | G5E950; E7EN65 | ECM | | 0.0023 | 0.0014 | 0.0035 | 0.0029 | 0.003 | 0.002 | 0.005 | 0.003 | 0.616 | 1.503 | 1.210 | 1.387 | 3.774 | 2.433 | 0.0161266 | 0.0085046 | 0.2106094 |
| MFAP4 | 2 | K7ES70 | ECM | | 0.0003 | 0.0011 | 0.0005 | 0.0002 | 0.001 | 0.010 | 0.001 | 0.000 | 3.145 | 1.822 | 0.749 | 22.726 | 3.941 | 0.889 | 0.0511382 | 0.7624414 | 0.0628136 |
| THBS4 | 13 | P35443 | ECM | | 0.0017 | 0.0043 | 0.0032 | 0.0048 | 0.001 | 0.002 | 0.003 | 0.004 | 2.549 | 1.790 | 2.594 | 1.846 | 2.070 | 2.628 | 7.431E-18 | 2.146E-13 | 2.198E-19 |
| | | | ER | | 0 | 0 | 0 | 0 | | | | | 0.942 | 0.901 | 1.025 | | | | 0.1193283 | 0.5664964 | 0.0443593 |
| AGL | 13 | P35573 | ER | | 0.0024 | 0.0027 | 0.0024 | 0.0027 | 0.001 | 0.001 | 0.001 | 0.001 | 1.160 | 1.036 | 1.131 | 0.400 | 0.420 | 0.393 | 0.0001358 | 0.3382879 | 0.1531575 |
| CANX | 13 | P27824 | ER | | 0.0057 | 0.0073 | 0.0066 | 0.0071 | 0.001 | 0.002 | 0.002 | 0.001 | 1.229 | 1.128 | 1.136 | 0.697 | 0.555 | 0.366 | 0.0173914 | 0.2783407 | 0.0166681 |
| GPI | 13 | A0A0A0MTS2 | ER | | 0.0254 | 0.0209 | 0.0215 | 0.0253 | 0.007 | 0.004 | 0.004 | 0.005 | 0.879 | 0.863 | 1.024 | 0.396 | 0.365 | 0.394 | 0.1274631 | 0.0883134 | 0.4191199 |
| HSPA5 | 13 | V9HWB4 | ER | Chaperone | 0.0150 | 0.0144 | 0.0136 | 0.0157 | 0.003 | 0.003 | 0.003 | 0.002 | 0.967 | 0.938 | 1.027 | 0.274 | 0.284 | 0.255 | 0.761456 | 0.7919002 | 0.2890578 |
| MGST3 | 3 | Q5VV89 | ER | | 0.0209 | 0.0233 | 0.0207 | 0.0214 | 0.005 | 0.006 | 0.006 | 0.004 | 0.418 | 0.668 | 0.441 | 0.741 | 0.570 | 0.280 | 0.8658962 | 0.9308569 | 0.615264 |
| OGN | 10 | B4DI63 | ER | | 0.0039 | 0.0104 | 0.0083 | 0.0125 | 0.004 | 0.020 | 0.013 | 0.008 | 2.652 | 2.085 | 2.922 | 10.702 | 3.639 | 2.943 | 9.49E-06 | 0.0007625 | 0.0002268 |
| P4HB | 8 | A0A024R8S5 | ER | Lumen | 0.0033 | 0.0038 | 0.0035 | 0.0037 | 0.001 | 0.001 | 0.001 | 0.001 | 1.197 | 1.065 | 1.232 | 0.630 | 0.488 | 0.614 | 0.0033582 | 0.0899666 | 0.0007451 |
| PDIA3 | 10 | P30101 | ER | Chaperone | 0.0071 | 0.0063 | 0.0073 | 0.0113 | 0.003 | 0.004 | 0.003 | 0.005 | 1.158 | 1.049 | 1.187 | 0.578 | 0.472 | 0.603 | 0.33379 | 0.2845129 | 0.0758068 |
| PDIA6 | 3 | Q15084 | ER | Chaperone | 0.0027 | 0.0029 | 0.0029 | 0.0030 | 0.001 | 0.001 | 0.000 | 0.000 | 1.059 | 1.060 | 1.111 | 0.317 | 0.266 | 0.265 | 0.6385089 | 0.398774 | 0.2981793 |
| SERPINA1 | 13 | A0A024R6I7; P010 | ER | | 0.0588 | 0.0288 | 0.0372 | 0.0377 | 0.013 | 0.024 | 0.011 | 0.017 | 0.511 | 0.623 | 0.627 | 0.437 | 0.245 | 0.315 | 9.582E-08 | 2.036E-06 | 0.0009501 |
| VCP | 13 | V9HW80 | ER | | 0.0063 | 0.0064 | 0.0060 | 0.0065 | 0.002 | 0.001 | 0.001 | 0.001 | 1.033 | 1.007 | 1.016 | 0.339 | 0.346 | 0.292 | 0.722364 | 0.9435498 | 0.6834773 |
| CALR | 5 | V9HW88 | ER | | 0.0041 | 0.0043 | 0.0039 | 0.0043 | 0.001 | 0.001 | 0.001 | 0.001 | 1.034 | 0.954 | 1.063 | 0.350 | 0.357 | 0.402 | 0.9741698 | 0.4448783 | 0.5366962 |
| ARL6IP5 | 4 | A0A024R371 | ER | Membrane | 0.0056 | 0.0066 | 0.0056 | 0.0057 | 0.001 | 0.002 | 0.001 | 0.001 | 1.163 | 1.046 | 0.999 | 0.503 | 0.428 | 0.284 | 0.1898251 | 0.7060732 | 0.360399 |
| | | | Extracellular | | | | | | | | | | 1.353 | 1.473 | 1.476 | | | | 6.067E-06 | 1.727E-05 | 0.0002555 |
| | | | Extracellular | Secreted | 0 | 0 | 0 | 0 | | | | | 1.325 | 1.502 | 1.561 | | | | 0.0001107 | 0.0002007 | 0.0006113 |
| AMBP | 4 | P02760 | Extracellular | Secreted | 0.0014 | 0.0018 | 0.0017 | 0.0027 | 0.001 | 0.002 | 0.002 | 0.001 | 1.327 | 1.298 | 1.758 | 1.739 | 1.706 | 1.143 | 0.1475171 | 0.2221009 | 0.0435067 |
| APOA1 | 12 | P02647 | Extracellular | Secreted | 0.0132 | 0.0206 | 0.0232 | 0.0270 | 0.005 | 0.014 | 0.010 | 0.009 | 1.534 | 1.861 | 2.121 | 1.473 | 1.050 | 1.225 | 4.277E-05 | 2.06E-07 | 1.051E-07 |
| APOA4 | 6 | P06727 | Extracellular | Secreted | 0.0004 | 0.0012 | 0.0014 | 0.0011 | 0.000 | 0.001 | 0.001 | 0.001 | 3.529 | 4.762 | 3.119 | 3.318 | 4.031 | 4.060 | 7.892E-05 | 1.158E-05 | 1.9E-05 |
| C3 | 13 | V9HWA9 | Extracellular | Secreted | 0.0036 | 0.0039 | 0.0051 | 0.0049 | 0.002 | 0.004 | 0.002 | 0.002 | 1.177 | 1.474 | 1.331 | 1.634 | 1.073 | 1.038 | 0.004626 | 0.000224 | 0.0004608 |
| HP | 13 | P00738 | Extracellular | Secreted | 0.0080 | 0.0056 | 0.0033 | 0.0031 | 0.009 | 0.012 | 0.008 | 0.003 | 0.673 | 0.562 | 0.383 | 1.732 | 1.083 | 0.596 | 0.0105589 | 1.815E-05 | 2.792E-08 |
| HPX | 10 | P02790 | Extracellular | Secreted | 0.0075 | 0.0073 | 0.0091 | 0.0097 | 0.002 | 0.008 | 0.004 | 0.004 | 1.054 | 1.137 | 1.136 | 1.405 | 0.901 | 0.740 | 0.5070372 | 0.553873 | 0.8559068 |
| ORM1 | 6 | P02763 | Extracellular | Secreted | 0.0116 | 0.0050 | 0.0055 | 0.0048 | 0.005 | 0.009 | 0.003 | 0.004 | 0.462 | 0.465 | 0.334 | 0.999 | 0.431 | 0.278 | 0.0188036 | 0.016316 | 0.1356219 |
| PPIA | 8 | A8K486 | Extracellular | Secreted | 0.0222 | 0.0235 | 0.0225 | 0.0221 | 0.005 | 0.005 | 0.004 | 0.004 | 1.040 | 1.012 | 0.970 | 1.260 | 0.952 | 1.437 | 0.5348099 | 0.8070018 | 0.9023629 |
| TF | 13 | B4E1B2; P02787 | Extracellular | Secreted | 0.0119 | 0.0196 | 0.0241 | 0.0239 | 0.007 | 0.015 | 0.010 | 0.013 | 1.582 | 2.044 | 1.924 | 1.768 | 1.640 | 1.710 | 5.389E-10 | 2.628E-11 | 6.658E-14 |
| TGFBI | 8 | Q15582 | Extracellular | Secreted | 0.0012 | 0.0032 | 0.0028 | 0.0026 | 0.001 | 0.004 | 0.002 | 0.002 | 2.564 | 2.120 | 2.120 | 3.422 | 2.616 | 1.902 | 1.139E-16 | 2.864E-09 | 1.985E-08 |
| TGM2 | 12 | V9HWG3 | Extracellular | Secreted | 0.0210 | 0.0279 | 0.0288 | 0.0351 | 0.009 | 0.006 | 0.011 | 0.005 | 1.291 | 1.342 | 1.584 | 0.751 | 0.886 | 0.823 | 0.0001205 | 7.896E-05 | 2.934E-09 |
| TINAGL1 | 7 | Q9GZM7 | Extracellular | Secreted | 0.0036 | 0.0048 | 0.0038 | 0.0045 | 0.001 | 0.002 | 0.002 | 0.001 | 1.385 | 1.060 | 1.190 | 0.594 | 0.466 | 0.424 | 0.005714 | 0.4943969 | 0.0909154 |
| TTR | 6 | A6XGL1 | Extracellular | Secreted | 0.0009 | 0.0021 | 0.0022 | 0.0016 | 0.001 | 0.003 | 0.002 | 0.002 | 2.270 | 2.489 | 2.325 | 4.783 | 4.514 | 4.545 | 0.0145395 | 0.0265509 | 0.0046924 |
| AHSG | 4 | B7Z8Q2 | Extracellular | Secreted | 0.0038 | 0.0059 | 0.0063 | 0.0074 | 0.002 | 0.004 | 0.003 | 0.002 | 1.795 | 1.927 | 2.313 | 1.864 | 1.779 | 2.049 | 0.038886 | 0.0633977 | 0.0007912 |
| GC | 10 | V9HWI6 | Extracellular | | 0.0034 | 0.0027 | 0.0032 | 0.0031 | 0.001 | 0.002 | 0.001 | 0.001 | 0.745 | 0.822 | 0.858 | 0.757 | 0.401 | 0.331 | 0.0400027 | 0.0084274 | 0.0266111 |
| GSN | 14 | B7Z992; A0A0A0M | Extracellular | | 0.0187 | 0.0227 | 0.0233 | 0.0263 | 0.004 | 0.012 | 0.008 | 0.005 | 1.195 | 1.170 | 1.360 | 0.506 | 0.507 | 0.384 | 0.0011037 | 0.0039401 | 0.0001768 |
| IGHA1 | 6 | Q6N092 | Extracellular | | 0.0021 | 0.0026 | 0.0041 | 0.0030 | 0.002 | 0.006 | 0.003 | 0.003 | 1.390 | 1.912 | 1.304 | 2.408 | 6.460 | 1.880 | 0.0405577 | 0.0605858 | 0.0690261 |
| IGHG1 | 6 | A0A087WYC5; Q7Z | Extracellular | | 0.0203 | 0.0425 | 0.0451 | 0.0425 | 0.021 | 0.048 | 0.017 | 0.033 | 2.030 | 2.161 | 2.022 | 3.625 | 2.834 | 3.095 | 6.563E-05 | 4.754E-05 | 0.0003486 |
| IGHG2 | 3 | Q6N093 | Extracellular | | 0.0067 | 0.0087 | 0.0135 | 0.0082 | 0.013 | 0.012 | 0.007 | 0.012 | 1.308 | 1.768 | 1.027 | 1.996 | 2.363 | 1.722 | 0.8301676 | 0.2686002 | 0.9172937 |
| LGALS1 | 8 | P09382 | Extracellular | | 0.0423 | 0.0454 | 0.0393 | 0.0469 | 0.009 | 0.014 | 0.010 | 0.008 | 1.076 | 0.929 | 1.115 | 0.515 | 0.391 | 0.391 | 0.6899339 | 0.7622695 | 0.6348356 |
| LGALS3BP | 9 | Q08380 | Extracellular | | 0.0019 | 0.0021 | 0.0022 | 0.0017 | 0.002 | 0.001 | 0.001 | 0.000 | 1.048 | 1.070 | 0.919 | 1.607 | 1.640 | 0.903 | 0.941185 | 0.8346018 | 0.0080552 |
| SERPINA3 | 7 | B3KS79 | Extracellular | | 0.0111 | 0.0030 | 0.0034 | 0.0033 | 0.002 | 0.004 | 0.001 | 0.002 | 0.260 | 0.314 | 0.300 | 0.383 | 0.182 | 0.163 | 2.703E-10 | 2.59E-10 | 1.066E-08 |
| LGALS3 | 4 | Q59FR8; Q6NVH9 | Extracellular | | 0.0138 | 0.0091 | 0.0137 | 0.0138 | 0.004 | 0.003 | 0.006 | 0.005 | 0.719 | 0.872 | 0.715 | 1.139 | 0.644 | 0.386 | 0.7592148 | 0.8857381 | 0.611879 |
| | | | Fibroblast | | 0 | 0 | 0 | 0 | | | | | 2.637 | 1.608 | 7.835 | | | | 4.012E-11 | 5.992E-07 | 1.193E-20 |
| VIM | 13 | V9HWE1 | Fibroblast | | 0.0216 | 0.0274 | 0.0258 | 0.0317 | 0.010 | 0.017 | 0.015 | 0.009 | 1.302 | 1.125 | 1.466 | 1.008 | 1.055 | 0.842 | 0.0003182 | 0.0130348 | 9.517E-06 |
| POSTN | 13 | A0A024RDS2; B1AI | Fibroblast | | 0.0020 | 0.0106 | 0.0065 | 0.0302 | 0.002 | 0.026 | 0.015 | 0.033 | 6.094 | 3.521 | 14.524 | 16.777 | 9.094 | 27.616 | 4.494E-17 | 1.69E-11 | 1.417E-28 |
| | | | Golgi apparatus | | 0 | 0 | 0 | 0 | | | | | 1.145 | 0.822 | 1.362 | | | | 0.7886458 | 0.0634174 | 0.1793496 |
| PTGES2 | 4 | Q9H7Z7 | Golgi apparatus | | 0.0027 | 0.0022 | 0.0022 | 0.0020 | 0.001 | 0.001 | 0.001 | 0.001 | 0.832 | 0.833 | 0.732 | 0.312 | 0.347 | 0.262 | 0.0621838 | 0.0105155 | 0.0027684 |
| ARF3 | 3 | P61204 | Golgi apparatus | | 0.0043 | 0.0039 | 0.0040 | 0.0030 | 0.001 | 0.001 | 0.002 | 0.001 | 0.924 | 0.936 | 0.701 | 0.640 | 0.568 | 0.232 | 0.8431045 | 0.783825 | 0.1239217 |
| DBI | 6 | P07108 | Golgi apparatus | | 0.0097 | 0.0085 | 0.0066 | 0.0110 | 0.004 | 0.005 | 0.004 | 0.006 | 1.330 | 0.750 | 1.661 | 4.237 | 2.049 | 4.030 | 0.8622436 | 0.1702874 | 0.9460866 |
| | | | Lysosome | | 0 | 0 | 0 | 0 | | | | | 0.706 | 0.752 | 0.707 | | | | 0.0001159 | 0.0023865 | 9.978E-05 |
| ASAH1 | 6 | D3DSQ1 | Lysosome | | 0.0056 | 0.0053 | 0.0051 | 0.0052 | 0.002 | 0.002 | 0.002 | 0.002 | 1.009 | 0.894 | 0.847 | 0.586 | 0.460 | 0.381 | 0.759114 | 0.7118894 | 0.5755627 |
| CTSD | 13 | V9HWI3 | Lysosome | | 0.0254 | 0.0161 | 0.0171 | 0.0161 | 0.008 | 0.009 | 0.007 | 0.005 | 0.618 | 0.643 | 0.551 | 0.637 | 0.509 | 0.368 | 0.0060474 | 0.0064225 | 0.0005436 |
| PRDX6 | 11 | V9HWC7 | Lysosome | | 0.0219 | 0.0150 | 0.0162 | 0.0157 | 0.006 | 0.003 | 0.003 | 0.003 | 0.668 | 0.762 | 0.723 | 0.358 | 0.367 | 0.307 | 0.0005396 | 0.0033005 | 0.0073088 |
| PCYOX1 | 5 | Q9UHG3 | Lysosome | | 0.0029 | 0.0022 | 0.0025 | 0.0022 | 0.001 | 0.001 | 0.001 | 0.001 | 0.737 | 0.874 | 0.695 | 0.350 | 0.409 | 0.321 | 0.1684176 | 0.5887042 | 0.0695017 |
| PSAP | 5 | B1AVU8 | Lysosome | | 0.0207 | 0.0157 | 0.0171 | 0.0170 | 0.003 | 0.009 | 0.007 | 0.005 | 0.790 | 0.815 | 0.824 | 0.473 | 0.352 | 0.230 | 0.3761355 | 0.6492798 | 0.3930774 |
| TPP1 | 4 | D3DQU2 | Lysosome | | 0.0073 | 0.0046 | 0.0051 | 0.0077 | 0.006 | 0.004 | 0.004 | 0.004 | 0.550 | 0.711 | 0.704 | 0.468 | 0.604 | 0.513 | 0.1971324 | 0.6108665 | 0.5950281 |
| CTSB | 4 | A0A024R374 | Lysosome | | 0.0046 | 0.0028 | 0.0032 | 0.0026 | 0.001 | 0.001 | 0.001 | 0.001 | 0.617 | 0.688 | 0.552 | 0.349 | 0.309 | 0.255 | 0.0017828 | 0.0048576 | 2.763E-05 |
| | | | Mitochondria | | | | | | | | | | 0.842 | 0.747 | 0.815 | | | | 0.0027983 | 5.58E-09 | 5.131E-07 |
| | | | Mitochondria | HSP | 0 | 0 | 0 | 0 | | | | | 0.930 | 0.821 | 0.923 | | | | 0.3175977 | 3.517E-06 | 0.059223 |
| HSPE1 | 6 | P61604 | Mitochondria | HSP | 0.0186 | 0.0189 | 0.0162 | 0.0174 | 0.012 | 0.003 | 0.004 | 0.003 | 0.978 | 0.868 | 0.918 | 0.586 | 0.466 | 0.506 | 0.7505011 | 0.0365258 | 0.4644285 |
| HSPA9 | 13 | P38646 | Mitochondria | HSP | 0.0235 | 0.0233 | 0.0125 | 0.0150 | 0.009 | 0.005 | 0.002 | 0.002 | 0.883 | 0.797 | 0.957 | 0.295 | 0.257 | 0.229 | 0.3391979 | 0.0006199 | 0.0903587 |
| HSPD1 | 14 | P10809 | Mitochondria | HSP | 0.0215 | 0.0205 | 0.0174 | 0.0197 | 0.004 | 0.003 | 0.003 | 0.004 | 0.939 | 0.796 | 0.901 | 0.367 | 0.278 | 0.260 | 0.6915184 | 0.0056 | 0.4058404 |
| | | | Mitochondria | Inner-membrane space | 0 | 0 | 0 | 0 | | | | | 0.822 | 0.730 | 0.783 | | | | 5.163E-09 | 3.34E-22 | 2.501E-21 |
| AIFM1 | 13 | P10809 | Mitochondria | Inner-membrane space | 0.0215 | 0.0205 | 0.0174 | 0.0197 | 0.004 | 0.003 | 0.003 | 0.004 | 0.916 | 0.803 | 0.901 | 0.202 | 0.182 | 0.212 | 0.0197044 | 9.608E-07 | 0.0098271 |
| AK2 | 3 | P54819 | Mitochondria | Inner-membrane space | 0.0060 | 0.0055 | 0.0049 | 0.0051 | 0.001 | 0.001 | 0.001 | 0.001 | 0.890 | 0.792 | 0.838 | 0.374 | 0.348 | 0.210 | 0.8533022 | 0.6159146 | 0.3426465 |
| ACADVL | 13 | P49748 | Mitochondria | Inner-membrane | 0.0436 | 0.0310 | 0.0258 | 0.0269 | 0.009 | 0.009 | 0.008 | 0.007 | 0.698 | 0.596 | 0.660 | 0.262 | 0.224 | 0.250 | 1.106E-17 | 9.863E-28 | 2.339E-18 |
| ATP5A1 | 11 | V9HW26 | Mitochondria | Inner-membrane | 0.3170 | 0.2431 | 0.2319 | 0.2684 | 0.043 | 0.031 | 0.045 | 0.046 | 0.775 | 0.734 | 0.809 | 0.169 | 0.196 | 0.189 | 9.127E-05 | 1.37E-07 | 0.0001158 |
| ATP5B | 11 | V9HW31 | Mitochondria | Inner-membrane | 0.2531 | 0.1965 | 0.1808 | 0.2027 | 0.045 | 0.034 | 0.038 | 0.043 | 0.817 | 0.739 | 0.800 | 0.338 | 0.290 | 0.270 | 0.0737838 | 0.0065142 | 0.0071013 |
| ATP5C1 | 7 | B4DL14 | Mitochondria | Inner-membrane | 0.0566 | 0.0463 | 0.0426 | 0.0444 | 0.012 | 0.011 | 0.013 | 0.010 | 0.816 | 0.737 | 0.832 | 0.269 | 0.277 | 0.259 | 0.1206388 | 0.0058695 | 0.0145915 |
| ATP5F1 | 9 | A8K4W2 | Mitochondria | Inner-membrane | 0.0481 | 0.0368 | 0.0334 | 0.0366 | 0.009 | 0.009 | 0.010 | 0.008 | 0.806 | 0.732 | 0.686 | 0.557 | 0.469 | 0.347 | 0.447867 | 0.214634 | 0.2390233 |
| ATP5H | 11 | O75947 | Mitochondria | Inner-membrane | 0.0563 | 0.0467 | 0.0414 | 0.0452 | 0.007 | 0.006 | 0.007 | 0.007 | 0.802 | 0.731 | 0.738 | 0.348 | 0.208 | 0.197 | 0.306176 | 0.5100078 | 0.2384814 |
| ATP5I | 5 | P56385 | Mitochondria | Inner-membrane | 0.0416 | 0.0342 | 0.0321 | 0.0313 | 0.008 | 0.008 | 0.009 | 0.006 | 0.819 | 0.786 | 0.748 | 0.289 | 0.308 | 0.223 | 0.1102985 | 0.0126681 | 0.0057929 |
| ATP5J | 3 | Q6IB54 | Mitochondria | Inner-membrane | 0.0337 | 0.0245 | 0.0201 | 0.0238 | 0.011 | 0.007 | 0.007 | 0.006 | 0.721 | 0.603 | 0.692 | 0.329 | 0.302 | 0.286 | 0.0273632 | 0.0043641 | 0.0592255 |
| ATP5L | 6 | O75964 | Mitochondria | Inner-membrane | 0.0312 | 0.0266 | 0.0238 | 0.0263 | 0.010 | 0.008 | 0.009 | 0.008 | 1.108 | 0.801 | 0.928 | 0.674 | 0.495 | 0.425 | 0.7157946 | 0.236435 | 0.3803371 |
| ATP5O | 9 | P48047 | Mitochondria | Inner-membrane | 0.1238 | 0.0905 | 0.1026 | 0.0965 | 0.074 | 0.071 | 0.074 | 0.038 | 0.798 | 0.747 | 0.825 | 0.402 | 0.367 | 0.360 | 0.2711295 | 0.1809093 | 0.4481051 |
| ATPIF1 | 4 | Q9UII2 | Mitochondria | Inner-membrane | 0.0174 | 0.0199 | 0.0176 | 0.0182 | 0.002 | 0.004 | 0.003 | 0.006 | 1.193 | 0.999 | 1.039 | 0.408 | 0.494 | 0.540 | 0.3821553 | 0.827535 | 0.970537 |
| ADCK3 | 3 | Q8NI60 | Mitochondria | Inner-membrane | 0.0027 | 0.0016 | 0.0015 | 0.0019 | 0.001 | 0.001 | 0.001 | 0.001 | 0.503 | 0.534 | 0.736 | 0.470 | 0.381 | 0.356 | 0.0067509 | 0.0013252 | 0.0765111 |
| ATP | 13 | Q8TA92 | Mitochondria | Inner-membrane | 0.0059 | 0.0055 | 0.0049 | 0.0052 | 0.004 | 0.002 | 0.001 | 0.001 | 0.838 | 0.738 | 0.791 | 0.395 | 0.326 | 0.353 | 0.0107387 | 3.555E-05 | 0.000277 |
| BDH1 | 5 | Q02338 | Mitochondria | Inner-membrane | 0.0043 | 0.0037 | 0.0035 | 0.0036 | 0.001 | 0.001 | 0.001 | 0.001 | 0.812 | 0.792 | 0.815 | 0.284 | 0.294 | 0.271 | 0.0178027 | 0.0045581 | 0.0010459 |
| CHCHD3 | 10 | C9JRZ6 | Mitochondria | Inner-membrane | 0.0105 | 0.0097 | 0.0088 | 0.0083 | 0.001 | 0.002 | 0.002 | 0.002 | 1.013 | 0.848 | 0.875 | 0.289 | 0.273 | 0.289 | 0.958139 | 0.3983345 | 0.6487674 |
| COQ5 | 5 | B3GK62 | Mitochondria | Inner-membrane | 0.0019 | 0.0013 | 0.0015 | 0.0015 | 0.001 | 0.001 | 0.000 | 0.001 | 0.849 | 0.794 | 0.802 | 0.676 | 0.514 | 0.371 | 0.1452567 | 0.2661344 | 0.699761 |
| COX4I1 | 11 | P13073 | Mitochondria | Inner-membrane | 0.0606 | 0.0417 | 0.0380 | 0.0361 | 0.021 | 0.017 | 0.022 | 0.014 | 0.632 | 0.638 | 0.457 | 0.709 | 0.756 | 0.353 | 0.0382064 | 0.0838218 | 0.0044562 |
| COX5A | 8 | P20674 | Mitochondria | Inner-membrane | 0.0680 | 0.0505 | 0.0426 | 0.0497 | 0.017 | 0.017 | 0.019 | 0.019 | 1.005 | 0.631 | 0.719 | 4.260 | 1.874 | 4.163 | 0.7770151 | 0.1966116 | 0.1722386 |
| COX5B | 5 | P10606 | Mitochondria | Inner-membrane | 0.0573 | 0.0447 | 0.0383 | 0.0428 | 0.014 | 0.014 | 0.014 | 0.014 | 0.836 | 0.675 | 0.757 | 0.978 | 0.846 | 1.098 | 0.3851009 | 0.2302307 | 0.7248868 |
| COX6B1 | 8 | P14854 | Mitochondria | Inner-membrane | 0.0732 | 0.0536 | 0.0478 | 0.0501 | 0.014 | 0.015 | 0.011 | 0.015 | 0.751 | 0.611 | 0.713 | 0.870 | 0.781 | 0.496 | 0.48712 | 0.1911493 | 0.0939536 |
| COX6C | 5 | P09669 | Mitochondria | Inner-membrane | 0.0623 | 0.0478 | 0.0419 | 0.0472 | 0.012 | 0.014 | 0.015 | 0.015 | 1.077 | 0.685 | 0.789 | 1.790 | 0.967 | 0.421 | 0.8528857 | 0.9840918 | 0.2970161 |
| CPT2 | 5 | Q53G79 | Mitochondria | Inner-membrane | 0.0015 | 0.0015 | 0.0013 | 0.0014 | 0.001 | 0.000 | 0.000 | 0.001 | 1.027 | 0.875 | 0.920 | 1.084 | 0.977 | 1.139 | 0.7539314 | 0.5983597 | 0.3620414 |
| CRAT | 6 | P43155 | Mitochondria | Inner-membrane | 0.0043 | 0.0039 | 0.0036 | 0.0038 | 0.001 | 0.001 | 0.001 | 0.001 | 0.882 | 0.827 | 0.819 | 0.321 | 0.324 | 0.339 | 0.2657394 | 0.0412949 | 0.0516384 |
| CYB5R1 | 10 | Q9UHQ9 | Mitochondria | Inner-membrane | 0.0109 | 0.0109 | 0.0104 | 0.0115 | 0.003 | 0.003 | 0.003 | 0.002 | 1.046 | 0.976 | 0.957 | 0.612 | 0.516 | 0.489 | 0.4279368 | 0.8916872 | 0.9338854 |
| CYB5R3 | 9 | P00387 | Mitochondria | Inner-membrane | 0.0132 | 0.0157 | 0.0150 | 0.0157 | 0.004 | 0.005 | 0.005 | 0.004 | 1.121 | 1.111 | 1.097 | 0.595 | 0.576 | 0.460 | 0.0063379 | 0.2227313 | 0.2970242 |
| CYC1 | 7 | P08574 | Mitochondria | Inner-membrane | 0.0429 | 0.0341 | 0.0295 | 0.0333 | 0.011 | 0.008 | 0.009 | 0.007 | 0.898 | 0.719 | 0.775 | 0.476 | 0.489 | 0.384 | 0.3749134 | 0.105995 | 0.1036411 |
| CYCS | 7 | P99999 | Mitochondria | Inner-membrane | 0.0815 | 0.0827 | 0.0714 | 0.0847 | 0.021 | 0.019 | 0.018 | 0.020 | 1.128 | 0.728 | 1.042 | 1.068 | 0.501 | 0.326 | 0.8030532 | 0.2816398 | 0.262773 |
| CKMT2 | 13 | A0A024RAK5 | Mitochondria | Inner-membrane | 0.2222 | 0.1596 | 0.1472 | 0.1719 | 0.046 | 0.034 | 0.043 | 0.038 | 0.720 | 0.670 | 0.754 | 0.336 | 0.256 | 0.270 | 0.003918 | 0.0345344 | 0.1536176 |
| COQ9 | 4 | A0A024R6U3 | Mitochondria | Inner-membrane | 0.0086 | 0.0064 | 0.0056 | 0.0062 | 0.002 | 0.001 | 0.002 | 0.002 | 0.775 | 0.654 | 0.678 | 0.282 | 0.253 | 0.228 | 0.0930571 | 0.0079757 | 0.0330947 |
| DBT | 6 | P11182 | Mitochondria | Inner-membrane | 0.0032 | 0.0030 | 0.0027 | 0.0025 | 0.001 | 0.001 | 0.001 | 0.001 | 1.081 | 0.819 | 0.664 | 0.721 | 0.517 | 0.308 | 0.838096 | 0.4435674 | 0.0323189 |
| DLAT | 13 | A0A024R3D8 | Mitochondria | Inner-membrane | 0.0243 | 0.0260 | 0.0241 | 0.0237 | 0.004 | 0.003 | 0.003 | 0.004 | 1.110 | 1.044 | 0.983 | 0.439 | 0.403 | 0.341 | 0.1534126 | 0.7945029 | 0.883784 |
| ETFDH | 14 | A7UNU5 | Mitochondria | Inner-membrane | 0.0117 | 0.0091 | 0.0077 | 0.0079 | 0.003 | 0.002 | 0.002 | 0.002 | 0.781 | 0.650 | 0.687 | 0.267 | 0.239 | 0.234 | 5.405E-08 | 4.284E-14 | 1.855E-11 |
| GOT1 | 12 | P17174 | Mitochondria | Inner-membrane | 0.0504 | 0.0364 | 0.0344 | 0.0405 | 0.016 | 0.007 | 0.008 | 0.014 | 0.715 | 0.686 | 0.777 | 0.264 | 0.266 | 0.333 | 1.317E-07 | 1E-07 | 0.0076647 |
| GOT2 | 12 | A0A024R6W0 | Mitochondria | Inner-membrane | 0.0768 | 0.0732 | 0.0698 | 0.0804 | 0.019 | 0.014 | 0.013 | 0.017 | 0.984 | 0.875 | 0.978 | 0.376 | 0.334 | 0.341 | 0.7888995 | 0.327753 | 0.7737298 |
| HADHA | 13 | E9KL44 | Mitochondria | Inner-membrane | 0.0490 | 0.0386 | 0.0315 | 0.0343 | 0.014 | 0.011 | 0.008 | 0.007 | 0.808 | 0.650 | 0.657 | 0.319 | 0.241 | 0.219 | 6.439E-07 | 4.79E-16 | 1.273E-14 |
| IMMT | 4 | Q16891 | Mitochondria | Inner-membrane | 0.0041 | 0.0040 | 0.0033 | 0.0031 | 0.001 | 0.001 | 0.001 | 0.001 | 0.960 | 0.801 | 0.622 | 0.788 | 0.546 | 0.451 | 0.5749238 | 0.6183747 | 0.6756088 |
| IMMT shared | 13 | H7C463; Q16891 | Mitochondria | Inner-membrane | 0.0271 | 0.0250 | 0.0246 | 0.0254 | 0.006 | 0.007 | 0.005 | 0.005 | 0.983 | 0.916 | 0.918 | 0.276 | 0.280 | 0.261 | 0.8385009 | 0.3430479 | 0.0796264 |
| LDHD | 4 | Q86WU2 | Mitochondria | Inner-membrane | 0.0021 | 0.0018 | 0.0019 | 0.0020 | 0.001 | 0.000 | 0.001 | 0.001 | 0.842 | 0.906 | 0.941 | 0.387 | 0.440 | 0.401 | 0.1674414 | 0.3544714 | 0.8024957 |
| LETM1 | 6 | O95202 | Mitochondria | Inner-membrane | 0.0018 | 0.0017 | 0.0015 | 0.0015 | 0.000 | 0.001 | 0.000 | 0.000 | 0.941 | 0.845 | 0.785 | 0.340 | 0.338 | 0.262 | 0.304535 | 0.0407341 | 0.0495568 |
| LRPPRC | 10 | E5KNY5 | Mitochondria | Inner-membrane | 0.0026 | 0.0024 | 0.0021 | 0.0023 | 0.001 | 0.001 | 0.000 | 0.001 | 0.868 | 0.840 | 0.889 | 0.371 | 0.353 | 0.426 | 0.019114 | 0.0023679 | 0.0212268 |
| MTCH2 | 7 | Q9Y6C9 | Mitochondria | Inner-membrane | 0.0042 | 0.0039 | 0.0034 | 0.0036 | 0.001 | 0.001 | 0.001 | 0.001 | 0.917 | 0.803 | 0.823 | 0.345 | 0.304 | 0.278 | 0.478899 | 0.011966 | 0.1049812 |
| NDUFA10 | 11 | E7ESZ7 | Mitochondria | Inner-membrane | 0.0128 | 0.0096 | 0.0090 | 0.0088 | 0.003 | 0.003 | 0.002 | 0.002 | 0.773 | 0.708 | 0.697 | 0.549 | 0.387 | 0.249 | 0.0981426 | 0.0448651 | 0.0623252 |
| NDUFA11 | 4 | Q86Y39 | Mitochondria | Inner-membrane | 0.0051 | 0.0044 | 0.0041 | 0.0036 | 0.002 | 0.002 | 0.002 | 0.002 | 0.898 | 0.782 | 0.712 | 0.469 | 0.428 | 0.342 | 0.6114114 | 0.2818218 | 0.127091 |
| NDUFA12 | 4 | Q53HG1 | Mitochondria | Inner-membrane | 0.0164 | 0.0091 | 0.0091 | 0.0098 | 0.005 | 0.006 | 0.006 | 0.006 | 0.631 | 0.615 | 0.672 | 0.307 | 0.309 | 0.330 | 0.0249463 | 0.0289476 | 0.0484037 |
| NDUFA13 | 5 | Q9P0J0 | Mitochondria | Inner-membrane | 0.0130 | 0.0104 | 0.0094 | 0.0097 | 0.004 | 0.003 | 0.002 | 0.002 | 0.829 | 0.714 | 0.656 | 0.456 | 0.365 | 0.285 | 0.2950279 | 0.0876743 | 0.0752488 |
| NDUFA2 | 6 | O43678 | Mitochondria | Inner-membrane | 0.0148 | 0.0119 | 0.0109 | 0.0118 | 0.003 | 0.002 | 0.002 | 0.002 | 0.824 | 0.724 | 0.808 | 0.226 | 0.195 | 0.217 | 0.0238627 | 0.0061484 | 0.0789449 |
| NDUFA5 | 5 | A0A024R745 | Mitochondria | Inner-membrane | 0.0154 | 0.0116 | 0.0113 | 0.0099 | 0.004 | 0.003 | 0.002 | 0.003 | 0.791 | 0.727 | 0.637 | 0.224 | 0.194 | 0.242 | 0.0123857 | 0.00643 | 0.0013819 |
| NDUFA6 | 4 | P56556 | Mitochondria | Inner-membrane | 0.0108 | 0.0081 | 0.0073 | 0.0073 | 0.002 | 0.003 | 0.002 | 0.002 | 0.735 | 0.689 | 0.690 | 0.407 | 0.365 | 0.294 | 0.1043218 | 0.0407793 | 0.0153519 |
| NDUFA7 | 5 | Q32Q14 | Mitochondria | Inner-membrane | 0.0094 | 0.0073 | 0.0066 | 0.0066 | 0.003 | 0.002 | 0.002 | 0.002 | 0.790 | 0.700 | 0.634 | 0.422 | 0.373 | 0.376 | 0.0012826 | 0.0001139 | 0.0002148 |
| NDUFA8 | 9 | P51970 | Mitochondria | Inner-membrane | 0.0180 | 0.0139 | 0.0122 | 0.0114 | 0.004 | 0.003 | 0.003 | 0.002 | 0.858 | 0.710 | 0.617 | 0.564 | 0.462 | 0.228 | 0.3381215 | 0.075071 | 0.0057114 |
| NDUFA9 | 10 | Q16795 | Mitochondria | Inner-membrane | 0.0182 | 0.0142 | 0.0135 | 0.0123 | 0.006 | 0.004 | 0.003 | 0.004 | 0.709 | 0.714 | 0.593 | 0.362 | 0.320 | 0.288 | 0.0019821 | 0.0001374 | 1.928E-06 |
| NDUFB10 | 6 | A8K761 | Mitochondria | Inner-membrane | 0.0094 | 0.0075 | 0.0068 | 0.0067 | 0.003 | 0.002 | 0.002 | 0.002 | 0.823 | 0.714 | 0.645 | 0.336 | 0.289 | 0.245 | 0.1820806 | 0.0175498 | 0.0067934 |
| NDUFB4 | 6 | O95168 | Mitochondria | Inner-membrane | 0.0157 | 0.0127 | 0.0114 | 0.0127 | 0.004 | 0.004 | 0.004 | 0.003 | 0.840 | 0.707 | 0.765 | 0.280 | 0.228 | 0.243 | 0.1245665 | 0.0450305 | 0.1346498 |
| NDUFB5 | 5 | H0Y886 | Mitochondria | Inner-membrane | 0.0072 | 0.0058 | 0.0055 | 0.0041 | 0.003 | 0.002 | 0.002 | 0.002 | 0.769 | 0.723 | 0.575 | 0.477 | 0.353 | 0.242 | 0.3675341 | 0.0634056 | 0.0046142 |
| NDUFB6 | 5 | O95139 | Mitochondria | Inner-membrane | 0.0042 | 0.0034 | 0.0032 | 0.0030 | 0.002 | 0.002 | 0.002 | 0.001 | 0.868 | 0.808 | 0.717 | 2.070 | 1.406 | 2.149 | 0.3685849 | 0.1102159 | 0.0938135 |
| NDUFB9 | 6 | Q9Y6M9 | Mitochondria | Inner-membrane | 0.0167 | 0.0131 | 0.0126 | 0.0124 | 0.004 | 0.003 | 0.003 | 0.003 | 0.809 | 0.697 | 0.614 | 0.314 | 0.290 | 0.226 | 0.0316819 | 0.0107959 | 0.0035839 |
| NDUFS1 | 13 | P28331 | Mitochondria | Inner-membrane | 0.0271 | 0.0199 | 0.0192 | 0.0203 | 0.006 | 0.005 | 0.005 | 0.005 | 0.742 | 0.730 | 0.727 | 0.249 | 0.216 | 0.235 | 1.146E-07 | 4.251E-09 | 1.195E-08 |

| | | | | | | | | | | | | | | | | | | | | | |
|---|---|---|---|---|---|---|---|---|---|---|---|---|---|---|---|---|---|---|---|---|---|
| NDUFS2 | 11 | O75306 | Mitochondria | Inner-membrane | 0.0262 | 0.0207 | 0.0196 | 0.0188 | 0.005 | 0.004 | 0.004 | 0.004 | **0.767** | **0.667** | **0.629** | 0.508 | 0.328 | 0.218 | 0.0632208 | 0.0055368 | 0.0448176 |
| NDUFS3 | 13 | O75489 | Mitochondria | Inner-membrane | 0.0208 | 0.0158 | 0.0152 | 0.0155 | 0.005 | 0.004 | 0.003 | 0.004 | **0.759** | **0.697** | **0.721** | 0.528 | 0.552 | 0.502 | 0.0905809 | 0.0296344 | 0.0659126 |
| NDUFS5 | 7 | O43920 | Mitochondria | Inner-membrane | 0.0127 | 0.0106 | 0.0091 | 0.0078 | 0.003 | 0.003 | 0.002 | 0.002 | **0.849** | **0.710** | **0.494** | 0.413 | 0.333 | 0.215 | 0.0346253 | 0.0152685 | 0.0005485 |
| NDUFS6 | 7 | O75380 | Mitochondria | Inner-membrane | 0.0106 | 0.0083 | 0.0074 | 0.0077 | 0.003 | 0.003 | 0.003 | 0.002 | **0.798** | **0.699** | **0.707** | 0.750 | 0.519 | 0.733 | 0.188785 | 0.0422254 | 0.1031171 |
| NDUFS8 | 3 | O00217 | Mitochondria | Inner-membrane | 0.0106 | 0.0076 | 0.0079 | 0.0086 | 0.003 | 0.002 | 0.003 | 0.002 | **0.753** | **0.722** | **0.835** | 0.314 | 0.305 | 0.327 | 0.0161775 | 0.0150193 | 0.1014714 |
| NDUFV1 | 14 | P49821 | Mitochondria | Inner-membrane | 0.0225 | 0.0199 | 0.0166 | 0.0197 | 0.004 | 0.006 | 0.007 | 0.004 | **0.838** | **0.724** | **0.732** | 0.357 | 0.294 | 0.255 | 0.0138323 | 0.0001949 | 0.000183 |
| NDUFV2 | 8 | E7EPT4 | Mitochondria | Inner-membrane | 0.0183 | 0.0139 | 0.0128 | 0.0125 | 0.004 | 0.004 | 0.004 | 0.003 | **0.751** | **0.684** | **0.627** | 0.393 | 0.364 | 0.335 | 0.0269034 | 0.014207 | 0.008884 |
| NNT | 13 | A0A024R0C3 | Mitochondria | Inner-membrane | 0.0471 | 0.0424 | 0.0398 | 0.0452 | 0.008 | 0.005 | 0.007 | 0.010 | **0.883** | **0.843** | **0.954** | 0.204 | 0.204 | 0.261 | 0.475173 | 0.007797 | 0.8660994 |
| OPA1 | 13 | E5KLJ5 | Mitochondria | Inner-membrane | 0.0034 | 0.0036 | 0.0032 | 0.0036 | 0.000 | 0.000 | 0.001 | 0.000 | **1.022** | **0.961** | **1.084** | 0.312 | 0.317 | 0.329 | 0.5852555 | 0.9690665 | 0.2482277 |
| PHB | 10 | A8K401 | Mitochondria | Inner-membrane | 0.0136 | 0.0125 | 0.0118 | 0.0121 | 0.002 | 0.002 | 0.002 | 0.002 | **0.922** | **0.871** | **0.945** | 0.205 | 0.184 | 0.191 | 0.3267396 | 0.0087954 | 0.0494226 |
| PHB2 | 11 | Q99623 | Mitochondria | Inner-membrane | 0.0090 | 0.0088 | 0.0078 | 0.0090 | 0.002 | 0.002 | 0.002 | 0.002 | **0.971** | **0.863** | **0.923** | 0.466 | 0.427 | 0.397 | 0.8701677 | 0.2202051 | 0.7642184 |
| SDHA | 13 | A0A024QZ30 | Mitochondria | Inner-membrane | 0.0245 | 0.0204 | 0.0186 | 0.0216 | 0.004 | 0.004 | 0.004 | 0.004 | **0.877** | **0.791** | **0.876** | 0.292 | 0.284 | 0.269 | 0.0153879 | 4.493E-05 | 0.053789 |
| SDHB | 9 | P21912 | Mitochondria | Inner-membrane | 0.0204 | 0.0163 | 0.0157 | 0.0156 | 0.004 | 0.003 | 0.004 | 0.003 | **0.828** | **0.758** | **0.883** | 0.291 | 0.265 | 0.436 | 0.1827595 | 0.0781098 | 0.2061469 |
| SLC25A11 | 11 | Q02978 | Mitochondria | Inner-membrane | 0.0137 | 0.0108 | 0.0109 | 0.0117 | 0.004 | 0.005 | 0.004 | 0.003 | **0.781** | **0.789** | **0.675** | 0.604 | 0.489 | 0.349 | 0.5148901 | 0.285234 | 0.0235457 |
| SLC25A12 | 14 | B3KMV8 | Mitochondria | Inner-membrane | 0.0104 | 0.0094 | 0.0087 | 0.0089 | 0.001 | 0.001 | 0.001 | 0.002 | **0.897** | **0.840** | **0.867** | 0.290 | 0.292 | 0.312 | 0.1489234 | 0.0033002 | 0.0028801 |
| SLC25A20 | 6 | O43772 | Mitochondria | Inner-membrane | 0.0055 | 0.0039 | 0.0037 | 0.0037 | 0.002 | 0.002 | 0.001 | 0.001 | **0.795** | **0.712** | **0.683** | 1.657 | 1.511 | 0.805 | 0.74791 | 0.4956828 | 0.578301 |
| SLC25A3 | 11 | Q00325 | Mitochondria | Inner-membrane | 0.0658 | 0.0612 | 0.0550 | 0.0588 | 0.015 | 0.014 | 0.015 | 0.015 | **0.878** | **0.826** | **0.742** | 2.229 | 1.953 | 1.063 | 0.6095483 | 0.556771 | 0.0134812 |
| SLC25A4 | 5 | P12235 | Mitochondria | Inner-membrane | 0.2196 | 0.1304 | 0.1374 | 0.1467 | 0.087 | 0.067 | 0.051 | 0.047 | **0.506** | **0.604** | **0.647** | 0.402 | 0.354 | 0.365 | 0.1512388 | 0.1745714 | 0.1426749 |
| SLC25A5 | 4 | P05141 | Mitochondria | Inner-membrane | 0.0067 | 0.0092 | 0.0087 | 0.0086 | 0.002 | 0.004 | 0.003 | 0.003 | **1.403** | **1.310** | **1.120** | 0.831 | 0.574 | 0.454 | 0.0423648 | 0.4248141 | 0.4927485 |
| TUFM | 13 | P49411 | Mitochondria | Inner-membrane | 0.0223 | 0.0196 | 0.0184 | 0.0212 | 0.004 | 0.002 | 0.004 | 0.003 | **0.855** | **0.824** | **0.883** | 0.261 | 0.274 | 0.235 | 0.1675025 | 0.0278689 | 0.1337571 |
| UQCR10 | 5 | Q9UDW1 | Mitochondria | Inner-membrane | 0.0070 | 0.0051 | 0.0054 | 0.0045 | 0.004 | 0.004 | 0.003 | 0.003 | **0.893** | **0.981** | **0.650** | 4.711 | 4.764 | 1.048 | 0.7343627 | 0.8106882 | 0.7821694 |
| UQCRB | 6 | P14927 | Mitochondria | Inner-membrane | 0.0337 | 0.0271 | 0.0236 | 0.0249 | 0.006 | 0.005 | 0.006 | 0.004 | **0.775** | **0.691** | **0.743** | 0.271 | 0.245 | 0.221 | 0.1678403 | 0.0794116 | 0.111693 |
| UQCRC1 | 12 | P31930 | Mitochondria | Inner-membrane | 0.0656 | 0.0478 | 0.0426 | 0.0486 | 0.010 | 0.008 | 0.010 | 0.009 | **0.720** | **0.643** | **0.710** | 0.250 | 0.200 | 0.198 | 0.0012589 | 6.331E-06 | 0.0002133 |
| UQCRC2 | 13 | P22695 | Mitochondria | Inner-membrane | 0.0814 | 0.0604 | 0.0550 | 0.0496 | 0.023 | 0.018 | 0.014 | 0.011 | **0.777** | **0.651** | **0.704** | 0.268 | 0.236 | 0.199 | 0.0014561 | 1.3E-06 | 5.493E-08 |
| UQCRFS1 | 10 | P47985 | Mitochondria | Inner-membrane | 0.0431 | 0.0309 | 0.0290 | 0.0289 | 0.010 | 0.008 | 0.007 | 0.006 | **0.740** | **0.670** | **0.742** | 0.234 | 0.227 | 0.200 | 2.556E-06 | 2.183E-08 | 1.262E-07 |
| UQCRH | 4 | P07919 | Mitochondria | Inner-membrane | 0.0270 | 0.0227 | 0.0183 | 0.0191 | 0.006 | 0.005 | 0.006 | 0.004 | **0.764** | **0.644** | **0.705** | 0.236 | 0.222 | 0.194 | 0.0124697 | 3.743E-05 | 0.0001571 |
| UQCRQ | 5 | O14949 | Mitochondria | Inner-membrane | 0.0202 | 0.0160 | 0.0140 | 0.0161 | 0.006 | 0.004 | 0.005 | 0.003 | **0.808** | **0.724** | **0.801** | 0.316 | 0.309 | 0.281 | 0.2479474 | 0.0616622 | 0.4132407 |
| | | | **Mitochondria** | **Matrix** | 0 | 0 | 0 | 0 | | | | | **0.872** | **0.780** | **0.871** | | | | 0.0009541 | 5.996E-09 | 1.118E-05 |
| ACAA2 | 13 | B2RB23 | Mitochondria | Matrix | 0.0350 | 0.0255 | 0.0206 | 0.0223 | 0.011 | 0.007 | 0.007 | 0.005 | **0.721** | **0.585** | **0.658** | 0.417 | 0.414 | 0.358 | 0.0001047 | 2.438E-07 | 5.539E-05 |
| ACADM | 12 | Q5T4U5 | Mitochondria | Matrix | 0.0300 | 0.0324 | 0.0302 | 0.0319 | 0.007 | 0.008 | 0.006 | 0.008 | **1.093** | **0.972** | **1.013** | 0.785 | 0.646 | 0.563 | 0.5277891 | 0.4506413 | 0.906547 |
| ACADS | 5 | D4QEZ8 | Mitochondria | Matrix | 0.0048 | 0.0044 | 0.0033 | 0.0037 | 0.002 | 0.002 | 0.002 | 0.002 | **0.881** | **0.731** | **0.650** | 0.515 | 0.410 | 0.461 | 0.264657 | 0.0717425 | 0.1131336 |
| ACADSB | 5 | P45954 | Mitochondria | Matrix | 0.0024 | 0.0028 | 0.0026 | 0.0032 | 0.001 | 0.001 | 0.001 | 0.000 | **1.149** | **1.125** | **1.320** | 0.441 | 0.352 | 0.386 | 0.109582 | 0.2341567 | 0.0032394 |
| ACAT1 | 12 | P24752 | Mitochondria | Matrix | 0.0622 | 0.0525 | 0.0518 | 0.0548 | 0.016 | 0.010 | 0.010 | 0.010 | **0.880** | **0.833** | **0.863** | 0.383 | 0.349 | 0.311 | 0.2866329 | 0.0511987 | 0.0421388 |
| ACSS1 | 6 | Q9NUB1 | Mitochondria | Matrix | 0.0029 | 0.0025 | 0.0023 | 0.0018 | 0.001 | 0.001 | 0.001 | 0.001 | **0.880** | **0.750** | **0.658** | 0.321 | 0.341 | 0.224 | 0.1430816 | 0.0009523 | 1.356E-05 |
| ACO2 | 13 | B4DLY4; Q99798 | Mitochondria | Matrix | 0.0912 | 0.0856 | 0.0772 | 0.0833 | 0.021 | 0.014 | 0.014 | 0.017 | **0.947** | **0.821** | **0.901** | 0.278 | 0.254 | 0.274 | 0.375938 | 0.0059668 | 0.3009513 |
| ACOT9 | 8 | Q9Y305 | Mitochondria | Matrix | 0.0043 | 0.0044 | 0.0040 | 0.0048 | 0.001 | 0.001 | 0.001 | 0.001 | **1.085** | **0.959** | **1.106** | 0.336 | 0.291 | 0.316 | 0.4127823 | 0.0688413 | 0.2911257 |
| ALDH5A1 | 6 | X5D299 | Mitochondria | Matrix | 0.0032 | 0.0030 | 0.0025 | 0.0025 | 0.001 | 0.000 | 0.001 | 0.001 | **0.912** | **0.787** | **0.772** | 0.237 | 0.259 | 0.225 | 0.1656733 | 4.546E-06 | 0.0001192 |
| APOA1BP | 5 | B4DP80 | Mitochondria | Matrix | 0.0035 | 0.0028 | 0.0028 | 0.0033 | 0.001 | 0.001 | 0.001 | 0.001 | **0.800** | **0.766** | **0.942** | 0.376 | 0.419 | 0.393 | 0.1340025 | 0.1938632 | 0.7245931 |
| ADSSL1 | 6 | Q8N142 | Mitochondria | Matrix | 0.0014 | 0.0011 | 0.0011 | 0.0013 | 0.001 | 0.000 | 0.000 | 0.000 | **0.749** | **0.788** | **0.796** | 0.557 | 0.588 | 0.622 | 0.0419155 | 0.0596953 | 0.1315756 |
| AK3 | 9 | Q7Z4Y4 | Mitochondria | Matrix | 0.0125 | 0.0122 | 0.0111 | 0.0134 | 0.002 | 0.002 | 0.002 | 0.002 | **0.948** | **0.875** | **1.019** | 0.241 | 0.229 | 0.217 | 0.112745 | 0.004502 | 0.6867514 |
| AK4 | 9 | P27144 | Mitochondria | Matrix | 0.0166 | 0.0180 | 0.0166 | 0.0190 | 0.007 | 0.021 | 0.024 | 0.005 | **1.069** | **1.004** | **1.120** | 1.065 | 0.972 | 0.906 | 0.6224631 | 0.7664359 | 0.7275097 |
| ALDH2 | 12 | P05091 | Mitochondria | Matrix | 0.0124 | 0.0104 | 0.0110 | 0.0102 | 0.003 | 0.002 | 0.002 | 0.002 | **0.826** | **0.860** | **0.856** | 0.264 | 0.282 | 0.263 | 0.0007992 | 0.1072901 | 0.0267224 |
| ALDH4A1 | 11 | A0A024RAD8 | Mitochondria | Matrix | 0.0039 | 0.0042 | 0.0043 | 0.0055 | 0.001 | 0.001 | 0.001 | 0.001 | **1.122** | **1.145** | **1.473** | 0.644 | 0.712 | 0.470 | 0.0431705 | 0.0523732 | 0.0429492 |
| ALDH6A1 | 11 | Q02252 | Mitochondria | Matrix | 0.0051 | 0.0046 | 0.0051 | 0.0050 | 0.002 | 0.001 | 0.001 | 0.002 | **0.939** | **0.963** | **0.849** | 0.418 | 0.474 | 0.406 | 0.0821217 | 0.0584261 | 0.0360815 |
| ALDH7A1 | 5 | P49419 | Mitochondria | Matrix | 0.0035 | 0.0027 | 0.0028 | 0.0026 | 0.001 | 0.001 | 0.001 | 0.001 | **0.599** | **0.724** | **0.632** | 0.305 | 0.339 | 0.278 | 0.0114968 | 0.1248711 | 0.0203865 |
| ALDOC | 11 | A0A024QZ64 | Mitochondria | Matrix | 0.0230 | 0.0168 | 0.0162 | 0.0169 | 0.011 | 0.004 | 0.005 | 0.007 | **0.675** | **0.706** | **0.715** | 0.458 | 0.333 | 0.280 | 0.0165645 | 0.3451697 | 0.4275706 |
| CS | 6 | O75390 | Mitochondria | Matrix | 0.0552 | 0.0533 | 0.0458 | 0.0550 | 0.015 | 0.008 | 0.010 | 0.013 | **0.962** | **0.829** | **1.047** | 0.291 | 0.274 | 0.377 | 0.916654 | 0.3335117 | 0.9050863 |
| DECR1 | 12 | A0A024R9D7 | Mitochondria | Matrix | 0.0656 | 0.0409 | 0.0378 | 0.0423 | 0.016 | 0.005 | 0.010 | 0.009 | **0.653** | **0.579** | **0.598** | 0.275 | 0.261 | 0.190 | 0.0034663 | 0.0003614 | 3.825E-05 |
| DLST | 9 | A0A024R6C8 | Mitochondria | Matrix | 0.0282 | 0.0281 | 0.0244 | 0.0284 | 0.004 | 0.004 | 0.004 | 0.006 | **0.984** | **0.908** | **1.061** | 1.005 | 0.647 | 1.033 | 0.4913992 | 0.4996627 | 0.4848751 |
| ECHS1 | 15 | P30084 | Mitochondria | Matrix | 0.0202 | 0.0173 | 0.0170 | 0.0184 | 0.004 | 0.003 | 0.004 | 0.004 | **0.843** | **0.837** | **0.829** | 0.571 | 0.566 | 0.313 | 0.8107689 | 0.8332932 | 0.6066028 |
| ETFA | 13 | P13804 | Mitochondria | Matrix | 0.1358 | 0.1290 | 0.1088 | 0.1148 | 0.024 | 0.030 | 0.023 | 0.019 | **0.806** | **0.694** | **0.783** | 0.249 | 0.248 | 0.230 | 0.0983676 | 0.021045 | 0.1116644 |
| ETFB | 12 | P38117 | Mitochondria | Matrix | 0.0382 | 0.0281 | 0.0251 | 0.0300 | 0.007 | 0.005 | 0.006 | 0.006 | **0.760** | **0.653** | **0.815** | 0.266 | 0.267 | 0.282 | 0.0440502 | 0.0057104 | 0.1823372 |
| ECI1 | 4 | P42126 | Mitochondria | Matrix | 0.0100 | 0.0101 | 0.0084 | 0.0087 | 0.002 | 0.002 | 0.002 | 0.002 | **1.029** | **0.877** | **0.832** | 0.763 | 0.439 | 0.262 | 0.5752142 | 0.827201 | 0.5917418 |
| GLUD1 | 12 | E9KL48 | Mitochondria | Matrix | 0.0075 | 0.0088 | 0.0079 | 0.0087 | 0.001 | 0.002 | 0.001 | 0.002 | **1.171** | **1.054** | **1.157** | 0.431 | 0.384 | 0.453 | 0.0053702 | 0.6294847 | 0.009051 |
| HIBADH | 6 | A0A024RA75 | Mitochondria | Matrix | 0.0047 | 0.0034 | 0.0028 | 0.0026 | 0.002 | 0.001 | 0.001 | 0.001 | **0.719** | **0.627** | **0.606** | 0.630 | 0.482 | 0.391 | 0.0332955 | 0.0012824 | 0.000201 |
| FH | 13 | P07954 | Mitochondria | Matrix | 0.0276 | 0.0263 | 0.0224 | 0.0261 | 0.005 | 0.004 | 0.003 | 0.004 | **0.967** | **0.811** | **0.955** | 0.343 | 0.287 | 0.347 | 0.3755662 | 0.00313 | 0.5445209 |
| HADH | 8 | A0A0A0MSE2 | Mitochondria | Matrix | 0.0300 | 0.0263 | 0.0231 | 0.0255 | 0.005 | 0.005 | 0.004 | 0.005 | **0.910** | **0.764** | **0.809** | 0.921 | 0.848 | 0.682 | 0.634222 | 0.1425457 | 0.0460996 |
| HAGH | 4 | Q16775 | Mitochondria | Matrix | 0.0024 | 0.0025 | 0.0026 | 0.0026 | 0.000 | 0.001 | 0.000 | 0.000 | **1.093** | **1.111** | **1.231** | 0.380 | 0.277 | 0.249 | 0.9854976 | 0.4455916 | 0.5078929 |
| IDH2 | 13 | P48735 | Mitochondria | Matrix | 0.1328 | 0.0880 | 0.0841 | 0.0942 | 0.035 | 0.019 | 0.024 | 0.022 | **0.680** | **0.613** | **0.762** | 0.324 | 0.311 | 0.333 | 1.52E-05 | 3.081E-06 | 0.0009731 |
| IDH3A | 10 | P50213 | Mitochondria | Matrix | 0.0139 | 0.0129 | 0.0116 | 0.0132 | 0.004 | 0.002 | 0.002 | 0.002 | **0.977** | **0.906** | **0.912** | 0.459 | 0.406 | 0.385 | 0.6009733 | 0.0929873 | 0.2415991 |
| IDH3B | 7 | A0A087WZN1 | Mitochondria | Matrix | 0.0035 | 0.0039 | 0.0031 | 0.0038 | 0.002 | 0.001 | 0.001 | 0.001 | **1.005** | **0.896** | **1.101** | 0.965 | 0.689 | 0.802 | 0.7305691 | 0.9038955 | 0.4525233 |
| LONP1 | 9 | B3KU28 | Mitochondria | Matrix | 0.0066 | 0.0047 | 0.0061 | 0.0083 | 0.010 | 0.008 | 0.006 | 0.005 | **0.990** | **0.887** | **1.015** | 0.480 | 0.489 | 0.576 | 0.456277 | 0.4604341 | 0.7316009 |
| MCCC1 | 5 | Q96RQ3 | Mitochondria | Matrix | 0.0022 | 0.0018 | 0.0018 | 0.0021 | 0.001 | 0.000 | 0.000 | 0.000 | **0.758** | **0.789** | **0.900** | 0.415 | 0.434 | 0.437 | 0.2232632 | 0.19722 | 0.241155 |
| MCCC2 | 9 | Q9HCC0 | Mitochondria | Matrix | 0.0047 | 0.0038 | 0.0035 | 0.0033 | 0.001 | 0.001 | 0.001 | 0.001 | **0.779** | **0.759** | **0.726** | 0.295 | 0.293 | 0.267 | 0.0072594 | 0.0066936 | 0.0007068 |
| MDH2 | 13 | P40926 | Mitochondria | Matrix | 0.1273 | 0.0993 | 0.0843 | 0.0957 | 0.034 | 0.014 | 0.024 | 0.014 | **0.768** | **0.662** | **0.759** | 0.216 | 0.224 | 0.214 | 0.0009475 | 3.373E-06 | 0.000453 |
| ME2 | 4 | P23368 | Mitochondria | Matrix | 0.0017 | 0.0013 | 0.0013 | 0.0013 | 0.001 | 0.001 | 0.001 | 0.001 | **0.960** | **1.064** | **0.960** | 0.666 | 0.640 | 0.538 | 0.830792 | 0.891452 | 0.6900702 |
| MLYCD | 6 | O95822 | Mitochondria | Matrix | 0.0022 | 0.0018 | 0.0016 | 0.0017 | 0.000 | 0.001 | 0.001 | 0.000 | **0.834** | **0.706** | **0.764** | 0.357 | 0.335 | 0.285 | 0.3409847 | 0.0388414 | 0.1879175 |
| OGDH | 12 | B4DH65; Q02218; | Mitochondria | Matrix | 0.0171 | 0.0158 | 0.0146 | 0.0165 | 0.003 | 0.003 | 0.003 | 0.004 | **0.969** | **0.869** | **0.970** | 0.291 | 0.291 | 0.299 | 0.878763 | 0.0198539 | 0.5599365 |
| OXCT1 | 10 | P55809 | Mitochondria | Matrix | 0.0193 | 0.0275 | 0.0232 | 0.0259 | 0.004 | 0.005 | 0.005 | 0.005 | **1.387** | **1.150** | **1.370** | 0.507 | 0.431 | 0.397 | 0.0001255 | 0.0320685 | 0.0445424 |
| PDHA1 | 11 | P08559 | Mitochondria | Matrix | 0.0291 | 0.0310 | 0.0274 | 0.0275 | 0.005 | 0.006 | 0.006 | 0.005 | **1.063** | **0.909** | **0.907** | 0.289 | 0.298 | 0.236 | 0.5887597 | 0.4529261 | 0.289547 |
| PDHB | 5 | P11177 | Mitochondria | Matrix | 0.0258 | 0.0247 | 0.0234 | 0.0246 | 0.005 | 0.005 | 0.004 | 0.003 | **0.950** | **0.905** | **0.908** | 0.340 | 0.338 | 0.231 | 0.7336952 | 0.2508115 | 0.5264821 |
| PDHX | 6 | B2R673 | Mitochondria | Matrix | 0.0058 | 0.0069 | 0.0062 | 0.0060 | 0.001 | 0.001 | 0.001 | 0.001 | **1.199** | **1.070** | **1.032** | 0.290 | 0.235 | 0.256 | 0.0035672 | 0.3767275 | 0.2404378 |
| PPIF | 6 | A0A024QZS4 | Mitochondria | Matrix | 0.0088 | 0.0089 | 0.0075 | 0.0072 | 0.002 | 0.002 | 0.002 | 0.001 | **0.979** | **0.832** | **0.793** | 0.274 | 0.248 | 0.216 | 0.588182 | 0.2999655 | 0.2365715 |
| PRDX3 | 7 | P30048 | Mitochondria | Matrix | 0.0424 | 0.0368 | 0.0331 | 0.0353 | 0.008 | 0.005 | 0.006 | 0.006 | **0.828** | **0.774** | **0.905** | 0.267 | 0.251 | 0.283 | 0.261675 | 0.0510925 | 0.2556965 |
| PRDX5 | 7 | P30044 | Mitochondria | Matrix | 0.0247 | 0.0194 | 0.0200 | 0.0219 | 0.007 | 0.003 | 0.003 | 0.004 | **0.761** | **0.795** | **0.863** | 0.384 | 0.382 | 0.335 | 0.1248296 | 0.1335951 | 0.1486835 |
| QDPR | 4 | P09417 | Mitochondria | Matrix | 0.0039 | 0.0032 | 0.0030 | 0.0030 | 0.001 | 0.001 | 0.001 | 0.000 | **0.829** | **0.777** | **0.777** | 0.269 | 0.219 | 0.195 | 0.0040025 | 7.09E-05 | 3.056E-05 |
| SUCLA2 | 12 | Q6N0B1; E5KS55 | Mitochondria | Matrix | 0.0202 | 0.0156 | 0.0146 | 0.0174 | 0.004 | 0.003 | 0.002 | 0.004 | **0.767** | **0.710** | **0.824** | 0.290 | 0.251 | 0.281 | 0.000376 | 0.0001718 | 0.022123 |
| SUCLG1 | 7 | P53597 | Mitochondria | Matrix | 0.0256 | 0.0191 | 0.0173 | 0.0205 | 0.006 | 0.003 | 0.005 | 0.004 | **0.699** | **0.618** | **0.799** | 0.284 | 0.288 | 0.247 | 0.0168594 | 0.0077848 | 0.0311622 |
| SUCLG2 | 11 | A0A024R325 | Mitochondria | Matrix | 0.0109 | 0.0080 | 0.0078 | 0.0085 | 0.002 | 0.002 | 0.002 | 0.002 | **0.732** | **0.706** | **0.798** | 0.234 | 0.238 | 0.281 | 0.0001223 | 0.0004164 | 0.0028578 |
| SOD2 | 9 | P04179 | Mitochondria | Matrix | 0.0477 | 0.0418 | 0.0368 | 0.0431 | 0.013 | 0.009 | 0.011 | 0.008 | **0.897** | **0.804** | **0.873** | 0.601 | 0.606 | 0.354 | 0.8584449 | 0.4409965 | 0.2947923 |
| | | | **Mitochondria** | **Outer-membrane** | 0 | 0 | 0 | 0 | | | | | **0.905** | **0.801** | **0.877** | | | | 0.029472 | 4.774E-06 | 7.52E-05 |
| ACSL1 | 11 | A8K9T3 | Mitochondria | Outer-membrane | 0.0152 | 0.0119 | 0.0099 | 0.0126 | 0.003 | 0.003 | 0.003 | 0.002 | **0.791** | **0.631** | **0.802** | 0.330 | 0.275 | 0.257 | 0.0016463 | 1.324E-06 | 0.0058742 |
| CISD1 | 3 | A0A024QZN7 | Mitochondria | Outer-membrane | 0.0200 | 0.0201 | 0.0180 | 0.0225 | 0.002 | 0.003 | 0.004 | 0.006 | **1.113** | **0.906** | **1.057** | 0.189 | 0.217 | 0.249 | 0.8025932 | 0.709036 | 0.6471334 |
| CPT1B | 12 | A5PLL0 | Mitochondria | Outer-membrane | 0.0126 | 0.0132 | 0.0117 | 0.0136 | 0.003 | 0.003 | 0.003 | 0.002 | **1.002** | **0.960** | **0.978** | 0.705 | 0.653 | 0.563 | 0.9714634 | 0.4071688 | 0.9093935 |
| HK1 | 14 | Q59FD4 | Mitochondria | Outer-membrane | 0.0112 | 0.0113 | 0.0102 | 0.0110 | 0.002 | 0.002 | 0.003 | 0.003 | **0.965** | **0.901** | **0.993** | 0.319 | 0.339 | 0.326 | 0.5170482 | 0.2276241 | 0.9752863 |
| MAOA | 6 | P21397 | Mitochondria | Outer-membrane | 0.0065 | 0.0061 | 0.0047 | 0.0053 | 0.002 | 0.001 | 0.002 | 0.001 | **0.930** | **0.710** | **0.813** | 0.344 | 0.328 | 0.293 | 0.6081042 | 0.0014957 | 0.0313041 |
| MAOB | 8 | P27338 | Mitochondria | Outer-membrane | 0.0097 | 0.0079 | 0.0080 | 0.0080 | 0.002 | 0.002 | 0.002 | 0.002 | **0.828** | **0.741** | **0.856** | 0.360 | 0.318 | 0.278 | 0.2589637 | 0.0939158 | 0.1126635 |
| SAMM50 | 9 | Q9Y512 | Mitochondria | Outer-membrane | 0.0053 | 0.0051 | 0.0046 | 0.0051 | 0.001 | 0.001 | 0.001 | 0.001 | **0.970** | **0.839** | **0.913** | 0.351 | 0.299 | 0.317 | 0.7328837 | 0.1482232 | 0.0234016 |
| VDAC1 | 13 | P21796 | Mitochondria | Outer-membrane | 0.0519 | 0.0475 | 0.0468 | 0.0557 | 0.010 | 0.012 | 0.010 | 0.010 | **0.902** | **0.819** | **0.920** | 0.307 | 0.280 | 0.257 | 0.3348615 | 0.0717145 | 0.0294245 |
| VDAC2 | 14 | A0A024QZN9 | Mitochondria | Outer-membrane | 0.0506 | 0.0401 | 0.0382 | 0.0433 | 0.009 | 0.007 | 0.006 | 0.008 | **0.837** | **0.736** | **0.759** | 0.278 | 0.252 | 0.206 | 0.0636359 | 0.0017026 | 0.0005771 |
| VDAC3 | 10 | Q9Y277 | Mitochondria | Outer-membrane | 0.0265 | 0.0249 | 0.0210 | 0.0270 | 0.011 | 0.008 | 0.009 | 0.010 | **0.832** | **0.770** | **0.780** | 0.381 | 0.358 | 0.403 | 0.0922009 | 0.0090019 | 0.0165998 |
| | | | **Mitochondria** | | 0 | 0 | 0 | 0 | | | | | **0.777** | **0.647** | **0.767** | | | | 0.0459438 | 2.204E-05 | 4.748E-05 |
| BCAT2 | 6 | M0QXF9 | Mitochondria | | 0.0037 | 0.0029 | 0.0025 | 0.0027 | 0.001 | 0.001 | 0.001 | 0.001 | **0.789** | **0.685** | **0.737** | 0.473 | 0.367 | 0.287 | 0.0557696 | 0.0268698 | 0.0139698 |
| C14orf159 | 13 | Q4LE40 | Mitochondria | | 0.0070 | 0.0055 | 0.0041 | 0.0036 | 0.004 | 0.003 | 0.002 | 0.001 | **0.716** | **0.569** | **0.560** | 0.659 | 0.461 | 0.327 | 0.1244339 | 0.0018885 | 0.0014183 |
| C21orf33 | 10 | P30042 | Mitochondria | | 0.0224 | 0.0192 | 0.0152 | 0.0212 | 0.004 | 0.003 | 0.004 | 0.004 | **0.887** | **0.673** | **0.927** | 0.420 | 0.373 | 0.349 | 0.2029467 | 0.0035625 | 0.0887412 |
| DNM1L | 6 | D3DUW5 | Mitochondria | | 0.0034 | 0.0025 | 0.0028 | 0.0045 | 0.004 | 0.004 | 0.004 | 0.004 | **0.875** | **0.923** | **0.909** | 0.570 | 0.589 | 0.620 | 0.5512826 | 0.8665181 | 0.7828565 |
| ECH1 | 12 | Q13011 | Mitochondria | | 0.0437 | 0.0323 | 0.0287 | 0.0301 | 0.006 | 0.009 | 0.009 | 0.010 | **0.705** | **0.588** | **0.644** | 0.640 | 0.469 | 0.316 | 0.0425189 | 0.0006369 | 0.0013182 |
| HSD17B10 | 4 | Q99714 | Mitochondria | | 0.0042 | 0.0036 | 0.0036 | 0.0039 | 0.001 | 0.001 | 0.001 | 0.001 | **0.841** | **0.859** | **0.900** | 0.255 | 0.257 | 0.281 | 0.6616837 | 0.5124673 | 0.9793215 |
| | | | **Nucleus** | | 0 | 0 | 0 | 0 | | | | | **0.948** | **0.901** | **0.917** | | | | 0.5947186 | 0.0849454 | 0.1721071 |
| ADPRHL1 | 11 | Q8NDY3 | Nucleus | Membrane | 0.0089 | 0.0075 | 0.0073 | 0.0078 | 0.002 | 0.002 | 0.002 | 0.002 | **0.899** | **0.804** | **0.835** | 1.416 | 0.867 | 1.358 | 0.3278712 | 0.1046473 | 0.1217115 |
| ANKRD1 | 9 | Q15327 | Nucleus | Transcription factor | 0.0015 | 0.0047 | 0.0038 | 0.0041 | 0.001 | 0.002 | 0.001 | 0.001 | **4.384** | **3.733** | **2.972** | 3.945 | 3.443 | 2.509 | 7.378E-10 | 5.524E-07 | 9.887E-07 |
| ANKRD2 | 7 | Q9GZV1 | Nucleus | Transcription factor | 0.0039 | 0.0022 | 0.0030 | 0.0024 | 0.002 | 0.003 | 0.003 | 0.002 | **0.575** | **0.760** | **0.602** | 0.790 | 0.899 | 0.717 | 0.0086767 | 0.0670415 | 0.036895 |
| ANXA1 | 5 | P04083 | Nucleus | Membrane | 0.0019 | 0.0022 | 0.0018 | 0.0021 | 0.001 | 0.001 | 0.001 | 0.001 | **1.142** | **0.963** | **1.170** | 0.990 | 0.839 | 0.575 | 0.0461857 | 0.1562275 | 0.1412262 |
| ANXA11 | 11 | B7Z6L0 | Nucleus | Membrane | 0.0038 | 0.0045 | 0.0041 | 0.0040 | 0.001 | 0.001 | 0.001 | 0.001 | **1.110** | **1.027** | **1.028** | 0.589 | 0.471 | 0.404 | 0.1047055 | 0.752843 | 0.8485062 |
| ANXA2 | 12 | A0A024R5Z7 | Nucleus | Membrane | 0.0151 | 0.0134 | 0.0127 | 0.0137 | 0.004 | 0.004 | 0.006 | 0.003 | **0.910** | **0.819** | **0.809** | 0.486 | 0.581 | 0.317 | 0.4507393 | 0.2688209 | 0.0182172 |
| ANXA5 | 11 | V9HWE0 | Nucleus | Membrane | 0.0105 | 0.0109 | 0.0104 | 0.0108 | 0.003 | 0.004 | 0.005 | 0.002 | **0.908** | **0.987** | **1.001** | 0.604 | 0.652 | 0.469 | 0.9459159 | 0.8442556 | 0.9106772 |
| ANXA6 | 13 | P08133 | Nucleus | Membrane | 0.0106 | 0.0108 | 0.0105 | 0.0098 | 0.003 | 0.002 | 0.003 | 0.002 | **1.005** | **0.999** | **0.950** | 0.411 | 0.448 | 0.371 | 0.298047 | 0.5628508 | 0.1730077 |
| ANXA7 | 4 | B2R657 | Nucleus | Membrane | 0.0025 | 0.0025 | 0.0026 | 0.0022 | 0.000 | 0.001 | 0.001 | 0.000 | **1.021** | **1.033** | **0.879** | 0.284 | 0.364 | 0.179 | 0.9647768 | 0.4880098 | 0.2205641 |
| APCS | 5 | P02743 | Nucleus | | 0.0120 | 0.0070 | 0.0093 | 0.0102 | 0.010 | 0.009 | 0.014 | 0.006 | **0.606** | **0.770** | **0.684** | 0.807 | 1.393 | 0.697 | 0.0687473 | 0.5106566 | 0.8203048 |
| CLIC4 | 4 | Q9Y696 | Nucleus | | 0.0029 | 0.0032 | 0.0027 | 0.0039 | 0.001 | 0.001 | 0.001 | 0.001 | **1.083** | **0.912** | **1.349** | 0.521 | 0.447 | 0.550 | 0.8128642 | 0.9012859 | 0.0460159 |
| CSRP1 | 5 | P21291 | Nucleus | | 0.0008 | 0.0012 | 0.0009 | 0.0008 | 0.000 | 0.004 | 0.000 | 0.001 | **1.375** | **0.878** | **1.112** | 5.113 | 0.776 | 1.599 | 0.004132 | 0.4556497 | 0.2421189 |
| CRIP2 | 7 | P52943 | Nucleus | Nucleoli | 0.0152 | 0.0169 | 0.0142 | 0.0153 | 0.004 | 0.005 | 0.005 | 0.004 | **1.046** | **0.932** | **0.966** | 0.577 | 0.570 | 0.561 | 0.9168841 | 0.5440778 | 0.7097571 |
| GSTP1 | 5 | V9HWE9 | Nucleus | | 0.0071 | 0.0059 | 0.0056 | 0.0051 | 0.003 | 0.002 | 0.002 | 0.001 | **0.830** | **0.734** | **0.792** | 0.725 | 0.646 | 0.418 | 0.8013969 | 0.5187252 | 0.2995855 |
| HIST1H1C | 7 | P16403; A3R0T7 | Nucleus | Chromosome | 0.0335 | 0.0304 | 0.0311 | 0.0326 | 0.010 | 0.011 | 0.009 | 0.008 | **0.917** | **0.873** | **1.047** | 0.750 | 0.560 | 0.643 | 0.633172 | 0.3945854 | 0.4619799 |
| HIST1H2BJ | 5 | B4DR52; P06899 | Nucleus | Chromosome | 0.0827 | 0.0765 | 0.0711 | 0.0688 | 0.034 | 0.032 | 0.030 | 0.016 | **0.920** | **0.845** | **0.819** | 0.445 | 0.432 | 0.315 | 0.8480147 | 0.4499742 | 0.4147545 |
| HIST1H4A | 5 | P62805 | Nucleus | Chromosome | 0.0785 | 0.0781 | 0.0770 | 0.0623 | 0.028 | 0.034 | 0.031 | 0.018 | **0.958** | **0.990** | **0.688** | 0.689 | 0.503 | 0.321 | 0.7168472 | 0.8424142 | 0.0475567 |
| HNRNPA2B1 | 7 | P22626 | Nucleus | | 0.0050 | 0.0048 | 0.0045 | 0.0040 | 0.001 | 0.002 | 0.001 | 0.001 | **0.949** | **0.885** | **0.831** | 0.422 | 0.375 | 0.244 | 0.5926771 | 0.5649392 | 0.0356399 |
| HNRNPK | 8 | Q5EC54 | Nucleus | | 0.0031 | 0.0029 | 0.0028 | 0.0030 | 0.001 | 0.001 | 0.001 | 0.001 | **0.940** | **0.891** | **0.958** | 0.404 | 0.422 | 0.300 | 0.5267337 | 0.5938302 | 0.6734068 |
| HSPB7 | 7 | Q68DG0 | Nucleus | HSP | 0.0125 | 0.0133 | 0.0131 | 0.0137 | 0.004 | 0.003 | 0.004 | 0.003 | **1.116** | **0.845** | **1.111** | 0.557 | 0.570 | 0.456 | 0.8473451 | 0.5178227 | 0.5016019 |
| LMNA | 13 | W8QEH3; P02545 | Nucleus | Linker | 0.0155 | 0.0156 | 0.0139 | 0.0152 | 0.003 | 0.005 | 0.005 | 0.002 | **1.050** | **0.904** | **0.965** | 0.454 | 0.443 | 0.286 | 0.2564484 | 0.9391629 | 0.477319 |
| MACROD1 | 3 | Q9BQ69 | Nucleus | | 0.0027 | 0.0028 | 0.0027 | 0.0026 | 0.001 | 0.001 | 0.001 | 0.001 | **0.939** | **1.012** | **0.930** | 0.688 | 0.653 | 0.503 | 0.7581412 | 0.9138647 | 0.7142344 |
| NME1-NME2 | 4 | Q32Q12 | Nucleus | Nucleoplasm | 0.0050 | 0.0053 | 0.0046 | 0.0054 | 0.002 | 0.001 | 0.001 | 0.001 | **1.075** | **0.901** | **1.022** | 0.776 | 0.615 | 0.552 | 0.6314125 | 0.2860403 | 0.8284524 |
| PARK7 | 12 | Q99497 | Nucleus | | 0.0235 | 0.0207 | 0.0215 | 0.0244 | 0.006 | 0.005 | 0.004 | 0.002 | **0.918** | **0.906** | **1.095** | 0.858 | 0.740 | 0.885 | 0.5662116 | 0.4664352 | 0.4908212 |
| PGAM2 | 5 | P15259 | Nucleus | Nucleoplasm | 0.0109 | 0.0062 | 0.0060 | 0.0064 | 0.005 | 0.002 | 0.002 | 0.003 | **0.544** | **0.515** | **0.523** | 0.477 | 0.448 | 0.369 | 0.0030792 | 0.0020659 | 0.007218 |
| PKM | 13 | A0A024R5Z9; P146 | Nucleus | | 0.0485 | 0.0380 | 0.0367 | 0.0431 | 0.015 | 0.006 | 0.008 | 0.007 | **0.821** | **0.782** | **0.910** | 0.343 | 0.413 | 0.448 | 0.0760364 | 0.0023323 | 0.2274801 |
| PPP1CB | 4 | V9HW04 | Nucleus | Nucleoplasm | 0.0028 | 0.0033 | 0.0030 | 0.0031 | 0.000 | 0.001 | 0.001 | 0.001 | **1.202** | **1.069** | **1.118** | 0.369 | 0.284 | 0.281 | 0.0048856 | 0.2780906 | 0.1349421 |
| RAN | 6 | B5MDF5 | Nucleus | Transport | 0.0044 | 0.0044 | 0.0044 | 0.0049 | 0.001 | 0.001 | 0.001 | 0.000 | **1.014** | **1.038** | **1.204** | 0.306 | 0.294 | 0.234 | 0.7064991 | 0.5661409 | 0.0741871 |
| SELENBP1 | 8 | Q13228 | Nucleus | Nucleioli | 0.0033 | 0.0017 | 0.0022 | 0.0015 | 0.002 | 0.001 | 0.001 | 0.001 | **0.535** | **0.672** | **0.422** | 0.528 | 0.674 | 0.459 | 3.747E-06 | 0.0007472 | 1.158E-07 |
| SMYD1 | 6 | Q8NB12 | Nucleus | Nucleoplasm | 0.0115 | 0.0102 | 0.0108 | 0.0095 | 0.001 | 0.002 | 0.002 | 0.001 | **0.911** | **0.942** | **0.877** | 0.246 | 0.279 | 0.203 | 0.3665126 | 0.4053293 | 0.1119859 |
| SOD1 | 9 | V9HWC9 | Nucleus | Nucleoplasm | 0.0282 | 0.0238 | 0.0235 | 0.0290 | 0.009 | 0.008 | 0.006 | 0.003 | **0.827** | **0.820** | **0.961** | 0.794 | 0.449 | 0.502 | 0.5273775 | 0.2970236 | 0.8242003 |
| TPI1 | 7 | P60174 | Nucleus | Nucleoplasm | 0.0354 | 0.0306 | 0.0306 | 0.0327 | 0.012 | 0.007 | 0.006 | 0.003 | **0.839** | **0.814** | **0.887** | 0.375 | 0.350 | 0.309 | 0.3089626 | 0.2457828 | 0.5016448 |
| TPI1 shared | 9 | Q2QD09; P60174 | Nucleus | Nucleoplasm | 0.0517 | 0.0423 | 0.0417 | 0.0458 | 0.016 | 0.007 | 0.006 | 0.005 | **0.833** | **0.818** | **0.904** | 0.383 | 0.325 | 0.324 | 0.6746982 | 0.5263415 | 0.9677264 |
| UBA1 | 7 | A0A024R1A3 | Nucleus | | 0.0025 | 0.0024 | 0.0024 | 0.0029 | 0.001 | 0.001 | 0.001 | 0.001 | **0.982** | **0.915** | **1.066** | 0.642 | 0.547 | 0.689 | 0.8292157 | 0.8800524 | 0.8510024 |
| UBE2L3 | 4 | A0A024R1A4 | Nucleus | | 0.0032 | 0.0030 | 0.0030 | 0.0034 | 0.001 | 0.001 | 0.001 | 0.001 | **0.935** | **0.919** | **1.099** | 0.532 | 0.523 | 0.524 | 0.7177303 | 0.4052787 | 0.6398196 |
| UGP2 | 12 | B4DUP2 | Nucleus | | 0.0071 | 0.0077 | 0.0078 | 0.0090 | 0.002 | 0.001 | 0.001 | 0.001 | **1.093** | **1.143** | **1.208** | 0.482 | 0.571 | 0.672 | 0.1435051 | 0.3735266 | 0.1517643 |
| | | | **Peroxisome** | | 0 | 0 | 0 | 0 | | | | | **0.782** | **0.677** | **0.750** | | | | 0.0181067 | 0.0023051 | 0.0023187 |
| ECI2 | 7 | O75521 | Peroxisome | | 0.0043 | 0.0033 | 0.0030 | 0.0030 | 0.001 | 0.001 | 0.001 | 0.001 | **0.767** | **0.708** | **0.684** | 0.365 | 0.358 | 0.297 | 0.0732975 | 0.0072223 | 0.0052019 |
| HSDL2 | 13 | Q6YN16 | Peroxisome | | 0.0322 | 0.0225 | 0.0256 | 0.0213 | 0.018 | 0.014 | 0.015 | 0.007 | **0.733** | **0.636** | **0.650** | 0.537 | 0.484 | 0.431 | 0.021642 | 0.011961 | 0.0051622 |
| GSTK1 | 3 | Q6FII1 | Peroxisome | | 0.0051 | 0.0052 | 0.0047 | 0.0055 | 0.001 | 0.001 | 0.001 | 0.001 | **1.005** | **0.879** | **1.179** | 0.265 | 0.244 | 0.284 | 0.7894769 | 0.6451391 | 0.5020641 |
| | | | **Plasma membrane** | | 0 | 0 | 0 | 0 | | | | | **0.909** | **0.902** | **1.013** | | | | 0.1713431 | 0.0167057 | 0.2173933 |
| ADH1B | 8 | P00325 | Plasma membrane | | 0.0366 | 0.0365 | 0.0392 | 0.0431 | 0.014 | 0.025 | 0.045 | 0.005 | **0.580** | **0.795** | **1.077** | 2.239 | 0.774 | 0.812 | 0.0931892 | 0.3713799 | 0.9224898 |
| ADH5 | 8 | P11766 | Plasma membrane | | 0.0065 | 0.0066 | 0.0065 | 0.0063 | 0.002 | 0.002 | 0.002 | 0.001 | **1.000** | **1.009** | **0.996** | 0.491 | 0.390 | 0.350 | 0.3085685 | 0.6531358 | 0.2240137 |

| | | | | | | | | | | | | | | | | | | | | | |
|---|---|---|---|---|---|---|---|---|---|---|---|---|---|---|---|---|---|---|---|---|---|
| AHNAK | 13 | Q09666 | Plasma membrane | | 0.0146 | 0.0163 | 0.0137 | 0.0154 | 0.004 | 0.005 | 0.005 | 0.003 | 1.146 | 1.029 | 1.067 | 0.499 | 0.511 | 0.380 | 0.0075984 | 0.240913 | 0.4947399 |
| ATP1A1,2,3 shared | 9 | A0A0A0MT26; P05 | Plasma membrane | | 0.0075 | 0.0065 | 0.0055 | 0.0059 | 0.002 | 0.001 | 0.001 | 0.001 | 0.831 | 0.728 | 0.791 | 0.363 | 0.329 | 0.298 | 0.1267228 | 0.0019289 | 0.0300333 |
| ATP1A3 | 10 | A0A0A0MT26 | Plasma membrane | | 0.0037 | 0.0028 | 0.0022 | 0.0026 | 0.001 | 0.001 | 0.001 | 0.001 | 0.727 | 0.566 | 0.686 | 0.333 | 0.267 | 0.249 | 0.0002512 | 5.458E-07 | 0.0009631 |
| ATP1B1 | 5 | A3KLL5 | Plasma membrane | | 0.0057 | 0.0052 | 0.0045 | 0.0046 | 0.001 | 0.001 | 0.001 | 0.001 | 0.910 | 0.833 | 0.792 | 0.300 | 0.275 | 0.223 | 0.4995361 | 0.0957566 | 0.1252874 |
| CPNE3 | 5 | A0A024R994 | Plasma membrane | | 0.0014 | 0.0016 | 0.0015 | 0.0016 | 0.001 | 0.001 | 0.000 | 0.000 | 1.242 | 1.161 | 1.133 | 0.736 | 0.603 | 0.585 | 0.0937912 | 0.2891834 | 0.2217897 |
| DCXR | 7 | Q7Z4W1 | Plasma membrane | | 0.0048 | 0.0039 | 0.0037 | 0.0039 | 0.001 | 0.001 | 0.001 | 0.001 | 0.789 | 0.778 | 0.803 | 0.310 | 0.277 | 0.238 | 0.4371902 | 0.2647506 | 0.2256696 |
| DYSF | 7 | O75923 | Plasma membrane | | 0.0010 | 0.0011 | 0.0011 | 0.0009 | 0.000 | 0.000 | 0.000 | 0.000 | 1.322 | 1.132 | 0.965 | 0.762 | 0.569 | 0.375 | 0.1327308 | 0.3266953 | 0.5892179 |
| EHD4 | 6 | A0A024R9N6 | Plasma membrane | | 0.0018 | 0.0023 | 0.0023 | 0.0023 | 0.000 | 0.000 | 0.001 | 0.000 | 1.227 | 1.236 | 1.271 | 0.361 | 0.475 | 0.287 | 0.0558255 | 0.013312 | 0.5513815 |
| ENO1 | 10 | P06733 | Plasma membrar | Cytosol | 0.0142 | 0.0106 | 0.0134 | 0.0114 | 0.003 | 0.005 | 0.003 | 0.004 | 0.927 | 0.931 | 0.887 | 0.412 | 0.373 | 0.344 | 0.0289542 | 0.3111601 | 0.0111596 |
| NCAM1 | 4 | A0A087WV75 | Plasma membrane | | 0.0020 | 0.0028 | 0.0024 | 0.0029 | 0.001 | 0.001 | 0.001 | 0.001 | 1.642 | 1.339 | 1.523 | 0.610 | 0.510 | 0.551 | 0.0004796 | 0.0117512 | 0.0006292 |
| PGAM1 | 5 | Q53G35 | Plasma membrane | | 0.0068 | 0.0089 | 0.0082 | 0.0099 | 0.002 | 0.002 | 0.002 | 0.002 | 1.243 | 1.206 | 1.600 | 0.822 | 0.708 | 0.872 | 0.0725866 | 0.2975038 | 0.0084662 |
| PLIN4 shared | 12 | Q96Q06; A8E631 | Plasma membrar | Lipid | 0.0190 | 0.0130 | 0.0123 | 0.0134 | 0.005 | 0.004 | 0.003 | 0.004 | 0.623 | 0.595 | 0.647 | 0.256 | 0.251 | 0.235 | 0.0177272 | 0.0106743 | 0.0552937 |
| PLIN4 | 10 | Q96Q06 | Plasma membrar | Lipid | 0.0032 | 0.0022 | 0.0018 | 0.0021 | 0.001 | 0.001 | 0.001 | 0.000 | 0.623 | 0.590 | 0.631 | 0.239 | 0.214 | 0.218 | 1.856E-12 | 1.156E-13 | 1.952E-09 |
| PRKAR1A | 6 | P10644 | Plasma membrar | Kinase | 0.0046 | 0.0045 | 0.0042 | 0.0048 | 0.001 | 0.001 | 0.001 | 0.001 | 0.954 | 0.894 | 1.039 | 0.274 | 0.215 | 0.251 | 0.3018558 | 0.0554978 | 0.5134837 |
| PRKAR2A | 3 | A0A024R2W3 | Plasma membrar | Kinase | 0.0027 | 0.0022 | 0.0022 | 0.0023 | 0.001 | 0.001 | 0.001 | 0.001 | 0.842 | 0.793 | 0.840 | 0.414 | 0.338 | 0.303 | 0.4066673 | 0.3195813 | 0.5188211 |
| PTRF | 6 | Q6NZI2 | Plasma membrar | Calveolae | 0.0197 | 0.0203 | 0.0188 | 0.0173 | 0.003 | 0.005 | 0.003 | 0.002 | 1.065 | 0.950 | 0.879 | 0.372 | 0.280 | 0.213 | 0.6039161 | 0.9262001 | 0.8019357 |
| SLMAP | 9 | Q14BN4 | Plasma membrane | | 0.0057 | 0.0078 | 0.0072 | 0.0075 | 0.001 | 0.002 | 0.002 | 0.001 | 1.324 | 1.258 | 1.309 | 0.503 | 0.491 | 0.432 | 2.114E-05 | 0.0015586 | 0.0001667 |
| | | | **Ribsomal** | | | | | | | | | | 0.972 | 1.031 | 1.068 | | | | 0.4281866 | 0.4523607 | 0.8019646 |
| NACA | 7 | E9PAV3 | Ribsomal | | 0.0033 | 0.0029 | 0.0028 | 0.0030 | 0.001 | 0.001 | 0.001 | 0.000 | 0.892 | 0.849 | 0.863 | 0.316 | 0.310 | 0.252 | 0.2033782 | 0.1430285 | 0.2807582 |
| RPL7A | 4 | P62424 | Ribsomal | | 0.0029 | 0.0023 | 0.0024 | 0.0034 | 0.001 | 0.001 | 0.001 | 0.001 | 0.904 | 0.849 | 1.068 | 0.300 | 0.315 | 0.404 | 0.0442714 | 0.2225934 | 0.2176889 |
| RPS27A | 5 | P62979 | Ribsomal | | 0.0728 | 0.0754 | 0.0748 | 0.0794 | 0.014 | 0.016 | 0.018 | 0.009 | 0.978 | 1.044 | 1.076 | 0.329 | 0.329 | 0.330 | 0.871058 | 0.8492522 | 0.5438982 |
| | | | **Sarcomeric reticulum** | | | | | | | | | | 1.023 | 0.987 | 0.808 | | | | 0.148148 | 0.0487928 | 0.0002953 |
| ATP2A2 | 13 | P16615 | Sarcomeric reticulum | | 0.0336 | 0.0173 | 0.0188 | 0.0167 | 0.007 | 0.006 | 0.005 | 0.005 | 0.515 | 0.548 | 0.496 | 0.226 | 0.192 | 0.175 | 8.061E-24 | 2.884E-18 | 1.433E-22 |
| CASQ2 | 13 | O14958 | Sarcomeric reticulum | | 0.0311 | 0.0405 | 0.0404 | 0.0321 | 0.006 | 0.009 | 0.010 | 0.008 | 1.343 | 1.242 | 0.973 | 0.893 | 0.706 | 0.452 | 0.1239617 | 0.5659388 | 0.3522162 |
| HRC | 12 | B5TMG5 | Sarcomeric reticulum | | 0.0158 | 0.0140 | 0.0145 | 0.0127 | 0.009 | 0.006 | 0.006 | 0.005 | 0.861 | 0.894 | 0.858 | 0.350 | 0.378 | 0.277 | 0.1444466 | 0.2024951 | 0.334294 |
| RYR2 | 13 | Q92736 | Sarcomeric reticulum | | 0.0018 | 0.0020 | 0.0017 | 0.0018 | 0.000 | 0.000 | 0.000 | 0.000 | 1.060 | 0.987 | 0.911 | 0.369 | 0.327 | 0.295 | 0.7227921 | 0.0707704 | 0.0285394 |
| SRL | 12 | Q86TD4 | Sarcomeric reticulum | | 0.0213 | 0.0199 | 0.0204 | 0.0166 | 0.006 | 0.005 | 0.005 | 0.002 | 0.921 | 0.955 | 0.754 | 0.375 | 0.375 | 0.251 | 0.7630001 | 0.6153181 | 0.014313 |
| | | | **Blood** | | | | | | | | | | 2.714 | 2.688 | 4.006 | | | | 3.792E-07 | 6.708E-06 | 2.147E-05 |
| A2M | 5 | P01023 | Blood | Protease Inhibitor | 0.0007 | 0.0011 | 0.0014 | 0.0013 | 0.001 | 0.001 | 0.001 | 0.001 | 1.443 | 1.789 | 1.809 | 1.571 | 1.507 | 1.319 | 0.0560963 | 0.040604 | 0.0465369 |
| ALB | 13 | P02768 | Blood | Plasma | 0.3772 | 0.5693 | 0.7253 | 0.8285 | 0.217 | 0.473 | 0.212 | 0.233 | 1.480 | 1.836 | 2.185 | 1.594 | 1.216 | 1.378 | 6.416E-09 | 6.603E-11 | 3.14E-16 |
| BSG | 4 | B4DNE1 | Blood | | 0.0028 | 0.0024 | 0.0022 | 0.0025 | 0.001 | 0.000 | 0.001 | 0.000 | 0.805 | 0.775 | 0.831 | 0.313 | 0.357 | 0.301 | 0.1044634 | 0.0326026 | 0.2829789 |
| CA1 | 8 | P00915 | Blood | | 0.0010 | 0.0046 | 0.0057 | 0.0062 | 0.001 | 0.004 | 0.006 | 0.005 | 6.782 | 6.794 | 8.670 | 6.868 | 8.052 | 7.530 | 9.725E-15 | 3.89E-14 | 1.617E-13 |
| CA3 | 3 | P07451; A0A024R8 | Blood | | 0.0003 | 0.0026 | 0.0050 | 0.0132 | 0.000 | 0.018 | 0.015 | 0.010 | 7.179 | 15.261 | 40.176 | 54.408 | 46.829 | 36.212 | 9.163E-09 | 8.132E-08 | 1.708E-10 |
| C4A | 14 | P0C0L4 | Blood | | 0.0007 | 0.0009 | 0.0011 | 0.0012 | 0.001 | 0.001 | 0.000 | 0.001 | 1.406 | 1.713 | 1.593 | 2.700 | 2.158 | 3.031 | 0.0150921 | 0.0009151 | 0.0034384 |
| CD36 | 9 | A4D1B1 | Blood | Platlet | 0.0114 | 0.0097 | 0.0086 | 0.0105 | 0.003 | 0.002 | 0.002 | 0.001 | 0.937 | 0.744 | 0.795 | 0.451 | 0.332 | 0.317 | 0.3149417 | 0.0106801 | 0.0746549 |
| CFH | 4 | A0A024R962 | Blood | Secreted | 0.0005 | 0.0009 | 0.0010 | 0.0008 | 0.000 | 0.002 | 0.000 | 0.000 | 1.813 | 2.137 | 1.383 | 3.883 | 2.143 | 2.071 | 0.0428267 | 0.1310041 | 0.565962 |
| CP | 5 | A8K5A4 | Blood | Metal binding | 0.0012 | 0.0011 | 0.0013 | 0.0012 | 0.001 | 0.001 | 0.000 | 0.001 | 0.826 | 0.997 | 0.950 | 1.176 | 0.702 | 0.737 | 0.6757048 | 0.8914496 | 0.7424544 |
| FGA | 10 | P02671 | Blood | | 0.0032 | 0.0038 | 0.0037 | 0.0039 | 0.001 | 0.004 | 0.003 | 0.001 | 1.252 | 1.112 | 1.214 | 1.431 | 0.979 | 0.628 | 0.0114687 | 0.1024472 | 0.8853486 |
| FGB | 10 | V9HVY1 | Blood | | 0.0034 | 0.0037 | 0.0036 | 0.0038 | 0.001 | 0.004 | 0.003 | 0.001 | 1.032 | 1.133 | 1.224 | 1.321 | 0.967 | 0.660 | 0.1392513 | 0.2971635 | 0.1718869 |
| FGG | 10 | P02679 | Blood | | 0.0046 | 0.0053 | 0.0054 | 0.0052 | 0.002 | 0.006 | 0.003 | 0.002 | 1.169 | 1.158 | 1.174 | 1.513 | 0.920 | 0.683 | 0.0018161 | 0.0405382 | 0.1640481 |
| HBB | 7 | Q6V0K9; D9YZU5 | Blood | Red blood cells | 0.0409 | 0.2311 | 0.2132 | 0.2912 | 0.043 | 0.214 | 0.290 | 0.218 | 5.743 | 5.310 | 7.728 | 20.928 | 24.326 | 27.988 | 0.0037602 | 0.0070964 | 0.0479897 |
| HBD | 4 | A0N071 | Blood | Red blood cells | 0.0016 | 0.0094 | 0.0093 | 0.0119 | 0.001 | 0.008 | 0.009 | 0.006 | 6.552 | 6.389 | 7.579 | 8.192 | 8.879 | 7.988 | 3.993E-06 | 4.213E-06 | 1.431E-05 |
| PFKP | 10 | Q01813 | Blood | Platlet | 0.0041 | 0.0053 | 0.0050 | 0.0060 | 0.002 | 0.002 | 0.001 | 0.001 | 1.451 | 1.229 | 1.470 | 0.826 | 0.729 | 0.769 | 0.0086594 | 0.0619819 | 0.009394 |
| PLG | 3 | B2R7F8 | Blood | Extracellular | 0.0009 | 0.0016 | 0.0016 | 0.0016 | 0.001 | 0.001 | 0.001 | 0.000 | 1.748 | 1.683 | 1.692 | 2.265 | 1.609 | 1.283 | 0.0053819 | 0.0144416 | 0.0300304 |
| SERPINB6 | 5 | A0A087X1N8 | Blood | Plasma | 0.0032 | 0.0029 | 0.0029 | 0.0033 | 0.001 | 0.002 | 0.002 | 0.000 | 0.866 | 0.896 | 1.039 | 0.761 | 0.752 | 0.415 | 0.9838722 | 0.7864772 | 0.2615021 |
| SERPINC1 | 5 | A0A024R944 | Blood | Plasma | 0.0013 | 0.0018 | 0.0020 | 0.0023 | 0.001 | 0.002 | 0.001 | 0.001 | 1.332 | 1.376 | 1.693 | 1.847 | 1.263 | 1.044 | 0.0016669 | 0.0004218 | 0.0001531 |
| | | | **Chaperone** | | | | | | | | | | 0.994 | 0.905 | 1.048 | | | | 0.8595961 | 0.5182777 | 0.708957 |
| CCT2 | 4 | P78371 | Chaperone | Actin | 0.0017 | 0.0015 | 0.0014 | 0.0015 | 0.001 | 0.000 | 0.000 | 0.000 | 0.895 | 0.845 | 0.934 | 0.381 | 0.354 | 0.323 | 0.0285748 | 0.008382 | 0.0491339 |
| CRYAB | 11 | P02511 | Chaperone | Actin/Titan/Desmin | 0.2705 | 0.2532 | 0.2283 | 0.2578 | 0.089 | 0.051 | 0.041 | 0.030 | 0.890 | 0.806 | 0.922 | 0.625 | 0.534 | 0.581 | 0.466153 | 0.1688285 | 0.796399 |
| HSP90AA1 | 13 | P07900 | Chaperone | | 0.0101 | 0.0098 | 0.0090 | 0.0099 | 0.003 | 0.003 | 0.002 | 0.001 | 1.002 | 0.962 | 0.977 | 0.565 | 0.489 | 0.453 | 0.686175 | 0.204252 | 0.596722 |
| HSP90AB1 | 13 | P08238 | Chaperone | | 0.0099 | 0.0102 | 0.0095 | 0.0106 | 0.003 | 0.002 | 0.001 | 0.001 | 1.088 | 1.045 | 1.062 | 0.403 | 0.378 | 0.379 | 0.6299435 | 0.8381006 | 0.7599169 |
| HSP90AA shared | 13 | P07900; P08238 | Chaperone | | 0.0191 | 0.0180 | 0.0169 | 0.0184 | 0.005 | 0.003 | 0.003 | 0.002 | 0.956 | 0.948 | 0.949 | 0.502 | 0.452 | 0.444 | 0.64755 | 0.9626696 | 0.8458278 |
| HSP90B1 | 11 | P14625 | Chaperone | | 0.0041 | 0.0042 | 0.0042 | 0.0052 | 0.001 | 0.001 | 0.001 | 0.001 | 1.194 | 1.069 | 1.159 | 0.640 | 0.683 | 0.545 | 0.2349048 | 0.2123443 | 0.1815986 |
| HSPA1A | 13 | P08107 | Chaperone | | 0.0225 | 0.0197 | 0.0193 | 0.0218 | 0.004 | 0.003 | 0.003 | 0.003 | 0.897 | 0.871 | 0.938 | 0.295 | 0.258 | 0.240 | 0.2214734 | 0.0184549 | 0.4825885 |
| HSPA8 | 13 | V9HW22 | Chaperone | | 0.0255 | 0.0265 | 0.0264 | 0.0271 | 0.006 | 0.004 | 0.004 | 0.003 | 0.859 | 0.883 | 0.762 | 0.522 | 0.434 | 0.268 | 0.7217326 | 0.2496313 | 0.0995938 |
| HSPB1 | 13 | V9HW43 | Chaperone | | 0.0701 | 0.0873 | 0.0785 | 0.0912 | 0.029 | 0.017 | 0.011 | 0.009 | 1.198 | 1.067 | 1.091 | 0.988 | 0.779 | 0.678 | 0.3555824 | 0.9689828 | 0.6903475 |
| HSPB6 | 7 | O14558 | Chaperone | | 0.0391 | 0.0522 | 0.0437 | 0.0656 | 0.020 | 0.016 | 0.015 | 0.017 | 1.246 | 1.084 | 1.670 | 0.780 | 0.715 | 0.950 | 0.0192347 | 0.0635883 | 0.0001968 |
| UNC45B | 7 | Q8IWX7 | Chaperone | Alpha-Beta myosin | 0.0024 | 0.0021 | 0.0021 | 0.0024 | 0.001 | 0.000 | 0.000 | 0.000 | 0.959 | 0.981 | 1.070 | 0.438 | 0.467 | 0.459 | 0.1361765 | 0.3530667 | 0.4658289 |
| | | | **Cytoplasmic** | | | | | | | | | | 1.473 | 1.235 | 1.536 | | | | 0.1813334 | 0.0047527 | 0.0132482 |
| ACO1 | 8 | P21399 | Cytoplasmic | | 0.0026 | 0.0027 | 0.0025 | 0.0032 | 0.001 | 0.001 | 0.001 | 0.001 | 1.002 | 1.000 | 1.006 | 0.792 | 0.548 | 0.517 | 0.7005795 | 0.4016901 | 0.7262996 |
| AK1 | 10 | Q5T9B7 | Cytoplasmic | Membrane | 0.0259 | 0.0225 | 0.0244 | 0.0254 | 0.008 | 0.004 | 0.005 | 0.002 | 0.884 | 0.928 | 0.981 | 0.301 | 0.341 | 0.328 | 0.0795554 | 0.1659849 | 0.8024418 |
| AKR1B1 | 8 | P15121 | Cytoplasmic | | 0.0096 | 0.0103 | 0.0093 | 0.0121 | 0.004 | 0.007 | 0.003 | 0.003 | 1.103 | 0.956 | 1.192 | 0.987 | 0.569 | 0.643 | 0.248053 | 0.6900308 | 0.0121768 |
| ALDH9A1 | 13 | P49189 | Cytoplasmic | | 0.0049 | 0.0051 | 0.0046 | 0.0065 | 0.001 | 0.001 | 0.001 | 0.001 | 1.116 | 1.012 | 1.200 | 0.582 | 0.530 | 0.498 | 0.4249474 | 0.9943817 | 0.7354124 |
| ALDOA | 12 | P04075 | Cytoplasmic | M-line | 0.0985 | 0.0823 | 0.0760 | 0.0710 | 0.036 | 0.014 | 0.016 | 0.019 | 0.831 | 0.782 | 0.720 | 0.399 | 0.368 | 0.332 | 0.0493665 | 0.018211 | 0.0017892 |
| CAMK2D | 5 | E9PF82 | Cytoplasmic | Membrane | 0.0031 | 0.0035 | 0.0033 | 0.0031 | 0.001 | 0.001 | 0.001 | 0.000 | 1.102 | 1.044 | 1.065 | 0.347 | 0.278 | 0.290 | 0.1885421 | 0.7926829 | 0.5333557 |
| CAST | 6 | E7EQ12 | Cytoplasmic | Membrane | 0.0023 | 0.0016 | 0.0014 | 0.0015 | 0.001 | 0.000 | 0.001 | 0.001 | 0.692 | 0.640 | 0.622 | 0.452 | 0.427 | 0.428 | 0.0092504 | 0.0209364 | 0.0154032 |
| CKB | 8 | V9HW77 | Cytoplasmic | Energy | 0.0254 | 0.0157 | 0.0165 | 0.0191 | 0.011 | 0.005 | 0.005 | 0.005 | 0.622 | 0.632 | 0.747 | 0.337 | 0.321 | 0.359 | 1.16E-06 | 2.479E-07 | 0.0001128 |
| CKM | 12 | P06732 | Cytoplasmic | Energy | 0.2731 | 0.1470 | 0.1585 | 0.1477 | 0.074 | 0.029 | 0.039 | 0.059 | 0.557 | 0.544 | 0.513 | 0.201 | 0.225 | 0.239 | 3.172E-09 | 8.802E-08 | 6.085E-08 |
| CMBL | 8 | B2RD36 | Cytoplasmic | | 0.0034 | 0.0054 | 0.0045 | 0.0047 | 0.001 | 0.001 | 0.001 | 0.001 | 1.714 | 1.337 | 1.448 | 0.533 | 0.575 | 0.402 | 5.29E-07 | 0.0012738 | 2.89E-05 |
| CRYM | 8 | Q14894 | Cytoplasmic | | 0.0118 | 0.0159 | 0.0137 | 0.0171 | 0.003 | 0.005 | 0.003 | 0.002 | 1.339 | 1.162 | 1.407 | 0.541 | 0.425 | 0.437 | 0.0025898 | 0.1746163 | 9.565E-05 |
| CRYZ | 6 | Q08257 | Cytoplasmic | | 0.0038 | 0.0034 | 0.0041 | 0.0045 | 0.001 | 0.001 | 0.001 | 0.001 | 0.840 | 1.064 | 1.172 | 0.360 | 0.390 | 0.359 | 0.0628866 | 0.5678779 | 0.014186 |
| CSRP3 | 10 | A2TDB8 | Cytoplasmic | | 0.0685 | 0.0997 | 0.0924 | 0.0992 | 0.016 | 0.020 | 0.019 | 0.016 | 1.435 | 1.309 | 1.456 | 0.727 | 0.613 | 0.572 | 0.0289696 | 0.2284407 | 0.1578432 |
| DLD | 11 | A0A024R713 | Cytoplasmic | | 0.0482 | 0.0513 | 0.0474 | 0.0525 | 0.008 | 0.006 | 0.007 | 0.006 | 1.050 | 0.940 | 1.088 | 0.293 | 0.294 | 0.268 | 0.4689546 | 0.6230141 | 0.4789358 |
| EEF1A1 | 8 | Q05639; P68104 | Cytoplasmic | Signal | 0.0394 | 0.0376 | 0.0359 | 0.0372 | 0.012 | 0.009 | 0.009 | 0.004 | 1.082 | 0.921 | 0.938 | 1.391 | 0.968 | 1.070 | 0.8669132 | 0.642007 | 0.6024481 |
| EEF1A2 | 7 | Q05639 | Cytoplasmic | Signal | 0.0110 | 0.0111 | 0.0097 | 0.0086 | 0.007 | 0.005 | 0.004 | 0.001 | 0.979 | 0.861 | 1.006 | 0.875 | 0.752 | 0.633 | 0.7291584 | 0.0505485 | 0.004193 |
| EEF2 | 12 | P13639 | Cytoplasmic | Signal | 0.0073 | 0.0060 | 0.0057 | 0.0062 | 0.002 | 0.001 | 0.001 | 0.001 | 0.854 | 0.804 | 0.854 | 0.277 | 0.258 | 0.261 | 0.0217014 | 0.0001421 | 0.0438186 |
| EIF4A2 | 4 | Q14240 | Cytoplasmic | Ribosome | 0.0031 | 0.0027 | 0.0027 | 0.0028 | 0.001 | 0.001 | 0.001 | 0.000 | 0.917 | 0.938 | 0.905 | 0.372 | 0.386 | 0.312 | 0.3765093 | 0.2755259 | 0.3330724 |
| ENO3 | 5 | P13929; B4DUJ6 | Cytoplasmic | | 0.0222 | 0.0161 | 0.0223 | 0.0193 | 0.007 | 0.006 | 0.008 | 0.004 | 0.800 | 0.996 | 0.919 | 0.354 | 0.474 | 0.312 | 0.038536 | 0.854834 | 0.6506528 |
| ESD | 7 | P10768 | Cytoplasmic | Vesicle | 0.0045 | 0.0038 | 0.0036 | 0.0040 | 0.001 | 0.001 | 0.001 | 0.000 | 0.831 | 0.816 | 0.848 | 0.489 | 0.472 | 0.254 | 0.5730625 | 0.4434015 | 0.2115178 |
| FABP3 | 9 | P05413 | Cytoplasmic | | 0.3330 | 0.3366 | 0.3116 | 0.3558 | 0.081 | 0.067 | 0.063 | 0.036 | 1.001 | 0.929 | 1.114 | 0.397 | 0.330 | 0.320 | 0.7839952 | 0.078165 | 0.3776909 |
| FABP4 | 3 | E7DVW4 | Cytoplasmic | | 0.0081 | 0.0072 | 0.0058 | 0.0076 | 0.004 | 0.003 | 0.002 | 0.003 | 0.871 | 0.781 | 0.477 | 0.743 | 0.749 | 0.504 | 0.3359143 | 0.102704 | 0.2359192 |
| FABP5 | 5 | Q01469 | Cytoplasmic | | 0.0099 | 0.0072 | 0.0060 | 0.0079 | 0.002 | 0.002 | 0.003 | 0.003 | 0.709 | 0.639 | 0.702 | 0.258 | 0.293 | 0.305 | 0.0031317 | 0.000542 | 0.0620376 |
| FAM129A | 7 | A0A024R978 | Cytoplasmic | | 0.0024 | 0.0032 | 0.0028 | 0.0030 | 0.001 | 0.001 | 0.001 | 0.001 | 1.373 | 1.109 | 1.182 | 0.612 | 0.495 | 0.386 | 0.0045393 | 0.2326195 | 0.6734131 |
| FAM162A | 5 | Q96A26 | Cytoplasmic | | 0.0065 | 0.0053 | 0.0047 | 0.0045 | 0.002 | 0.002 | 0.002 | 0.002 | 0.979 | 0.649 | 0.480 | 0.593 | 0.547 | 0.234 | 0.6727511 | 0.2200367 | 0.22512 |
| FHL2 | 13 | A0A0A0MSG2 | Cytoplasmic | Sarcomeric | 0.0792 | 0.0609 | 0.0673 | 0.0235 | 0.023 | 0.023 | 0.017 | 0.019 | 0.756 | 0.850 | 0.311 | 0.413 | 0.366 | 0.287 | 0.0013205 | 0.0111586 | 7.325E-15 |
| FTH1 | 7 | P02794 | Cytoplasmic | | 0.0068 | 0.0086 | 0.0065 | 0.0075 | 0.004 | 0.003 | 0.003 | 0.004 | 1.104 | 0.933 | 1.119 | 0.854 | 0.809 | 0.832 | 0.9556028 | 0.2157733 | 0.2780656 |
| GAPDH | 10 | P04406 | Cytoplasmic | | 0.0930 | 0.0923 | 0.0814 | 0.1048 | 0.025 | 0.019 | 0.023 | 0.018 | 1.024 | 0.864 | 1.084 | 0.808 | 0.599 | 0.520 | 0.9796776 | 0.1871667 | 0.7108868 |
| GBAS | 7 | O75323 | Cytoplasmic | | 0.0234 | 0.0211 | 0.0208 | 0.0228 | 0.008 | 0.005 | 0.005 | 0.006 | 0.989 | 0.868 | 0.982 | 1.404 | 0.840 | 0.366 | 0.4075056 | 0.7262454 | 0.6702991 |
| GBE1 | 5 | Q59ET0 | Cytoplasmic | | 0.0021 | 0.0028 | 0.0023 | 0.0030 | 0.001 | 0.001 | 0.000 | 0.001 | 1.384 | 1.113 | 1.419 | 0.546 | 0.430 | 0.567 | 0.0024874 | 0.1766607 | 0.0006659 |
| GDI1 | 5 | P31150; B4DLV7 | Cytoplasmic | | 0.0026 | 0.0026 | 0.0026 | 0.0026 | 0.001 | 0.001 | 0.001 | 0.001 | 1.020 | 1.026 | 1.038 | 1.743 | 0.951 | 0.276 | 0.3160686 | 0.3588079 | 0.0726577 |
| GDI2 | 4 | B4DLV7 | Cytoplasmic | | 0.0025 | 0.0027 | 0.0029 | 0.0025 | 0.001 | 0.001 | 0.001 | 0.000 | 1.069 | 1.187 | 0.970 | 0.321 | 0.353 | 0.252 | 0.352678 | 0.3004148 | 0.831542 |
| GLRX | 4 | A0A024RAM2 | Cytoplasmic | | 0.0065 | 0.0076 | 0.0071 | 0.0087 | 0.001 | 0.002 | 0.001 | 0.002 | 1.217 | 1.085 | 1.328 | 0.450 | 0.326 | 0.424 | 0.0091452 | 0.0494064 | 3.026E-05 |
| GPD1L | 8 | Q8N335 | Cytoplasmic | | 0.0090 | 0.0056 | 0.0064 | 0.0072 | 0.003 | 0.002 | 0.002 | 0.002 | 0.616 | 0.689 | 0.706 | 0.276 | 0.319 | 0.359 | 0.000359 | 0.0078843 | 0.073432 |
| GRHPR | 7 | Q9UBQ7 | Cytoplasmic | | 0.0051 | 0.0045 | 0.0039 | 0.0043 | 0.001 | 0.001 | 0.001 | 0.001 | 0.896 | 0.757 | 0.919 | 0.406 | 0.334 | 0.425 | 0.5531712 | 0.0240611 | 0.9016251 |
| GSTM3 | 5 | Q59EJ5 | Cytoplasmic | | 0.0026 | 0.0017 | 0.0018 | 0.0020 | 0.001 | 0.001 | 0.001 | 0.001 | 0.649 | 0.669 | 0.790 | 0.636 | 0.692 | 0.434 | 0.2967786 | 0.3002434 | 0.0620318 |
| GSTO1 | 6 | B2R983 | Cytoplasmic | | 0.0067 | 0.0058 | 0.0059 | 0.0059 | 0.002 | 0.002 | 0.001 | 0.002 | 0.844 | 0.902 | 0.905 | 0.448 | 0.456 | 0.446 | 0.0291384 | 0.163017 | 0.1422069 |
| GYG1 | 4 | B2R5R5 | Cytoplasmic | | 0.0062 | 0.0071 | 0.0066 | 0.0075 | 0.002 | 0.002 | 0.002 | 0.002 | 1.178 | 1.091 | 1.306 | 0.514 | 0.496 | 0.489 | 0.4402965 | 0.7177265 | 0.3406643 |
| GYS1 | 11 | P13807 | Cytoplasmic | | 0.0048 | 0.0064 | 0.0060 | 0.0070 | 0.001 | 0.001 | 0.001 | 0.002 | 1.318 | 1.191 | 1.363 | 0.701 | 0.560 | 0.559 | 0.1636531 | 0.1910915 | 0.2879013 |
| HBA2 | 10 | D1MGQ2 | Cytoplasmic | | 0.0463 | 0.2935 | 0.2595 | 0.3079 | 0.041 | 0.224 | 0.291 | 0.187 | 6.289 | 4.912 | 6.661 | 10.366 | 11.970 | 7.110 | 5.153E-05 | 0.0008404 | 0.1242585 |
| IARS2 | 7 | Q9NSE4 | Cytoplasmic | | 0.0028 | 0.0028 | 0.0026 | 0.0028 | 0.001 | 0.001 | 0.001 | 0.001 | 1.027 | 0.929 | 0.998 | 0.419 | 0.371 | 0.320 | 0.3696164 | 0.7861828 | 0.8994855 |
| LDHA | 11 | P00338 | Cytoplasmic | | 0.0106 | 0.0080 | 0.0078 | 0.0102 | 0.005 | 0.003 | 0.003 | 0.004 | 0.793 | 0.767 | 0.879 | 0.958 | 0.967 | 0.888 | 0.2435939 | 0.5223632 | 0.4698767 |
| LDHB | 10 | Q5U077 | Cytoplasmic | | 0.1121 | 0.0849 | 0.0832 | 0.1004 | 0.042 | 0.015 | 0.018 | 0.024 | 0.788 | 0.693 | 0.887 | 0.632 | 0.689 | 0.581 | 0.100073 | 0.0374894 | 0.2429559 |
| MB | 12 | P02144 | Cytoplasmic | Oxygen reserve | 1.0792 | 0.7715 | 0.8368 | 0.8017 | 0.315 | 0.189 | 0.235 | 0.219 | 0.781 | 0.784 | 0.748 | 0.674 | 0.669 | 0.425 | 0.1816675 | 0.2640976 | 0.0045685 |
| MDH1 | 8 | B9A041; P40925 | Cytoplasmic | | 0.1092 | 0.0918 | 0.0827 | 0.0974 | 0.028 | 0.015 | 0.013 | 0.016 | 0.825 | 0.756 | 0.845 | 0.327 | 0.317 | 0.303 | 0.1586659 | 0.0811047 | 0.4483361 |
| NPEPPS | 7 | P55786 | Cytoplasmic | | 0.0021 | 0.0017 | 0.0017 | 0.0019 | 0.001 | 0.000 | 0.000 | 0.000 | 0.896 | 0.822 | 0.867 | 0.444 | 0.409 | 0.340 | 0.100668 | 0.0713089 | 0.2105336 |
| PACSIN3 | 13 | P40926 | Cytoplasmic | | 0.1273 | 0.0993 | 0.0843 | 0.0957 | 0.034 | 0.014 | 0.024 | 0.014 | 0.768 | 0.662 | 0.759 | 0.216 | 0.224 | 0.214 | 0.0009475 | 3.373E-06 | 0.000453 |
| PCMT1 | 5 | H7BY58 | Cytoplasmic | | 0.0058 | 0.0052 | 0.0056 | 0.0053 | 0.001 | 0.001 | 0.001 | 0.001 | 0.945 | 0.954 | 0.873 | 0.269 | 0.282 | 0.246 | 0.1305846 | 0.2785355 | 0.2847928 |
| PEBP1 | 7 | D9IAI1 | Cytoplasmic | | 0.0347 | 0.0306 | 0.0310 | 0.0315 | 0.007 | 0.007 | 0.006 | 0.004 | 0.843 | 0.923 | 0.865 | 0.270 | 0.245 | 0.204 | 0.2798498 | 0.2790935 | 0.2766184 |
| PFKM | 13 | P08237 | Cytoplasmic | | 0.0227 | 0.0173 | 0.0168 | 0.0181 | 0.006 | 0.003 | 0.004 | 0.003 | 0.762 | 0.696 | 0.762 | 0.275 | 0.295 | 0.248 | 0.000103 | 0.0002075 | 0.0005159 |
| PFN1 | 6 | P07737 | Cytoplasmic | | 0.0078 | 0.0090 | 0.0089 | 0.0096 | 0.002 | 0.003 | 0.002 | 0.002 | 1.143 | 1.093 | 1.066 | 0.760 | 0.686 | 0.319 | 0.586528 | 0.7457952 | 0.6754383 |
| PGK1 | 12 | P00558 | Cytoplasmic | Kinase | 0.0445 | 0.0404 | 0.0421 | 0.0488 | 0.013 | 0.006 | 0.007 | 0.004 | 0.945 | 0.933 | 1.061 | 0.289 | 0.318 | 0.310 | 0.6576656 | 0.774521 | 0.3021952 |
| PRDX1 | 8 | Q06830 | Cytoplasmic | | 0.0144 | 0.0127 | 0.0123 | 0.0134 | 0.004 | 0.003 | 0.003 | 0.002 | 0.926 | 0.846 | 0.884 | 0.381 | 0.351 | 0.301 | 0.3776729 | 0.2349275 | 0.3842244 |
| PRDX2 | 9 | V9HW12 | Cytoplasmic | | 0.0374 | 0.0343 | 0.0327 | 0.0349 | 0.013 | 0.007 | 0.007 | 0.005 | 0.894 | 0.899 | 0.925 | 0.446 | 0.402 | 0.416 | 0.5218556 | 0.5352131 | 0.8906532 |
| PRKACA | 6 | P17612 | Cytoplasmic | Kinase | 0.0040 | 0.0033 | 0.0035 | 0.0036 | 0.001 | 0.001 | 0.001 | 0.001 | 0.812 | 0.885 | 0.907 | 0.457 | 0.451 | 0.360 | 0.1123397 | 0.1110898 | 0.3142553 |
| PYGB | 13 | P11216; Q59GM9 | Cytoplasmic | | 0.0109 | 0.0160 | 0.0151 | 0.0162 | 0.004 | 0.004 | 0.003 | 0.002 | 1.418 | 1.362 | 1.489 | 0.748 | 0.650 | 0.673 | 1.846E-06 | 0.0006103 | 0.0023727 |
| PYGM | 12 | B2RB32 | Cytoplasmic | | 0.0038 | 0.0031 | 0.0035 | 0.0034 | 0.001 | 0.001 | 0.001 | 0.001 | 0.780 | 0.921 | 0.937 | 0.614 | 0.586 | 0.518 | 0.0147096 | 0.2583489 | 0.3040293 |
| RNH1 | 12 | P13489 | Cytoplasmic | | 0.0050 | 0.0044 | 0.0043 | 0.0051 | 0.003 | 0.001 | 0.001 | 0.001 | 0.864 | 0.865 | 0.963 | 0.507 | 0.474 | 0.464 | 0.0662486 | 0.1459937 | 0.5840191 |
| RSU1 | 5 | Q32Q10 | Cytoplasmic | | 0.0021 | 0.0022 | 0.0019 | 0.0019 | 0.000 | 0.001 | 0.000 | 0.000 | 1.086 | 0.983 | 1.079 | 0.502 | 0.388 | 0.231 | 0.2039961 | 0.8808879 | 0.5031315 |
| SDPR | 8 | A8K2W3 | Cytoplasmic | Caveolae | 0.0048 | 0.0035 | 0.0032 | 0.0029 | 0.001 | 0.001 | 0.001 | 0.000 | 0.769 | 0.698 | 0.620 | 0.339 | 0.302 | 0.218 | 0.0002891 | 0.0001138 | 3.141E-05 |
| SPR | 5 | P35270 | Cytoplasmic | | 0.0018 | 0.0019 | 0.0019 | 0.0022 | 0.000 | 0.001 | 0.002 | 0.002 | 1.064 | 0.987 | 1.238 | 0.544 | 0.630 | 0.658 | 0.1127544 | 0.207708 | 0.0101491 |
| ST13 | 5 | P50502 | Cytoplasmic | | 0.0035 | 0.0039 | 0.0038 | 0.0041 | 0.001 | 0.001 | 0.001 | 0.001 | 1.082 | 1.105 | 1.142 | 0.479 | 0.421 | 0.401 | 0.1435739 | 0.2720638 | 0.115866 |
| TMOD1 | 14 | P28289 | Cytoplasmic | Sarcomeric | 0.0078 | 0.0067 | 0.0070 | 0.0064 | 0.001 | 0.001 | 0.001 | 0.001 | 0.901 | 0.901 | 0.800 | 0.228 | 0.237 | 0.183 | 0.1532216 | 0.2616689 | 0.0019001 |
| | | | **Miscellaneous** | **Unknown** | | | | | | | | | 1.479 | 1.652 | 1.774 | | | | 0.9465916 | 0.0351965 | 0.1872465 |
| YWHAE | 6 | G9K388 | Miscellaneous | Unknown | 0.0156 | 0.0159 | 0.0152 | 0.0151 | 0.003 | 0.003 | 0.003 | 0.002 | 1.020 | 0.982 | 0.952 | 0.326 | 0.323 | 0.249 | 0.8661065 | 0.2175384 | 0.2701099 |
| YWHAG | 8 | B3KNB4 | Miscellaneous | Unknown | 0.0100 | 0.0103 | 0.0098 | 0.0099 | 0.003 | 0.003 | 0.002 | 0.002 | 1.052 | 0.927 | 1.058 | 0.370 | 0.351 | 0.333 | 0.3797277 | 0.7184273 | 0.8618205 |
| YWHAZ | 6 | P63104 | Miscellaneous | Unknown | 0.0080 | 0.0087 | 0.0076 | 0.0082 | 0.001 | 0.002 | 0.002 | 0.002 | 1.082 | 0.959 | 1.081 | 0.402 | 0.368 | 0.284 | 0.0726131 | 0.7812848 | 0.7760774 |
| blank | 8 | B4E1Z4 | Miscellaneous | Unknown | 0.0011 | 0.0013 | 0.0014 | 0.0014 | 0.001 | 0.002 | 0.001 | 0.001 | 1.174 | 1.421 | 1.394 | 2.165 | 1.316 | 1.327 | 0.0490759 | 0.0569329 | 0.1202222 |
| blank | 5 | Q8TCD0 | Miscellaneous | Unknown | 0.0221 | 0.0388 | 0.0452 | 0.0472 | 0.026 | 0.032 | 0.018 | 0.031 | 1.881 | 2.159 | 2.320 | 2.413 | 2.348 | 2.675 | 0.0067174 | 0.00068 | 0.0007167 |
| | | | **Proeasome** | | | | | | | | | | 0.869 | 0.893 | 0.867 | | | | 0.304253 | 0.2747861 | 0.2062896 |
| PSMA2 | 2 | P25787 | Proteasome | | 0.0003 | 0.0005 | 0.0004 | 0.0003 | 0.000 | 0.000 | 0.000 | 0.000 | 1.335 | 1.040 | 0.835 | 1.530 | 1.140 | 0.864 | 0.7935151 | 0.8689651 | 0.2480135 |
| PSMA3 | 2 | P25788 | Proteasome | | 0.0023 | 0.0016 | 0.0024 | 0.0020 | 0.002 | 0.004 | 0.002 | 0.001 | 0.716 | 1.041 | 0.857 | 1.905 | 1.276 | 0.730 | 0.1415905 | 0.3906674 | 0.1628383 |
| PSMA5 | 1 | P28066 | Proteasome | | 0.0026 | 0.0021 | 0.0020 | 0.0021 | 0.001 | 0.000 | 0.000 | 0.000 | 0.793 | 0.781 | 0.789 | 0.225 | 0.235 | 0.219 | 0.2543669 | 0.1241869 | 0.1813337 |
| PSMA7 | 2 | O14818 | Proteasome | | 0.0015 | 0.0013 | 0.0012 | 0.0014 | 0.001 | 0.000 | 0.000 | 0.000 | 0.897 | 0.842 | 0.931 | 0.569 | 0.537 | 0.464 | 0.1694569 | 0.1667385 | 0.384064 |
| PSMB3 | 2 | P49720 | Proteasome | | 0.0008 | 0.0006 | 0.0007 | 0.0009 | 0.000 | 0.000 | 0.000 | 0.000 | 0.911 | 0.828 | 1.016 | 0.982 | 0.778 | 0.780 | 0.8384268 | 0.6545277 | 0.8090663 |
| PSMB6 | 2 | P28072 | Proteasome | | 0.0022 | 0.0021 | 0.0020 | 0.0019 | 0.001 | 0.000 | 0.000 | 0.000 | 0.928 | 0.857 | 0.854 | 0.305 | 0.315 | 0.324 | 0.5162544 | 0.4015117 | 0.4644119 |
