## Supplemental methods and figures for "Metabolic and Proteomic Defects in Human Hypertrophic Cardiomyopathy"

### Supplemental Material

### Expanded Materials and Methods

**Patient sample acquisition.** HCM was diagnosed on the basis of standard criteria and in the absence of any cause for secondary hypertrophy. Genetic testing was performed on a clinical basis by Clinical Laboratory Improvement Amendments–certified laboratories using standard HCM gene panels (including at least all standard sarcomere genes and infiltrative genetic causes). Patient demographic and clinical data were recorded at the time of tissue collection, including genotype status. All variants in *MYBPC3* were truncating variants, and all variants in *MYH7* were missense variants. HCM hearts were perfused with warm blood cardioplegia before tissue resection. Myectomy tissue was snap-frozen in liquid $N_2$ immediately in the OR upon excision. Control donor hearts were identified through a research protocol with Gift of Life and all had normal systolic function and minimal to no hypertrophy. Donor hearts were perfused with HTK-enriched (histidine, tryptophan, and ketogluterate) cold cardioplegia solution before removal, and tissue was snap-frozen in liquid $N_2$ in the operating room immediately upon explantation. Due to unavoidable differences in the cardioplegia solution between control and HCM hearts, we discounted blood cell-derived proteins in the proteomic analysis, as well as histidine, tryptophan and alpha-ketogluterate values in the metabolomics analysis.

**Sample preparation for proteomic analyses.** Each piece of muscle was solubilized in 150 µl 0.1% RapiGest SF Surfactant (Waters Corporation) in a dissection chamber by mechanical trituration with forceps. The sample was transferred to a 1.5 ml microcentrifuge tube using a pipette and heated at 50°C for 1 hour. The samples were reduced by adding 0.75 µl of 1M dithiothreitol and heating (100°C, 10 min), then alkylated with 22.5 µl of 100 mM iodoacetamide in 50 mM ammonium bicarbonate with incubation in the dark (22°C, 30 min). The proteins were digested to peptides with the addition of 25 µl of 0.2 µg/µl trypsin (Promega) and incubated (37°C, 18 hours). The samples were dried down in a speed vacuum device. A 100-µl aliquot of 7% formic acid in 50 mM ammonium bicarbonate was added and incubated (37°C, 1 hour), to deactivate the trypsin and cleave the RapiGest. The sample was dried down again. A 100-µl aliquot of 0.1% trifluoroacetic acid was added and incubated (37°C, 1 hour) to cleave the RapiGest again. The resultant peptides were dried down and reconstituted in 150 µl of 0.1% trifluoroacetic acid. The tubes were centrifuged at 14,000 rpms for 5 minutes to pellet the surfactant. The top 125 µl of solution was transferred into a mass spectrometry analysis vial.

**Liquid chromatography mass spectrometry (LCMS) for proteomic analyses.** Liquid chromatography was performed on an Acquity UPLC HSS T3 column (100 Å, 1.8 µm, 1 × 150 mm) (Waters Corporation) attached to an UltiMate 3000 ultra-high pressure liquid chromatography (UHPLC) system (Dionex). A 20 µL aliquot of each sample was injected into 0.1% formic acid in 2% acetonitrile with a flow rate of 100 µL/min. Starting at 2 min, the gradient was ramped linearly to 35% acetonitrile over 90 min, followed by another linear increase to 50% acetonitrile over 10 min. Finally, the gradient was ramped linearly to 90% acetonitrile over 6 sec, and held isocratic for 6 min. The column was then re-equilibrated over 27 min, prior to the next injection. The total run time was 135 min per injection. Formic acid was kept at 0.1% concentration in the mobile phase during all acetonitrile ramping steps. The UHPLC effluent was directly infused into a Q Exactive Hybrid Quadrupole-Orbitrap mass spectrometer through an electrospray ionization source (Thermo Fisher Scientific). Peptides were identified from the MS/MS spectra using SEQUEST run through the Proteome Discoverer 2.2 (PD 2.2) software package (Thermo Fisher Scientific) to search against the human proteome database (downloaded from UniProt 2/15/2018). Variable mass changes were accounting for the potential loss of methionine from the N-terminus of each protein (-131.20 Da), the loss of methionine with addition of acetylation (-89.16 Da), addition of carbamidomethyl (C; 57.02 Da), oxidation (M, P; 15.99 Da: M; 32.00 Da) and phosphorylation (S, T, Y; 79.98 Da) were added to the searches. Raw peptide abundances were determined from the area under each LCMS peak using PD 2.2 with the Minora Feature Detector enabled. The Minora Feature Detector provides generates peak areas for all LCMS peaks matching the exact mass, charge states, elution time, and isotope pattern of the SEQUEST derived peptide spectral matches (PSMs).

**Quantification of protein abundances and classification of subcellular compatmentalization.** The raw peptide abundances were exported to Excel (Microsoft) and then normalized by dividing their abundance by the mean abundance of the top 15 peptides shared between α- and β-myosin heavy chain isoforms (MYH6 and MYH7). This normalization scheme corrects for small differences in the amount of sample digested and injected onto the UHPLC column as previously described. Normalization using the top 15 peptides for titin produced similar results, as there was no significant difference between the abundance of myosin heavy chain or titin between the experimental groups. Relative molar abundances for each protein were determined from the average abundance of the top 3 ionizing tryptic peptides that were unique to each protein or specific protein isoform. Relative changes in protein abundances were determined using a pairwise ratio analysis of the abundances of the top 3 and up to the top 15 peptides, which originated from the digestion of each protein or specific protein isoform. The median abundance of each peptide in the sample groups (3 HCM groups) were divided by the median abundance of the peptide in the control group (donor hearts) to determine a pairwise abundance ratio. A ratio or fold-change of 1 was indicative of there being no difference in the protein abundance between the sample and control groups. A grouped protein abundance ratio was determined from the average of all the pairwise ratios (max=15) for each protein or specific protein ratio. Outliers were identified from within the grouped protein abundance ratio if the pairwise ratio for any given peptide was greater than 2 standard deviations from the mean grouped protein abundance ratio. Such outliers could result from a point mutation in the protein, miscleavage of a peptide, and/or post-translational modifications, such as phosphorylation. When an outlier was identified in any group, the peptide was removed from all 4 groups, and the mean grouped protein abundance ratio was recalculated. The removal of the peptide from all 4 groups was performed to avoid biasing the downstream statistics (i.e. t-tests).

Proteins were grouped into subcellular compartments (e.g. mitochondria, sarcomere). Some large protein groups were further divided into subgroups based on the location or function within their group (e.g. inner-membrane of the mitochondria, thick filament of the sarcomere). Large groups of proteins were then compared between samples and controls, to determine for example, if there was a reduction in mitochondrial content in HCM patients. A weighted average of the proteins within each group was created by multiplying the relative molar abundance by the grouped protein ratio for each protein within a group. The weighted average ensures that higher abundant proteins or isoforms have a proportional effect on the average.

**Quantitative targeted LCMS metabolomics.** Metabolites were extracted in cold aqueous/organic solvent mixtures according to validated, optimized protocols in our previously published studies. These protocols use cold conditions and solvents to arrest cellular metabolism and maximize the stability and recovery of metabolites. Each class of metabolites was separated with a unique HPLC method to optimize their chromatographic resolution and sensitivity. Quantitation of metabolites in each assay module was achieved using multiple reaction monitoring of calibration solutions and study samples on an Agilent 1290 Infinity UHPLC/6495 triple quadrupole mass spectrometer. Raw data were processed using Mass

Hunter quantitative analysis software (Agilent). Calibration curves ($R^2$ = 0.99 or greater) were either fitted with a linear or a quadratic curve with a 1/X or $1/X^2$ weighting.

**Supplemental Figures**

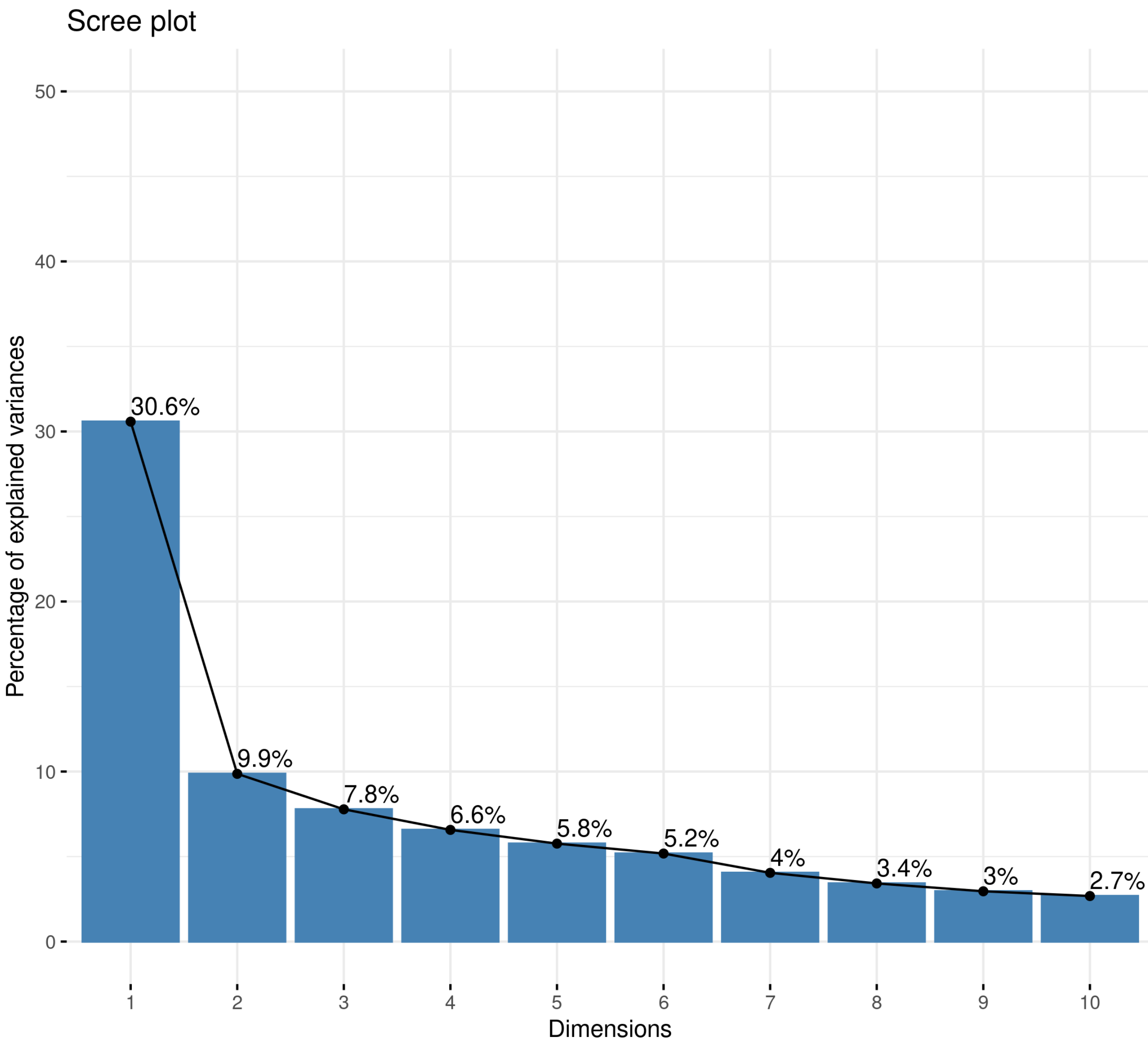


**Figure S1. PCA Scree plot** showing the percentage of explained variances for each dimension.

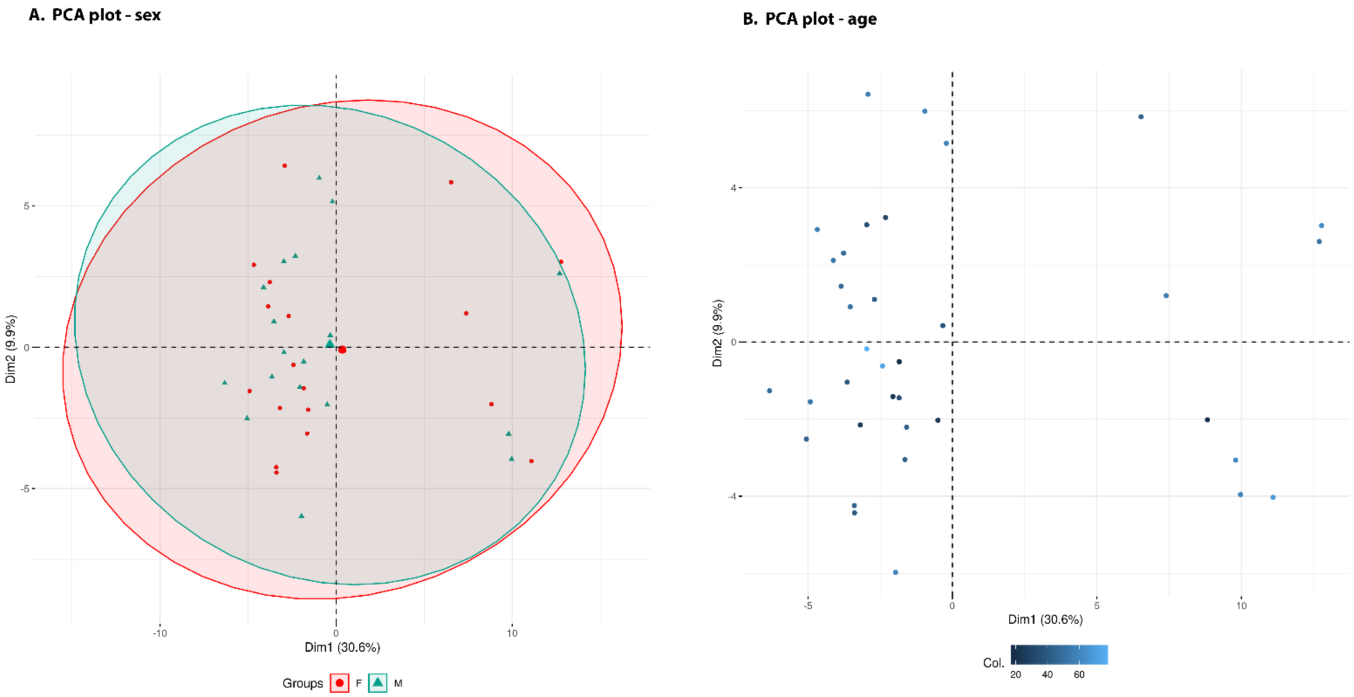


**Figure S2.** **(A)** PCA analysis showing no influence of **(A)** sex or **(B)** age on the abundance of cardiac metabolites.
