## Supplemental Table 2 for "Metabolic and Proteomic Defects in Human Hypertrophic Cardiomyopathy"

| Metabolite | Mean NF | Mean HCM MYBPC | Mean HCM MYH7 | Mean HCM none | Stdev NF | Stdev HCM MYBPC | Stdev HCM MYH7 | Stdev HCM none | overall p.value | overall p.value.adj | adj.p.value HCM MyBPC-NF | adj.p.value HCM MYH7-NF | adj.p.value HCM none-NF | adj.p.value HCM MYH7-MyBPC | adj.p.value HCM none-MyBPC | adj.p.value HCM none-MYH7 |
|---|---|---|---|---|---|---|---|---|---|---|---|---|---|---|---|---|
| **Organic acids** | | | | | | | | | | | | | | | | |
| 3-HBA | 383.6994 | 1340.317257 | 1304.775649 | 1405.636199 | 364.9441 | 803.3544421 | 763.7187181 | 545.7907232 | 0.010762374 | 0.020696874 | 0.017783875 | 0.044031352 | 0.021535576 | 0.999416243 | 0.996424217 | 0.990169013 |
| Citrate | 3807.275 | 5488.704458 | 5242.354991 | 6407.816619 | 1165.89 | 1855.700554 | 1682.402305 | 1602.94343 | 0.026199291 | 0.045171192 | 0.12964227 | 0.311237539 | 0.016052068 | 0.987278564 | 0.61028621 | 0.4913925 |
| Fumarate | 533.961 | 272.9264513 | 279.9865674 | 309.1192069 | 212.165 | 123.8067178 | 90.47296117 | 152.2347789 | 0.002443141 | 0.005964656 | 0.002792312 | 0.008758776 | 0.023436382 | 0.999584365 | 0.94988637 | 0.979043833 |
| Lactate | 48596.84 | 84260.76637 | 82526.49567 | 80299.67557 | 28393.52 | 21978.23233 | 17393.02822 | 32862.8009 | 0.020161733 | 0.036003095 | 0.021080208 | 0.054324651 | 0.079573678 | 0.998780026 | 0.986011757 | 0.998042072 |
| Malate | 3612.457 | 3036.293474 | 2930.203142 | 3458.33886 | 1257.659 | 1394.216367 | 988.8539053 | 1756.479426 | 0.696623435 | 0.716297565 | 0.796991231 | 0.756591371 | 0.995982935 | 0.998262343 | 0.907510795 | 0.869231829 |
| Pyruvate | 242.038 | 323.2972747 | 349.2118929 | 389.6431014 | 92.34623 | 103.5986204 | 155.6781831 | 126.7931204 | 0.111906662 | 0.157615016 | 0.458046913 | 0.297583708 | 0.085593794 | 0.964353667 | 0.623653888 | 0.90597013 |
| Succinate | 3855.074 | 3722.908873 | 3874.025087 | 3116.159185 | 2538.742 | 1510.635762 | 791.9407741 | 1087.956638 | 0.756370354 | 0.764010459 | 0.997888127 | 0.999995201 | 0.79533292 | 0.996855948 | 0.841767337 | 0.782735844 |
| **Amino acids** | | | | | | | | | | | | | | | | |
| 1-Methylhistidine | 104.1176 | 59.08046632 | 61.32583559 | 73.49262123 | 90.86615 | 16.85344981 | 22.08989608 | 31.40238999 | 0.192766009 | 0.253741019 | 0.177651456 | 0.285626949 | 0.570789392 | 0.999585323 | 0.907903996 | 0.954722045 |
| 3-Methylhistidine | 93.96022 | 185.5409771 | 145.2735319 | 138.5280832 | 32.00834 | 107.664967 | 43.6100214 | 66.90726311 | 0.083179486 | 0.125823492 | 0.053374631 | 0.52618444 | 0.637022197 | 0.644511141 | 0.523150353 | 0.997877385 |
| Alanine | 25018.89 | 22180.44185 | 23115.15012 | 20260.3948 | 6318.526 | 8759.66814 | 5429.612173 | 2473.400588 | 0.537792597 | 0.57827161 | 0.778537665 | 0.936973732 | 0.476807203 | 0.989205291 | 0.917454865 | 0.819329132 |
| Arginine | 658.079 | 873.0382532 | 824.9631523 | 843.6441786 | 401.1896 | 208.7130355 | 195.5154024 | 191.6559905 | 0.319528856 | 0.380391496 | 0.280514314 | 0.57363383 | 0.485566984 | 0.976638986 | 0.994430076 | 0.998893821 |
| Asparagine | 3283.041 | 2033.958609 | 2141.441732 | 1690.074946 | 800.4474 | 505.5136101 | 316.2712259 | 477.0538387 | 1.01E-05 | 6.72E-05 | 1.13E-04 | 0.001181678 | 1.05E-05 | 0.972860755 | 0.522902933 | 0.366625289 |
| Aspartate | 6126.907 | 10879.58077 | 9951.044302 | 10819.27398 | 4340.189 | 3809.282841 | 3135.081947 | 3780.364151 | 0.045470037 | 0.07333877 | 0.046087167 | 0.203179913 | 0.083459903 | 0.949501549 | 0.99998447 | 0.967601553 |
| Citrulline | 59.17409 | 155.64782 | 128.2096301 | 159.4678709 | 37.36954 | 74.52496918 | 37.45104757 | 80.40835894 | 0.008248744 | 0.016834172 | 0.010150614 | 0.144723446 | 0.015667697 | 0.773771184 | 0.999133778 | 0.752443844 |
| Glutamate | 36251.79 | 41559.99573 | 42502.1115 | 45835.46199 | 5438.238 | 8639.281776 | 7226.418449 | 13482.30386 | 0.229339587 | 0.290303275 | 0.585594609 | 0.527011385 | 0.174845688 | 0.995837946 | 0.735468262 | 0.88417177 |
| Glutamine | 61641.38 | 77827.61874 | 60395.19794 | 71595.9221 | 18822.43 | 23842.56561 | 12708.68125 | 15273.80755 | 0.164602252 | 0.222435476 | 0.294790426 | 0.999251538 | 0.742240676 | 0.204577809 | 0.888318801 | 0.64273695 |
| Glycine | 3084.873 | 3202.705117 | 3886.047001 | 3611.139353 | 813.772 | 622.9943009 | 1200.771601 | 839.1039348 | 0.218680761 | 0.28035995 | 0.990569468 | 0.268167568 | 0.620705365 | 0.324673766 | 0.730494486 | 0.91965014 |
| Isoleucine | 157.1065 | 422.7590272 | 394.11407 | 472.939052 | 79.87765 | 177.3262094 | 113.2259123 | 104.4611561 | 1.43E-04 | 5.97E-04 | 6.00E-04 | 0.005658173 | 2.06E-04 | 0.963875715 | 0.838130311 | 0.634211904 |
| Leucine | 373.8909 | 784.0680563 | 779.1324713 | 871.1553159 | 184.6267 | 215.6591289 | 268.0928737 | 193.5290912 | 2.58E-04 | 9.55E-04 | 0.001325733 | 0.004001607 | 3.85E-04 | 0.999954959 | 0.816800874 | 0.832232773 |
| Lysine | 1719.121 | 1914.195569 | 1610.967945 | 2006.231305 | 775.9104 | 364.1236438 | 205.3656344 | 447.7045405 | 0.333345778 | 0.383640428 | 0.809782649 | 0.969014227 | 0.633494924 | 0.518106058 | 0.974629905 | 0.367653992 |
| Methionine | 607.8271 | 307.2608938 | 275.717382 | 263.2350363 | 259.9419 | 61.13886459 | 48.4647736 | 49.97683471 | 1.18E-05 | 6.95E-05 | 1.03E-04 | 9.05E-05 | 5.22E-05 | 0.951637487 | 0.881420529 | 0.997486358 |
| Ornithine | 316.8468 | 412.8426289 | 411.9002867 | 423.1366549 | 166.2133 | 89.65856069 | 156.7229845 | 165.7301171 | 0.400635793 | 0.445150882 | 0.460821677 | 0.546293341 | 0.451536564 | 0.999998873 | 0.998545292 | 0.998560678 |
| Phenylalanine | 427.5886 | 400.2536144 | 363.1428753 | 420.1944366 | 123.6218 | 115.7791547 | 68.48696678 | 57.75089175 | 0.566869789 | 0.603052967 | 0.928833186 | 0.56455906 | 0.998768517 | 0.842087477 | 0.970410078 | 0.656938453 |
| Proline | 814.0535 | 998.3481352 | 942.1007514 | 942.2420736 | 215.8113 | 289.9668352 | 379.1800065 | 285.7336256 | 0.604068555 | 0.635861637 | 0.534047299 | 0.824537473 | 0.824064343 | 0.975596149 | 0.975771235 | 1 |
| Serine | 3882.237 | 2513.939165 | 3024.766436 | 2035.942285 | 1443.109 | 507.5361853 | 1172.785773 | 535.6301224 | 0.003165473 | 0.007361565 | 0.017776558 | 0.291981658 | 0.002660225 | 0.646850063 | 0.692591589 | 0.182750462 |
| Threonine | 3288.717 | 2262.26228 | 2335.909177 | 2259.530491 | 1515.618 | 503.0914556 | 755.4970228 | 789.3785151 | 0.078785902 | 0.121209079 | 0.090018388 | 0.185950817 | 0.136359479 | 0.998056756 | 0.9999999 | 0.998348905 |
| Tyrosine | 416.6377 | 566.2342771 | 469.8573287 | 529.0787648 | 133.114 | 151.0757213 | 124.5200528 | 87.63206151 | 0.084301739 | 0.125823492 | 0.074296782 | 0.843935802 | 0.322730716 | 0.377081248 | 0.922231753 | 0.797556974 |
| Valine | 349.2671 | 1022.106208 | 1040.303036 | 1065.908555 | 169.3578 | 316.6319485 | 331.9277062 | 223.249001 | 9.52E-06 | 6.72E-05 | 4.02E-05 | 1.06E-04 | 6.18E-05 | 0.99889236 | 0.985170352 | 0.997665367 |
| **Acyl carnitines** | | | | | | | | | | | | | | | | |
| C02 | 904.6053 | 193.1394148 | 157.2739172 | 222.1342477 | 1816.27 | 91.97788043 | 38.54983514 | 77.477 | 0.243506378 | 0.300625158 | 0.278721085 | 0.31366995 | 0.392004871 | 0.999712855 | 0.999848101 | 0.998715816 |
| C03 | 31.73248 | 9.133992813 | 7.660826948 | 10.1548541 | 21.16805 | 4.458373029 | 2.012074576 | 5.681894618 | 1.16E-04 | 5.28E-04 | 2.96E-04 | 4.27E-04 | 0.001591647 | 0.990067203 | 0.996629459 | 0.965180672 |
| C03-DC | 4.651758 | 2.360599272 | 2.337953973 | 2.839818704 | 1.860217 | 1.021097738 | 0.797556126 | 1.273557684 | 0.001756573 | 0.004622561 | 0.002132069 | 0.004905629 | 0.035676257 | 0.99997803 | 0.840959062 | 0.857891234 |
| C04 Butyryl | 90.54529 | 49.74996191 | 60.82647898 | 63.37411775 | 35.41957 | 18.21689042 | 10.49557614 | 43.9977992 | 0.034453879 | 0.057423132 | 0.020368102 | 0.189510006 | 0.256846581 | 0.835361649 | 0.731963967 | 0.998014355 |
| C04 Isobutyryl | 16.98745 | 1.357823002 | 1.13181144 | 1.098103265 | 10.63792 | 1.491950366 | 0.716583469 | 0.750679764 | 1.60E-07 | 2.00E-06 | 6.42E-07 | 3.06E-06 | 2.94E-06 | 0.999957494 | 0.999906963 | 0.999999138 |
| C04:1 | 0.331054 | 1.083251704 | 1.382332549 | 3.025090659 | 0.131327 | 0.950308951 | 1.311551986 | 2.648810567 | 0.006111171 | 0.013002491 | 0.615550174 | 0.577023246 | 0.004039316 | 0.997691646 | 0.036205085 | 0.091061359 |
| C04-DC MeMalonyl | 2.294486 | 1.328502995 | 1.48645577 | 1.401931116 | 1.132711 | 0.483191577 | 0.513110472 | 0.402321978 | 0.019688599 | 0.035797452 | 0.018006929 | 0.097710323 | 0.057144922 | 0.955231732 | 0.99511198 | 0.99434187 |
| C04-DC Succinyl | 8.625231 | 4.844185612 | 5.333430581 | 5.09037861 | 4.607108 | 1.446246536 | 1.424927762 | 2.662397156 | 0.022763214 | 0.039935463 | 0.022288391 | 0.092100854 | 0.062833287 | 0.978712849 | 0.997156164 | 0.997912048 |
| C04-OH Butyryl | 4.96678 | 8.956382068 | 12.17418586 | 14.75386248 | 4.452462 | 2.547112354 | 7.207211101 | 8.706043861 | 0.013042167 | 0.024607863 | 0.456326571 | 0.08714851 | 0.011157355 | 0.631257349 | 0.15574447 | 0.816030654 |
| C04-OH Isobutyryl | 69.09157 | 276.5257137 | 280.1640551 | 207.2106138 | 44.30759 | 87.32232226 | 99.82955856 | 71.45468588 | 0.001266634 | 0.003518427 | 0.004743632 | 0.015938927 | 0.923129172 | 0.998351664 | 0.026952327 | 0.068778487 |
| C05 2-Methylbutyryl | 15.23 | 3.525719017 | 2.509558162 | 3.898884397 | 9.237264 | 1.942635349 | 0.437376108 | 1.430615621 | 3.96E-06 | 3.30E-05 | 1.63E-05 | 1.85E-05 | 1.09E-04 | 0.960248351 | 0.997857478 | 0.926736971 |
| C05 Isovaleryl | 29.59436 | 2.055938887 | 0.882720567 | 2.423602012 | 23.16457 | 3.395944389 | 0.941528074 | 2.387137191 | 9.80E-06 | 6.72E-05 | 3.12E-05 | 6.65E-05 | 1.48E-04 | 0.995512543 | 0.999859127 | 0.99233983 |
| C05 Valeryl | 1.169189 | 0.352445056 | 0.324053481 | 0.299137121 | 1.053027 | 0.193748184 | 0.13064893 | 0.235487752 | 0.003980151 | 0.00884478 | 0.008562715 | 0.013908429 | 0.010917324 | 0.999377821 | 0.995941815 | 0.999679419 |
| C05:1 | 3.320352 | 2.087862652 | 1.717771592 | 1.615859896 | 1.412602 | 0.895247706 | 0.587195346 | 0.435257945 | 0.002445509 | 0.005964656 | 0.027564864 | 0.00686036 | 0.003791719 | 0.809727612 | 0.670498683 | 0.995959793 |
| C05-DC | 0.961555 | 0.630345805 | 0.433363129 | 0.697209496 | 0.581088 | 0.350579919 | 0.177485442 | 0.479653562 | 0.105454644 | 0.150649491 | 0.319012373 | 0.072996226 | 0.588214907 | 0.72948148 | 0.984777004 | 0.589696082 |
| C05-OH | 15.62572 | 12.02142063 | 8.456420559 | 8.734560134 | 7.393037 | 5.154679668 | 2.406445923 | 2.032450949 | 0.017273153 | 0.03198732 | 0.372849585 | 0.027100563 | 0.035587586 | 0.38240382 | 0.453300228 | 0.999439128 |
| C06 | 7.932216 | 4.853104531 | 6.128154494 | 5.864939557 | 3.878965 | 1.461142432 | 2.104085718 | 2.869295422 | 0.101541684 | 0.147161862 | 0.06542581 | 0.518112779 | 0.400559061 | 0.708939997 | 0.829916337 | 0.997022863 |
| C06-OH | 1.413365 | 2.103746602 | 2.353000682 | 2.95054932 | 0.739764 | 0.89048541 | 0.830452883 | 1.850878146 | 0.075878669 | 0.118560421 | 0.550935203 | 0.364573218 | 0.050586973 | 0.962901943 | 0.375762222 | 0.72098321 |
| C08 | 2.884578 | 1.845806503 | 1.901724297 | 1.885120861 | 0.917482 | 0.714051448 | 0.589123927 | 1.033554775 | 0.037117528 | 0.060848407 | 0.04235108 | 0.09678399 | 0.088972847 | 0.998776354 | 0.999572756 | 0.999975423 |
| C08-OH | 0.782169 | 0.662439439 | 0.704358629 | 0.864186789 | 0.327707 | 0.297769763 | 0.200686806 | 0.533277732 | 0.628835815 | 0.655037308 | 0.879373072 | 0.97102412 | 0.966356277 | 0.993732598 | 0.600347936 | 0.802707887 |
| C10 | 2.747855 | 1.20315547 | 1.389122735 | 1.191434893 | 0.82115 | 0.450535108 | 0.392765579 | 0.52072146 | 2.59E-06 | 2.35E-05 | 5.06E-06 | 1.60E-04 | 2.06E-05 | 0.883767513 | 0.999964037 | 0.892498076 |
| C10-OH | 0.580265 | 0.271316939 | 0.283496697 | 0.310719328 | 0.269319 | 0.130356892 | 0.146591418 | 0.226925161 | 0.006934839 | 0.014447581 | 0.007480843 | 0.022082217 | 0.042838333 | 0.99905236 | 0.970158186 | 0.992144803 |
| C12 | 3.274682 | 0.1275 | 0.165 | 0.07125 | 1.485157 | 0.239055908 | 0.26043919 | 0.201525433 | 7.87E-11 | 3.94E-09 | 4.54E-10 | 4.97E-09 | 2.53E-09 | 0.999468332 | 0.998217361 | 0.993818251 |
| C14 | 17.61811 | 3.386026974 | 3.804871752 | 2.723112289 | 6.89883 | 2.736196683 | 1.181182616 | 0.967787458 | 1.11E-09 | 3.70E-08 | 6.28E-09 | 8.55E-08 | 1.72E-08 | 0.99441565 | 0.978685767 | 0.934693727 |
| C14:1 | 5.704254 | 1.390249764 | 1.339788105 | 1.285017038 | 2.799619 | 2.443743841 | 1.313764426 | 1.779381853 | 3.59E-04 | 0.001237757 | 8.30E-04 | 0.002064256 | 0.001800627 | 0.999953373 | 0.999578595 | 0.999954643 |
| C14:1-OH | 0.558714 | 0.183041996 | 0.254277399 | 0.140741592 | 0.343061 | 0.1727476 | 0.192248225 | 0.091313587 | 0.001490746 | 0.004029042 | 0.002921087 | 0.036945272 | 0.002480999 | 0.884999531 | 0.972437175 | 0.715971139 |
| C14:2 | 2.039541 | 0.779045916 | 0.555638164 | 0.5026064 | 1.366386 | 1.950109225 | 0.638092362 | 0.955794854 | 0.118351474 | 0.164377047 | 0.22790758 | 0.176062272 | 0.153148681 | 0.985536923 | 0.973291012 | 0.99984585 |
| C14-OH | 2.012044 | 0.691291546 | 0.893751432 | 0.566852303 | 0.685021 | 0.364447359 | 0.731005408 | 0.285631064 | 1.11E-05 | 6.92E-05 | 3.17E-05 | 0.001077786 | 3.23E-05 | 0.838032088 | 0.955520998 | 0.613511614 |
| C16 | 85.7375 | 12.27988735 | 13.85562269 | 10.31395886 | 53.36792 | 7.998311306 | 3.159672469 | 3.768390511 | 7.46E-07 | 8.29E-06 | 2.57E-06 | 1.79E-05 | 8.01E-06 | 0.999095005 | 0.9982498 | 0.992368215 |
| C16:1 | 32.97157 | 9.954022378 | 12.97963461 | 12.47683692 | 14.24195 | 7.682624521 | 9.238358341 | 9.568760466 | 1.34E-04 | 5.80E-04 | 1.26E-04 | 0.002276497 | 0.001735911 | 0.914062587 | 0.947576324 | 0.999645765 |
| C16:1-OH | 3.580813 | 0.955356245 | 1.352626646 | 0.675388298 | 1.683907 | 0.570912855 | 0.905035047 | 0.303494101 | 1.96E-06 | 1.96E-05 | 7.06E-06 | 3.37E-04 | 5.92E-06 | 0.803966653 | 0.919772879 | 0.506566111 |
| C16:2 | 4.924908 | 1.360746609 | 1.750932663 | 0.815748851 | 2.065993 | 2.001859381 | 0.914268944 | 0.729811354 | 4.66E-05 | 2.22E-04 | 1.82E-04 | 0.002302048 | 8.77E-05 | 0.951383774 | 0.880612304 | 0.65728011 |
| C16:2-OH | 0.590649 | 0.156406859 | 0.300307255 | 0.112950223 | 0.398783 | 0.147118408 | 0.288323833 | 0.088067511 | 6.51E-04 | 0.00203344 | 0.002252842 | 0.111705772 | 8.43E-04 | 0.51988741 | 0.869606075 | 0.224571688 |
| C16-OH | 6.219388 | 1.706683568 | 2.273119408 | 1.455416445 | 1.737728 | 0.973337657 | 1.521066128 | 0.947103743 | 1.93E-08 | 3.85E-07 | 6.71E-08 | 5.08E-06 | 1.44E-07 | 0.77551512 | 0.974020144 | 0.595215621 |
| C18 | 33.09037 | 10.66204332 | 11.36863857 | 8.586925325 | 13.46255 | 4.291493405 | 2.77123293 | 3.675597296 | 7.94E-08 | 1.13E-06 | 4.73E-07 | 4.62E-06 | 4.98E-07 | 0.996285895 | 0.918559463 | 0.862032797 |
| C18:1 | 263.9745 | 81.41727592 | 80.23459035 | 116.9909547 | 162.7415 | 63.3997669 | 20.61420679 | 80.21617693 | 0.002183332 | 0.005598286 | 0.001815804 | 0.004473535 | 0.023152602 | 0.99999045 | 0.790480931 | 0.83249221 |
| C18:1-OH | 18.79395 | 4.818308245 | 6.949693412 | 4.001832999 | 6.394893 | 2.780135726 | 3.971132005 | 1.857571994 | 1.56E-08 | 3.85E-07 | 5.20E-08 | 7.08E-06 | 1.06E-07 | 0.646943987 | 0.969164306 | 0.459258216 |
| C18:2 | 151.8584 | 86.85745951 | 93.52001341 | 104.8288048 | 106.8839 | 73.22763611 | 22.82490254 | 68.79248063 | 0.287556806 | 0.346453983 | 0.25121397 | 0.450038736 | 0.599960618 | 0.997653999 | 0.95271767 | 0.991276591 |
| C18:2-OH | 6.473278 | 1.749879256 | 2.762914083 | 1.361927577 | 3.71087 | 1.076093642 | 1.685310224 | 0.720699666 | 3.97E-05 | 1.98E-04 | 8.86E-05 | 0.004945861 | 1.06E-04 | 0.698004957 | 0.975088031 | 0.522957758 |
| C18-OH | 3.074087 | 1.822072036 | 2.320962058 | 1.537380801 | 0.847323 | 0.904604848 | 1.499047278 | 1.01172173 | 0.033374265 | 0.056566551 | 0.070482758 | 0.507187111 | 0.035294984 | 0.740462947 | 0.937171036 | 0.473318892 |
| C20 | 0.522848 | 0.688870331 | 0.561360291 | 0.420172394 | 0.357691 | 0.566337625 | 0.220564235 | 0.145817333 | 0.505709862 | 0.549684632 | 0.789966744 | 0.997252094 | 0.952651502 | 0.891219954 | 0.447922298 | 0.888137195 |
| C20:1 | 7.461472 | 2.562070048 | 3.330163719 | 1.563329967 | 4.147564 | 2.452581207 | 1.151440204 | 0.799057671 | 2.06E-04 | 7.92E-04 | 8.47E-04 | 0.012033676 | 2.55E-04 | 0.906676788 | 0.817923197 | 0.501144672 |
| C20:1-OH | 2.548005 | 1.197171712 | 1.441780834 | 0.950949776 | 1.237119 | 0.771275311 | 0.594648126 | 0.660431173 | 0.002848249 | 0.006781546 | 0.007101383 | 0.060739756 | 0.003397088 | 0.919902921 | 0.918478472 | 0.653766199 |
| C20:2 | 6.804897 | 1.96143475 | 3.410387904 | 1.181132881 | 4.395261 | 1.981906341 | 1.674069534 | 0.47649664 | 3.17E-04 | 0.001133681 | 9.52E-04 | 0.049031956 | 4.65E-04 | 0.587567036 | 0.902261819 | 0.298939255 |
| C20:2-OH | 1.981666 | 0.903753155 | 1.480745898 | 0.771433355 | 1.079869 | 0.435944899 | 0.924778716 | 0.593696394 | 0.009155854 | 0.018311708 | 0.020487899 | 0.563303188 | 0.016949359 | 0.364386146 | 0.981105675 | 0.267150648 |
| C20:3 | 2.558373 | 1.351919114 | 2.387193506 | 1.613188468 | 1.455525 | 0.45130694 | 2.566845782 | 0.93936344 | 0.238459657 | 0.298074571 | 0.2941649 | 0.99548036 | 0.579548091 | 0.426065884 | 0.979690388 | 0.720859627 |
| C20:4 | 2.679885 | 7.92482057 | 13.13125968 | 10.75223104 | 1.756419 | 3.112503934 | 15.01237887 | 5.419254698 | 0.060891834 | 0.096653705 | 0.456951063 | 0.050533866 | 0.178387427 | 0.463332631 | 0.853186595 | 0.926338914 |
| C20-OH/C22:6 | 0.777302 | 0.738559687 | 0.974632764 | 1.43703053 | 0.551397 | 0.273338377 | 0.64094759 | 1.631860437 | 0.337603577 | 0.383640428 | 0.999665125 | 0.968886032 | 0.444113411 | 0.933919428 | 0.315554967 | 0.717211605 |
| C22:4 | 1.13841 | 0.458706311 | 0.607087473 | 0.487974752 | 0.837561 | 0.10665995 | 0.342746047 | 0.179663575 | 0.009514319 | 0.018655527 | 0.009015868 | 0.09024336 | 0.026434478 | 0.877863867 | 0.998841399 | 0.946859096 |
| C22:5 | 0.847247 | 0.623245296 | 0.851194514 | 0.819730798 | 0.692908 | 0.147820611 | 0.745646163 | 0.37944085 | 0.701971614 | 0.716297565 | 0.777076462 | 0.999998679 | 0.999554739 | 0.76785447 | 0.837327861 | 0.999335236 |
| **Nucleotides** | | | | | | | | | | | | | | | | |
| ADP | 5736.439 | 6534.78252 | 6071.034883 | 7143.086921 | 1040.791 | 1458.307868 | 828.6435006 | 866.9453274 | 0.091522452 | 0.134591842 | 0.423871823 | 0.934106734 | 0.081769717 | 0.806361911 | 0.645557719 | 0.251042424 |
| ADPR | 677.5151 | 476.8680758 | 439.4164313 | 443.6910694 | 145.3217 | 130.747382 | 77.66699768 | 91.86926436 | 5.60E-04 | 0.001805459 | 0.003626688 | 0.001544913 | 0.001889528 | 0.89553274 | 0.924311415 | 0.9998565 |
| AMP | 2121.268 | 2571.604272 | 2351.649502 | 2717.209744 | 630.0838 | 934.5049615 | 332.3280434 | 1050.926084 | 0.470411183 | 0.516935366 | 0.61828786 | 0.940201031 | 0.463777723 | 0.932407424 | 0.978768263 | 0.802322176 |
| ATP | 23402.31 | 12194.44773 | 13381.4622 | 14358.10506 | 3248.19 | 3244.992726 | 1847.953897 | 4132.040446 | 7.52E-08 | 1.13E-06 | 6.67E-08 | 3.41E-06 | 1.95E-05 | 0.851098676 | 0.467263777 | 0.929609708 |
| CDP | 54.16262 | 66.18859245 | 77.54480574 | 63.3678795 | 8.178938 | 27.46838006 | 29.10699848 | 18.85516359 | 0.265311223 | 0.323550271 | 0.669872921 | 0.202714113 | 0.856444636 | 0.707823471 | 0.993230992 | 0.616867792 |
| CMP | 70.3876 | 61.84518142 | 69.92264664 | 61.40425101 | 13.71079 | 10.51654169 | 14.53724064 | 15.81191279 | 0.329700376 | 0.383640428 | 0.511727689 | 0.999877926 | 0.54571206 | 0.55805593 | 0.999863164 | 0.588368174 |
| CTP | 300.7831 | 152.3097417 | 188.1753577 | 151.8052 | 74.77088 | 79.97508944 | 61.13198599 | 87.52796581 | 8.43E-04 | 0.002480059 | 0.001003536 | 0.030141237 | 0.002671458 | 0.738083352 | 0.999998909 | 0.780540884 |
| GDP | 316.4989 | 447.0388059 | 485.689667 | 482.2493423 | 55.9247 | 81.55777482 | 64.57552614 | 87.9351863 | 1.65E-04 | 6.60E-04 | 0.00300611 | 4.28E-04 | 5.56E-04 | 0.671509695 | 0.73143473 | 0.999713344 |
| GMP | 99.64168 | 141.1837404 | 130.2017371 | 143.3266685 | 32.85922 | 38.00527563 | 25.58818036 | 76.84991481 | 0.212693495 | 0.276225318 | 0.226258033 | 0.561815303 | 0.258078949 | 0.95446921 | 0.999624761 | 0.942052884 |
| GTP | 405.9272 | 320.6684834 | 383.4467915 | 385.4822863 | 68.25945 | 86.92106984 | 86.72401574 | 122.1802764 | 0.192843174 | 0.253741019 | 0.200936705 | 0.961458416 | 0.970529201 | 0.455427555 | 0.427641902 | 0.999968535 |
| IMP | 66.74993 | 274.7828805 | 188.792764 | 300.9322532 | 24.39603 | 403.572969 | 53.40887084 | 402.666039 | 0.394971374 | 0.443788061 | 0.448756343 | 0.852131311 | 0.425009111 | 0.924790356 | 0.997567609 | 0.880784127 |
| ITP | 27.42395 | 10.86070542 | 8.355881625 | 11.22930943 | 9.957813 | 9.348013849 | 1.767021644 | 2.375976563 | 2.53E-05 | 0.00014055 | 1.28E-04 | 6.20E-05 | 5.85E-04 | 0.876305153 | 0.999509848 | 0.860868444 |

| | | | | | | | | | | | | | | | | |
|---|---|---|---|---|---|---|---|---|---|---|---|---|---|---|---|---|
| UDP | 2520.368 | 2242.709195 | 2141.894476 | 2140.490299 | 386.9019 | 373.0372364 | 338.4300079 | 320.3526641 | 0.33482082 | 0.383640428 | 0.440387664 | 0.333768548 | 0.847903296 | 0.983589729 | 0.923300832 | 0.804803119 |
| UMP | 141.2703 | 188.3252184 | 161.0476161 | 186.8945579 | 21.99509 | 41.97066087 | 19.0142319 | 10.66839336 | 0.980298424 | 0.980298424 | 0.999994082 | 0.999982224 | 0.98666049 | 0.999898432 | 0.985856816 | 0.982024562 |
| UTP | 381.2643 | 639.9155967 | 550.6786588 | 646.5475492 | 104.8672 | 150.106749 | 65.0853739 | 196.300979 | 0.004583082 | 0.009963223 | 0.004581939 | 0.132354749 | 0.012976411 | 0.623956796 | 0.999744305 | 0.735969925 |
| **NAD and acetyl CoA** | | | | | | | | | | | | | | | | |
| NAD | 4.691106 | 12.3141794 | 11.79886416 | 13.01712482 | 2.268056 | 3.635992588 | 3.833278077 | 3.24445263 | 0.132522684 | 0.181537923 | 0.340460043 | 0.17005649 | 0.167605927 | 0.925851293 | 0.923032757 | 0.999999822 |
| NADP | 199.9719 | 249.1546881 | 260.8739814 | 228.8726125 | 80.69252 | 70.31612506 | 66.53985054 | 66.35787664 | 0.003628957 | 0.008247629 | 0.005416782 | 0.517474995 | 0.015734281 | 0.17710261 | 0.999516278 | 0.287213674 |
| NAM | 210.8816 | 210.1928638 | 211.9701997 | 200.8376425 | 65.44582 | 60.05838118 | 55.7045496 | 57.13112113 | 0.001227043 | 0.003505838 | 0.001618132 | 0.092351106 | 0.003275739 | 0.508662744 | 0.999588044 | 0.525023876 |
| NMN | 841.631 | 530.1764085 | 632.9293953 | 537.6971461 | 90.74577 | 189.491955 | 140.2830998 | 270.9081012 | 3.74E-05 | 1.97E-04 | 1.15E-04 | 9.47E-04 | 1.20E-04 | 0.986506083 | 0.967073788 | 0.88495605 |
| NADH | 671.19 | 357.4554968 | 224.7362281 | 349.0073931 | 277.0276 | 190.7259987 | 77.27970202 | 163.8656536 | 4.04E-04 | 0.00134531 | 0.005580148 | 2.85E-04 | 0.01002297 | 0.436312852 | 0.99966673 | 0.568766741 |
| NADPH | 62.10633 | 3.540607896 | 2.825483862 | 14.32107885 | 18.54873 | 3.144214107 | 1.778700864 | 4.466867971 | 7.42E-15 | 7.42E-13 | 1.35E-13 | 4.20E-13 | 5.13E-12 | 0.999691545 | 0.574790776 | 0.591032998 |
| Acetyl CoA | 17.1996 | 3.19754622 | 2.16691901 | 5.535998101 | 14.65081 | 2.297166964 | 1.168762596 | 5.394320357 | 7.69E-04 | 0.002329498 | 0.001359561 | 0.001705774 | 0.018369106 | 0.990125696 | 0.900866333 | 0.80225519 |

Cells highlighted in pink represent significant adjusted p values in one or more between groups comparisons.
